# Human neurogenesis is altered via glucocorticoid-mediated regulation of *ZBTB16* expression

**DOI:** 10.1101/2022.08.21.504700

**Authors:** Anthi C. Krontira, Cristiana Cruceanu, Leander Dony, Christina Kyrousi, Marie-Helen Link, Nils Kappelmann, Dorothee Pöhlchen, Catarina Raimundo, Signe-Penner Goeke, Alicia Schowe, Darina Czamara, Marius Lahti-Pulkkinen, Sara Sammallahti, Elina Wolford, Kati Heinonen, Simone Roeh, Vincenza Sportelli, Barbara Wölfel, Maik Ködel, Susann Sauer, Monika Rex-Haffner, Katri Räikkönen, Marta Labeur, Silvia Cappello, Elisabeth B. Binder

## Abstract

Glucocorticoids are important for proper organ maturation and their levels are tightly regulated during development. Here we use human cerebral organoids and mice to study cell-type specific effects of glucocorticoids on neurogenesis. We show that glucocorticoids increase a specific type of basal progenitors (co-expressing *PAX6* and *EOMES*) that has been shown to drive cortical expansion in gyrified species. This effect is mediated via the transcription factor *ZBTB16* and leads to increased production of neurons. A phenome-wide mendelian randomization analysis of an enhancer variant that moderates glucocorticoid-induced *ZBTB16* levels, reveals causal relationships with higher educational attainment and altered brain structure. The relationship with postnatal cognition is supported by data from a prospective pregnancy cohort. This study provides a novel cellular and molecular pathway for the effects of glucocorticoids on human neurogenesis that relates to lasting postnatal phenotypes.

## Introduction

Prenatal development relates to postnatal health. The ‘developmental origin of health and disease (DOHaD) hypothesis’^1^ proposes that environmental exposures during critical prenatal periods have lasting effects on cells and tissues impacting human health throughout life. This is also true for the central nervous system (CNS) with a number of environmental factors shown to impact its cellular architecture and function^2^.

An important factor for CNS development during pregnancy are glucocorticoids (GCs)^3^. These steroid hormones, the main one being cortisol in humans, are endogenously present prenatally. They regulate fetal organ development and maturation, being especially important for lung and brain^4^. GC levels are very tightly regulated during gestation in a species-specific pattern. While there is a physiological rise of GCs close to term in most species that is important for final organ development and parturition, their levels throughout gestation differ. For example, corticosterone, the main GC in mice, exhibits a sharp increase late in gestation, between embryonic day 16 and 17, while in humans cortisol levels increase progressively during gestation starting at the beginning of the second trimester^5^.

GCs are found increased outside the physiological range and/or time-window due to the therapeutic use of synthetic GCs (sGCs) during pregnancy but also as a result of endocrine and stress-related disorders of the mother^3^. The placenta constitutes a protective barrier for endogenous maternal GCs, with only as little as 10% of the maternal circulating hormones reaching the fetal compartments. This barrier has been shown to be affected by maternal stress and depression resulting in reduced placental cortisol metabolism^6^. Furthermore, sGCs readily cross the placenta^7^ leading to higher exposure of the fetus. sGCs, either betamethasone or dexamethasone, are most commonly prescribed starting at 22 and until 33 gestational weeks (GWs) in pregnancies with high risk for preterm delivery to facilitate fetal lung maturation^8^. More than 1 in 10 babies are born prematurely every year, a number which amounts to ∼15 million preterm births (< GW37) per year^9^ of which ∼ 615,000 are born extremely preterm (< GW28)^10^, highlighting the clinical and societal importance of prenatal sGCs use. sGC treatment has also been associated with a 3.5-fold increase of survival rates in preterm infants born at GW22^11^, highlighting the importance of an even more extensive use of sGCs in extremely preterm births. Given the clear anti-inflammatory effects of sGCs, they were also used during the COVID-19 pandemic in patients requiring oxygen therapy and mechanical ventilation^12^. Pregnant women were not excluded from this treatment course, with the guidelines specifically promoting the use of dexamethasone in women that fulfill the afore-mentioned criteria^13^.

Exposure to levels of prenatal GCs outside of the physiological range has been related to lasting postnatal effects at the behavioral and brain structural levels^14^. While the effects of steroids on brain structure and function may also be mediated by inflammatory agents, direct effects are supported by data from animal models. In fact, the overall molecular and cellular effects of GCs on the term and adult brain are well-characterized in rodents^15^. Interestingly, their impact on early stages of brain development and especially during the neurogenic period, which extends until GW28^16^ in humans and thus in the timeframe of sGC administration for extremely preterm births, have rarely been studied and are absent for the human brain or complex models of the human developing cortex. To address this important question, we combined experiments in human cerebral organoids (hCOs) and mouse embryos with human genetic analyses and mechanistically linked enhanced prenatal sGC exposure to human cortical neurogenesis and possibly to lasting effects on mental and cognitive abilities and brain structure.

## Results

### Glucocorticoids increase the number of basal progenitors

To study GC regulation of neurogenic trajectories in the developing neocortex we treated hCOs^17^ for 7 days with 100nM of dexamethasone (dex), a dose and time consistent with therapeutic guidelines followed in clinical settings^8^ (see Methods for detailed explanation). This treatment was initiated at day 43 (Figure 1A) in hCOS derived from 2 independent iPSC (induced pluripotent stem cell) lines. Days 40-50 were chosen as a time-range when hCOs are actively performing neurogenic processes with all the progenitor cell types present while birth of deep layer neurons is peaking and of upper layer neurons has started^18^. First, we analyzed the specific effects of dex on different progenitor cell types defined by the expression of PAX6 (Paired Box 6) and EOMES (Eomesodermin – also known as T-box brain protein 2 or TBR2). PAX6 is highly expressed in radial glia (RG) cells, EOMES, but not PAX6, in intermediate basal progenitors (IPs), while both can be expressed in certain basal progenitors (BPs). Dex consistently led to a significant increase of PAX6+EOMES+ BPs (Figure 1B,C) in hCOs derived from both iPSC lines compared to veh-treated hCOs. The increased PAX6+EOMES+ BPs were localized at the basal side of the germinal zones, in the subventricular-like zone (SVZ) (Figure 1D and Figure S1A,B). Moreover, we confirmed the effects of dex on the increase of double-positive BPs by analyzing the number of PAX6+EOMES-, PAX6-EOMES+ and PAX6+EOMES+ progenitor subtypes in No.1– hCOs with flow cytometry analysis (FCa). We observed a significant increase (+11%) in PAX6+EOMES+ BPs when hCOs were treated with dex compared to veh (Figure S2A-C). Co-administration of the glucocorticoid receptor (GR) antagonist RU486 supported that dex effects are mainly mediated by the GR and not the mineralocorticoid receptor (Figure S2A,D,E). To study whether dex treatment affected the transcriptomic profile of PAX6+EOMES+ BPs we generated and analyzed single-cell RNA sequencing (sc-RNA seq) data of HPS0076 hCOs treated with 100nM of dex for 10 days (treatment initiation at day 60). We found a 3.38-fold increase in the numbers of the PAX6+EOMES+ cells after dex, similar to the fold changes we found with our immunostaining analysis (Figure S1C). Interestingly, gene set enrichment analysis of the differentially expressed (DE) genes between treatment groups in these double positive cells pinpointed gene sets associated with cell fate commitment and glia cell proliferation and differentiation (Figure S1D & Tables S2,3), highlighting the importance of these cells in regulating neurogenic processes (see Supplemental Results for more details).

**Figure 1.**
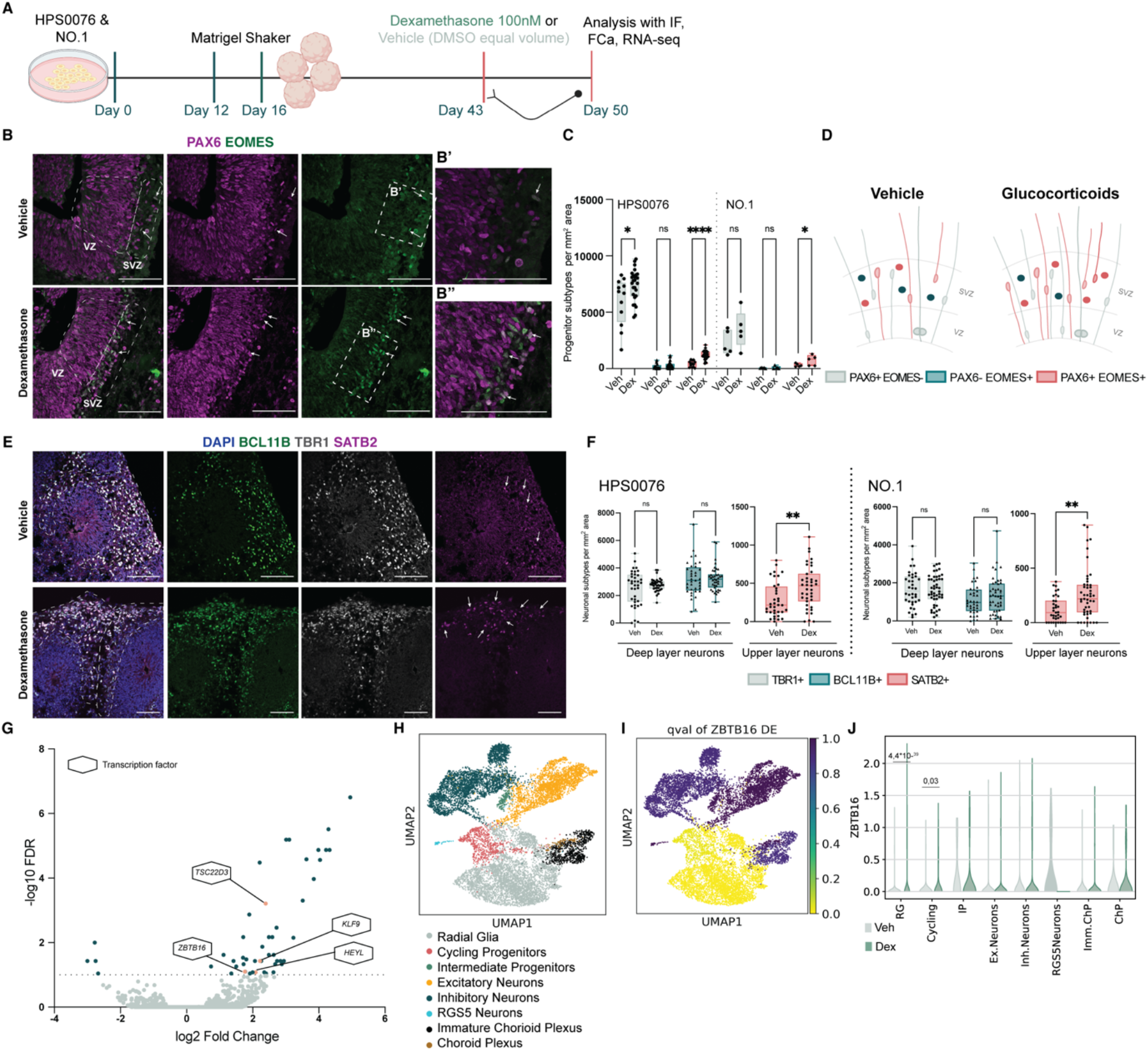
| Glucocorticoids increase basal progenitors that co-express PAX6 and EOMES. **A**, Treatment and analysis workflow in human cerebral organoids (hCOs) derived from two iPSC lines. **B,** Representative images of day 50 hCOs at veh and dex conditions stained for PAX6 and EOMES. Arrows indicate cells that co-express PAX6 and EOMES; Scale bars, 100μm. **B’ and B’’,** Zoom-ins of the areas shown in vehicle and dex ventricles respectively in Figure B. Scale bars, 100μm. **C,** Quantification of the progenitor subtypes in each treatment condition normalized by mm^2^ of quantified total area in hCOs produced from two iPSC lines. **D**, Schematic representation of the effects of dex on progenitors, highlighting the increased numbers of basal progenitors co-expressing PAX6 and EOMES. **E,** Representative images of day 50 hCOs at vehicle and dex conditions stained for layer VI neurons with TBR1, layer V neurons with BCL11B, layer IV neurons with SATB2 and DAPI. Arrows indicate SATB2 positive cells in the CP; Scale bars, 100μm. **F,** Quantification of TBR1, BCL11B and SATB2 cells found in the CP normalized per mm^2^ of area. **G,** Volcano plot of DE gene expression analysis in bulk RNA sequencing. Grey dots, genes with non-significant expression changes at an FDR cutoff of 10%; Blue dots, genes with significant expression changes; Orange dots denote TFs labeled with their gene name. **H,** Single-cell clusters of HPS0076 hCOs treated with dex 100nM for 10 days starting at day 60. **I,** UMAP plot showing *ZBTB16* FDR-value of dex response per cluster. **J,** Violin plot of *ZBTB16* expression changes per cluster. FDR, false discovery rate with Benjamini-Hochberg correction. DMSO, dimethyl sulfoxide; IF, Immunofluorescence; Seq, sequencing; Veh, vehicle; Dex, dexamethasone; VZ, ventricular-like zone; SVZ, subventricular-like zone; BPs, basal progenitors; CP, cortical plate; TFs, transcription factors; RG, Radial Glia; Cycling, Cycling Progenitors; IP, Intermediate Progenitors; Ex. Neurons, Excitatory Neurons; Inh. Neurons, Inhibitory Neurons; Imm. ChP, Immature Choroid Plexus; ChP, Choroid Plexus. Significance was tested with two tailed Mann-Whitney comparison between treatment and vehicle. P-values: **** <=0.0001, ** <=0.01, * <=0.05, ns >0.05

We next sought to characterize the effects of dex on different neuronal populations. To this end, we labeled deep-layer VI and V neurons with TBR1 (T-box brain transcription factor) and BCL11B (BAF Chromatin Remodeling Complex Subunit-also known as CTIP2) respectively and upper layer IV neurons with SATB2 (SATB Homeobox 2, Figure 1E). Dex consistently led to a significant increase of upper-layer SATB2 positive neurons in hCOs derived from both iPSC lines compared to veh-treated ones. No significant change was found in deep layer neurons (Figure 1F). To further validate these results we used FCa and found increased numbers of upper-layer IV neurons (SATB2+ cells, 9% significant increase, Figure S3A,B) following dex. In addition, dex led to an increase of immature neuronal somata (doublecortin, DCX+) at the basal parts of the germinal zone (Figure S3C,D) and to a decrease of the DCX zone thickness in the cortical-like plate (CP, Figure S3C,E), potentially pointing to later-born neurons still migrating to their final destination. This putatively prolongated neurogenesis can be related to the increased numbers of PAX6+EOMES+ BPs. These BPs have high proliferative capacity, undergoing not only neurogenic but also self-renewing proliferative divisions, which comes in contrast to PAX6-EOMES+ IPs that primarily undergo one neurogenic division producing two neurons^19–24^. Interestingly, PAX6+EOMES+ BPs are abundant in the inner and outer SVZ of mammals with a gyrified brain, such as ferrets, primates and humans, and are one of the mechanisms responsible for the increased neurogenic potential of these species^20^. In lissencephalic species, like rodents, this cell type is rare with the vast majority of BPs being IPs^19–24^. Overall, GCs seem to increase neurogenic processes that are enriched in gyrified species, highlighting the importance of steroid hormones in regulating developmental processes.

### Transcriptional response to glucocorticoids during neurogenesis

We next aimed to identify genes and pathways responsible for the effects of GCs on neurogenesis. For this we used bulk RNA sequencing of No.1– day 45 hCOs following the exact treatment paradigm as above (100nM of dex or vehicle for 7 days, treatment start at day 38). At a 10% FDR cutoff, fifty genes were DE (Table S4). Given the essential role and developmental specificity of transcription factors (TFs) in determining neurodevelopmental processes^25^, we decided to focus on this class of proteins. Out of the 50 DE genes only 4 were TFs, *TSC22D3* (TSC22 Domain family member 3), *KLF9* (Kruppel-like factor 9), *ZBTB16* (Zinc finger and BTB domain-containing protein 16) and *HEYL* (HEY-like protein) (Figure 1G). To narrow in on progenitor-specific responses, we used two scRNA seq datasets. First, to define cell-type specific expression of these TFs we used an already published dataset of No1 day 30 hCOs^26^. *HEYL* and *KLF9* were very lowly expressed in the majority of cell types, *TSC22D3* was found across all cell types and only the expression *ZBTB16* coincided with *PAX6* positive cells (Figure S4D,E and Table S8). To study cell-type specific effects of extended dex treatment on these TFs we used the scRNA seq dataset of HPS0076 hCOs with chronic dex stimulation. *ZBTB16* was differentially overexpressed only in the Radial glia and Cycling progenitors clusters (Figure 1I,J & Table S5), and thus the strongest candidate for dex effects on neural progenitors. *ZBTB16* (also known as *PLZF*-promyelocytic leukemia zinc finger protein) has been associated with regulation of the balance between self-renewal and differentiation of stem and progenitor cells in multiple organ systems including the brain^27^. Thus, we decided to focus on this TF. It belongs to the Krüppel-like zinc finger family of TFs and is very well conserved across species. It has nine zinc finger motifs in the C terminus that comprise the DNA binding domain of the protein, a protein-protein interaction domain at the N terminus and a less well characterized middle RD2 (repressor domain 2) domain and can act as a transcriptional activator or repressor^27^.

### Glucocorticoids alter the very dynamic neurodevelopmental expression pattern of *ZBTB16*

*ZBTB16* is dynamically expressed during rodent and human neurodevelopment with high brain expression early in gestation and a subsequent downregulation to low levels. In rodents, *Zbtb16* is expressed until E10.5 (embryonic day 10.5) in the forebrain^28^ when it is downregulated to non-detectable levels during neurogenesis (Figure S4A,B). In contrast, in human fetal cortex *ZBTB16* is expressed during the initial stages of neurogenesis (Figure S4C), indicating expression of this TF during the neurogenic period in humans but not in rodents.

We analyzed the ZBTB16 expression pattern in our model of the developing human neocortex, the hCOs. ZBTB16 was dynamically expressed in hCOs with high RNA (Figure 2A) and protein (Figure 2B) levels at the early stages of organoid development, until approximately day 40, with a subsequent decrease when mature neurons emerge (MAP2+ cells, Microtubule-associated protein 2) (day 50, Figure 2A). We found ZBTB16 enriched in the apical and basal side of the VZ, mainly expressed by progenitor cells (SOX2+ cells, SRY-Box Transcription Factor 2) and not expressed by mature neurons (MAP2+) (Figure 2C). Thus, *ZBTB16* exhibits a very dynamic expression pattern in hCOs with high protein expression during the initial period of neurogenesis, consistent with data reported from human fetal cortex (Figure S4C).

**Figure 2.**
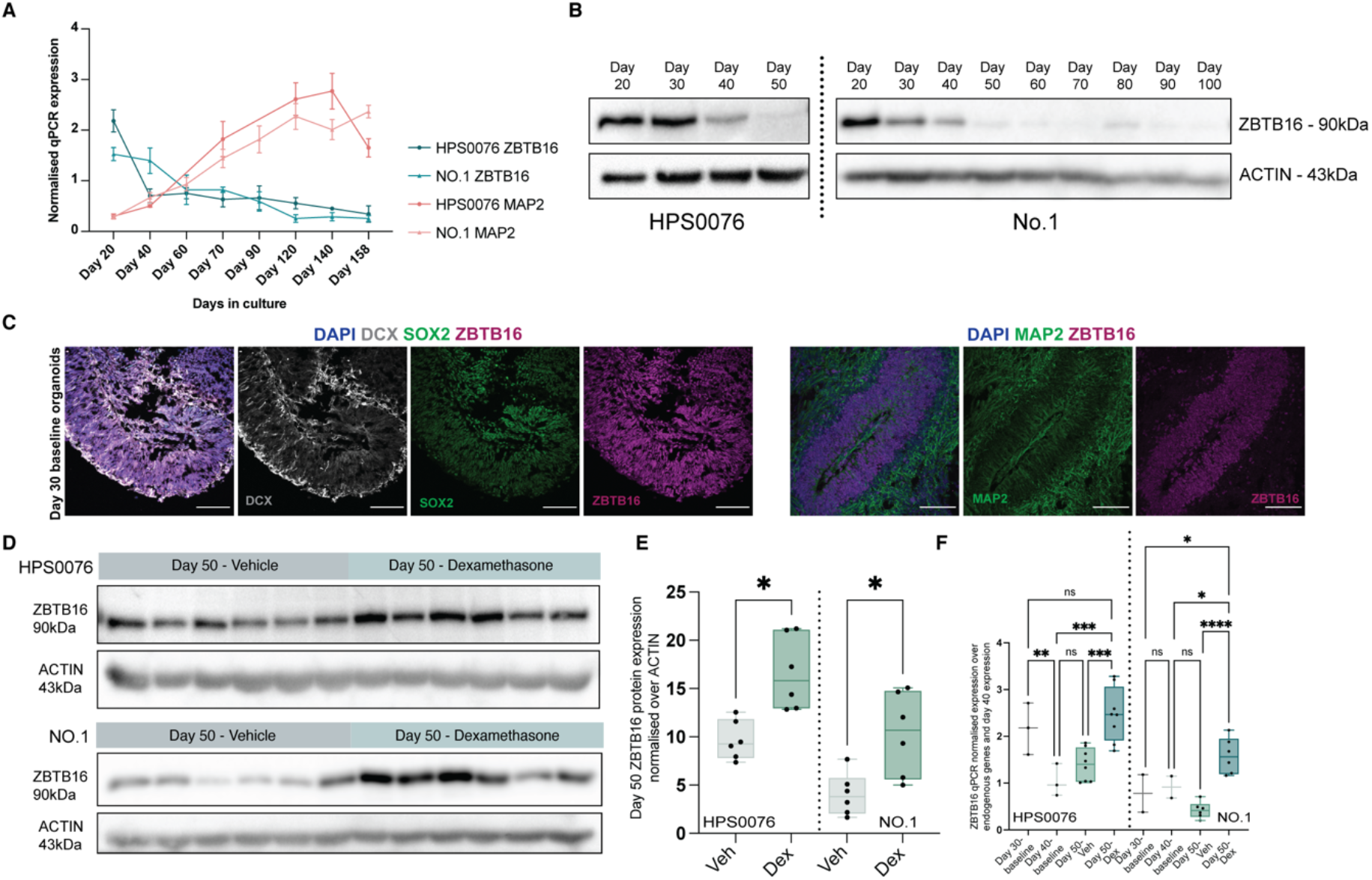
| Glucocorticoids alter the expression profile of *ZBTB16*. **A**, qPCR results (mean +/-SEM) for ZBTB16 and MAP2 in hCOs of different developmental stages. **B,** Western blots of ZBTB16 and ACTIN proteins in hCOs of different developmental stages. **C,** Representative images of day 30 baseline hCOs stained for the immature neuron marker DCX, the progenitor marker SOX2, the mature neuronal marker MAP2, ZBTB16 and the nuclear marker DAPI Scale bars, 100μm. **D,** Western blots of ZBTB16 and ACTIN proteins in hCOs treated with 100nM of dex at day 43 and analysed 7 days later at day 50. Each lane contains protein from a pool of 3 organoids and the 6 replicates were generated in two independent hCOs batches, thus showing protein of 18 organoids from two independene batches in each treatment paradigm. **E**, Quantification of the effect of 100nM 7 days dex treatment on ZBTB16 protein expression in day 50 hCOs normalized over ACTIN. **F,** qPCR plot for ZBTB16 mRNA expression in hCOs and expression at days 30-50, with vehicle and dex condition at day 50. qPCR, quantitative polymerase chain reaction; hCOs, human cerebral organoids; Veh, vehicle; Dex, dexamethasone. For **E** significance was tested with two tailed Mann-Whitney comparison between treatment and vehicle. For **F** significance was tested with one-way ANOVA with Benjamini, Krieger and Yekutieli multiple testing correction (p= 0.0003, F= 10.39, DF= 3). Mann-Whitney p-values for E or post-hoc p-values for F: **** <=0.0001, *** <=0.001, ** <=0.01, * <=0.05, ns >0.05

Given the specificity of *ZBTB16* expression in progenitor clusters, we took advantage of the chronic scRNA seq data to investigate the effects of *ZBTB16* expression in the transcriptional signature of progenitor cells. We sub-setted our dataset to the progenitor clusters in the vehicle condition and performed a DE analysis for ZBTB16-positive versus negative cells (Table S6). We found 442 DE genes at a nominal p-value cutoff of 0.05, one of which was PAX6 (p-value 0.003). Gene Set Enrichment analysis revealed significantly enriched gene sets associated with neuron differentiation, regulation of microtubule organization and transcription highlighting the importance of *ZBTB16* in regulating proliferative and neurogenic processes (Figure S4F, Table S7).

Next, we examined the levels of ZBTB16 expression after dex administration in hCOs (100nM dex, for 7 days starting at day 43-Figure S5A). Treatment with dex resulted in increased expression of *ZBTB16* at the RNA level (Figure S5B, similar to what we have previously reported after 4– and 12– hours dex treatments^26^) and at the protein level (Figure 2D,E) in the progenitor cells that line the ventricular-like zone (VZ, Figure S5C,D). This strongly resembles what we observed at a single-cell genome wide level (Figure 1I,J). In fact, dex alters the tightly-regulated developmental expression pattern of this TF by reversing its levels to those of day 30 and younger hCOs (Figure 2F). Together, these results suggest that dex maintains high *ZBTB16* expression during later stages of neurogenesis, at developmental time-windows with physiologically lower levels of this TF, in the progenitor cells that line the germinal zones.

### *ZBTB16* mimics the effects of glucocorticoids on basal progenitors

To test whether the effect of dex on BPs is mediated via *ZBTB16*, we overexpressed ZBTB16 and GFP from a bicistronic plasmid or GFP from a monocistronic control plasmid, in hCOs starting at day 43 when ZBTB16 expression is already declining (Figure 2A,B). Subsequent analyses were performed 7 days after the electroporation, at day 50 (Figure 3A). In order to explore effects on progenitor subtypes of the VZ and the SVZ and on neurons of the CP, we divided the electroporated area in three bins of equal height starting from the most apical part of the VZ and stopping at the outermost electroporated cell in the CP and analysed the effects of ZBTB16 overexpression on GFP+ electroporated cells.

**Figure 3.**
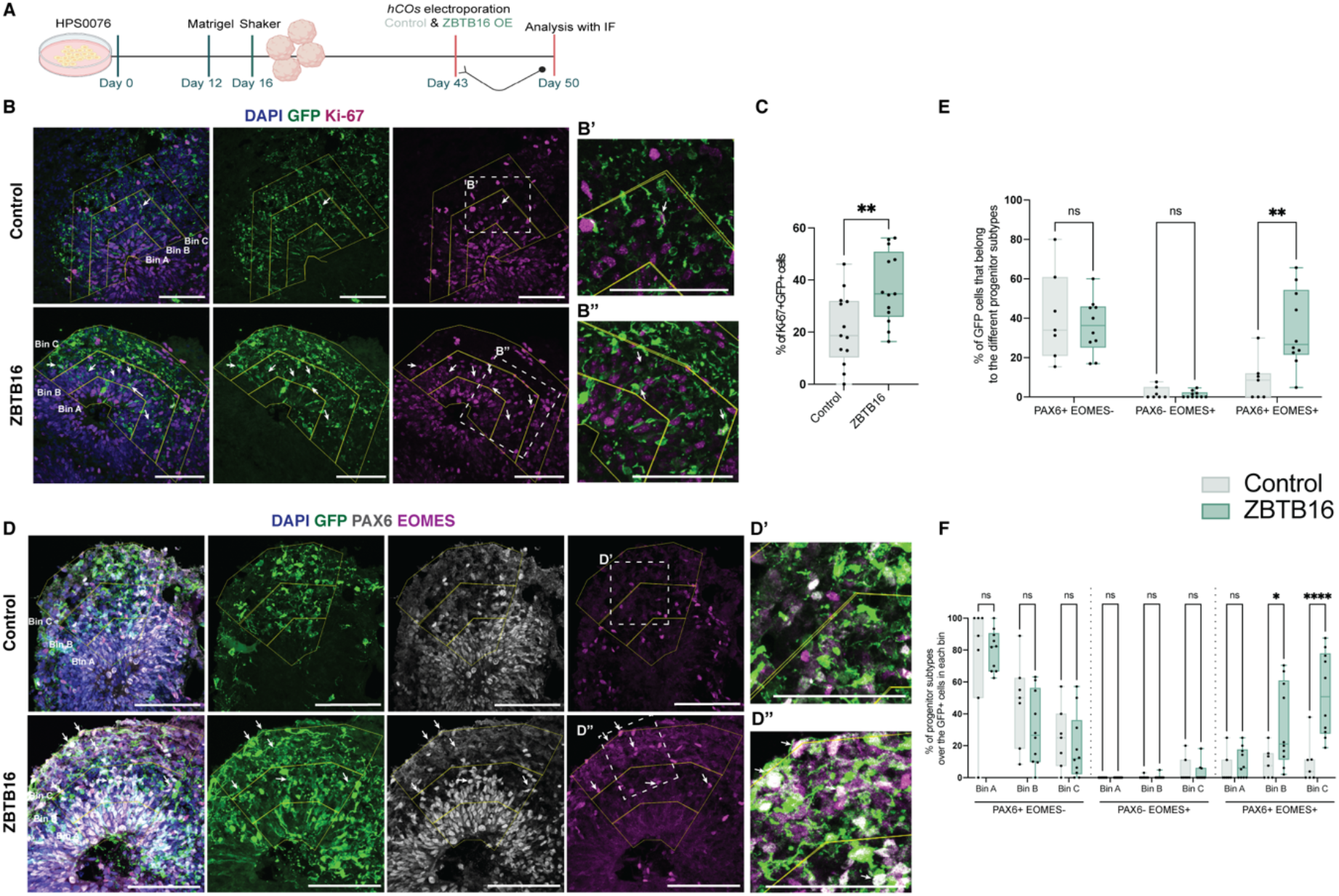
| *ZBTB16* increases PAX6+EOMES+ basal progenitors in human cerebral organoids. **A**, Schematic of HPS0076-derived hCOs electroporations and analysis workflow for ZBTB16 OE. **B,** Representative images of day 50 hCOs at control and ZBTB16 OE conditions stained for Ki-67, GFP and DAPI. White arrows indicate GFP cells that express Ki-67; Scale bars, 100μm. **C,** Quantification of the total number of GFP cells that are Ki-67 positive normalized over total GFP cells. **D,** Representative images of day 50 hCOs at control and ZBTB16 OE conditions stained for PAX6, EOMES, GFP and DAPI. Arrows indicate GFP cells that co-express PAX6 and EOMES; Scale bars, 100μm. **D’ and D”,** Zoom-ins of the areas shown in control and ZBTB16-overexpressing ventricles respectively in Figure D. Scale bars, 100μm. **E,** Quantification of the GFP cells belonging in each progenitor subtype normalized by total GFP cells. **F,** Quantification of the GFP cells belonging in the different single positive progenitor subtypes in each bin and condition normalized by total GFP cells of each bin. hCOs, human cerebral organoids; OE, overexpression; IF, Immunofluorescence. For **C** significance was tested with two tailed Mann-Whitney comparison between ZBTB16-overexpression and control plasmid. For **E&F** significance was tested with two-way ANOVA with Benjamini, Krieger and Yekutieli multiple testing correction (E: p.interaction= 0.0069, F= 5.5, DF= 2/ F: p.interaction = 0.0002, F= 4.17, DF= 8). Mann-Whitney p-values for C or post-hoc p-values for E&F: **** <=0.0001, ** <=0.01, * <=0.05, ns >0.05

ZBTB16 overexpression led to increased numbers of Ki67+ cells (Figure 3B,C), indicating an increase in proliferation potential. In analogy to the experiments with dex, we next co-analyzed PAX6 and EOMES expression. Indeed, ZBTB16 overexpression led to a similar phenotype to that of dex, with an overall 25.7% increase in PAX6+EOMES+ BPs (Figure 3D,E) in bins B and C (Figure 3F, 23.8% increase in bin B and 43.1% increase in bin C), which reflect the basal parts of the VZ, the SVZ and the CP. Interestingly, there was a substantial increase of these BPs in bin C (43.1%) which associates to the CP of the organoids, pointing to a potential identity change of this bin to resemble an area with cells typically found in the inner and outer SVZ of gyrencephalic species. Using GFP cell morphology reconstructions, we found that the double-positive BPs exhibit both IP-related morphologies with no processes and bRG-related morphologies with monopolar or bipolar cells (Figure S6, see Supplemental Results for more details), similar to what has been described before for these double positive cells in macaques and humans^20, 29^.

We next studied the effects of ZBTB16 overexpression on deep-layer V and upper-layer IV neurons by staining with BCL11B and SATB2 respectively. We found a significant 19% increase of SATB2+ cells after ZBTB16 overexpression (Figure S7B,C), whereas the effect of dex on BCL11B+ deep-layer neurons was less-pronounced (non-significant 11% increase, Figure S7A,C), indicating increased neuronal production. Thus, ZBTB16 overexpression increases the amount of double-positive BPs and of upper-layer neurons, effects that resemble the ones of dex (Figure 1E,F).

### *ZBTB16* is necessary for the effects of glucocorticoids on basal progenitors

In view of the similarity of the dex and ZBTB16 overexpression phenotypes on BPs, we sought to determine whether ZBTB16 is in fact necessary for the dex-induced phenotype. To achieve this, we used CRISPR-Cas9 to knock-out (KO) exon 2 of the *ZBTB16* locus in the No.1– iPSCs. Exon 2 encodes for more than 50% of the protein and includes the initiating ATG, the BTB/POZ domain and the first two zinc fingers of the binding domain^30^. We created heterozygous No.1– (from now on called ZBTB16^+/-^) iPSCs where one allele of the *ZBTB16* locus is wild-type and one allele has exon 2 excised. A full KO of exon 2 in both alleles was not viable at the iPSC cell stage. We then treated ZBTB16^+/+^– derived (CRISPR control No.1– iPSCs) and ZBTB16^+/-^-derived day 43 hCOs with 100nM dex for 7 days and analysed ZBTB16 protein expression as well as the relative abundance of PAX6+EOMES-, PAX6-EOMES+ and PAX6+EOMES+ progenitor subtypes with FCa and immunofluorescence at day 50 (Figure 4A). ZBTB16^+/-^ hCOS showed significantly less increase of the ZBTB16 protein following dex treatment than the ZBTB16^+/+^ hCOs (46% less increase, Figure 4B,C). FCa of the ZBTB16^+/+^ hCOs validated the increase of PAX6+EOMES+ BPs under dex treatment (Figure 4D, E-22.2% increase, similarly to what we found with the No.1 wild-type hCOs (Figure S2B,C). However, in the ZBTB16^+/-^ hCOs, the number of double-positive BPs was not significantly increased in the dex condition in respect to veh (Figure 4F,G-6% non-significant increase). This was further validated at the immunofluorescence level where dex significantly increased the numbers of PAX6+EOMES+ BPs in ZBTB16^+/+^ hCOs but not in ZBTB16^+/-^ hCOs (Figure S8), recapitulating the effects we see with our FC analysis. Together, these results indicate that ZBTB16 is necessary for the effects of GCs on PAX6+EOMES+ BPs and suggests that it could potentially play a key role in the maintenance of the PAX6+EOMES+ BP population in gyrified species under baseline conditions.

**Figure 4.**
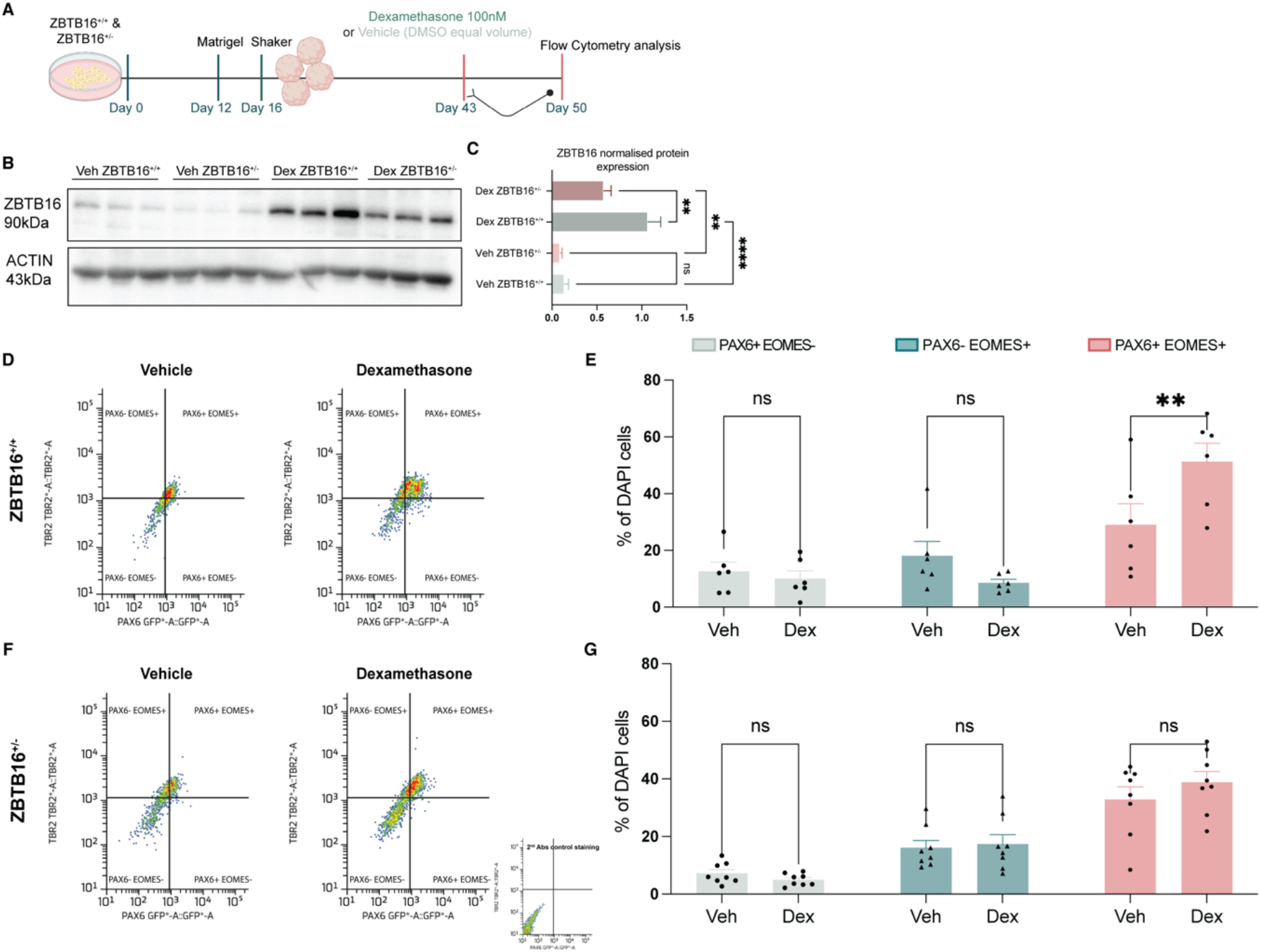
| *ZBTB16* mediates the effects of glucocorticoids on PAX6+EOMES+ basal progenitors. **A**, Treatment paradigm and analysis workflow in hCOs derived from edited No.1– iPSCs either with ZBTB16^+/+^ or ZBTB16^+/-^ genotypes. **B,** Western blots for ZBTB16 and ACTIN proteins in ZBTB16^+/+^ or ZBTB16^+/-^ derived hCOs at different treatment conditions. Each lane contains protein from a pool of 3 organoids thus showing protein of 9 organoids in each treatment group and genetic background. **C,** Quantification of the Western blot results for ZBTB16 normalized over ACTIN. **D,** Representative images of FCa of ZBTB16^+/+^-derived hCOs per dex treatment condition. TBR2 is an alternative name for EOMES. **E,** Quantification of the FCa results. Percentages of DAPI cells in each progenitor subtype and treatment condition. **F** Representative images of FCa analysis of ZBTB16^+/-^-derived hCOs per treatment condition. **G,** Quantification of the FCa results. Percentages of DAPI cells in each progenitor subtype and treatment condition. DMSO, dimethyl sulfoxide; hCOs, human cerebral organoids; Veh, vehicle; Dex, dexamethasone; FCa, flow cytometry analysis. For **C,E&G** significance was tested with two-way ANOVA with Benjamini, Krieger and Yekutieli multiple testing correction (C: p.interaction = 0.03, F= 6.6, DF= 1/ E: p.interaction = 0.0068, F= 5.9, DF= 2/ G: p.interaction = 0.97, F= 0.38, DF= 2). Post-hoc p-values: ** <=0.01, ns >0.05

### Heterochronic *ZBTB16* expression in mouse fetal brain is sufficient to induce basal progenitors typically enriched in gyrified species and increase neurogenesis

While the principles of neurogenesis are similar among all mammalian species, differences exist in respect to their temporal progression as well as to the different progenitor subtypes populating the SVZ which play a key role in the overall neurogenic potential. In lissencephalic species BPs double-positive for PAX6 and EOMES are very rare^19^, the neurogenic period is much shorter^31^ (9 days in mice compared to 110 days in humans) and *ZBTB16* is not physiologically expressed at any point during the neurogenic period (Figure S4C,D). In order to analyse if altered expression of *ZBTB16* during neurogenesis would lead to increased numbers of PAX6+EOMES+ BPs also in a lissencephalic species, we performed *in utero* electroporations at E13.5 in mouse brains with the same plasmids as for the hCOs and analysed the effects of human *ZBTB16* overexpression at E16.5 (Figure 5A). Considering the more complex cortical cellular architecture of the mouse brain as compared to hCOs, we divided the electroporated area in five bins of equal height where bin A includes the VZ and the SVZ, bin B the SVZ and intermediate zone (IZ), bin C the IZ and bins D and E the CP.

**Figure 5.**
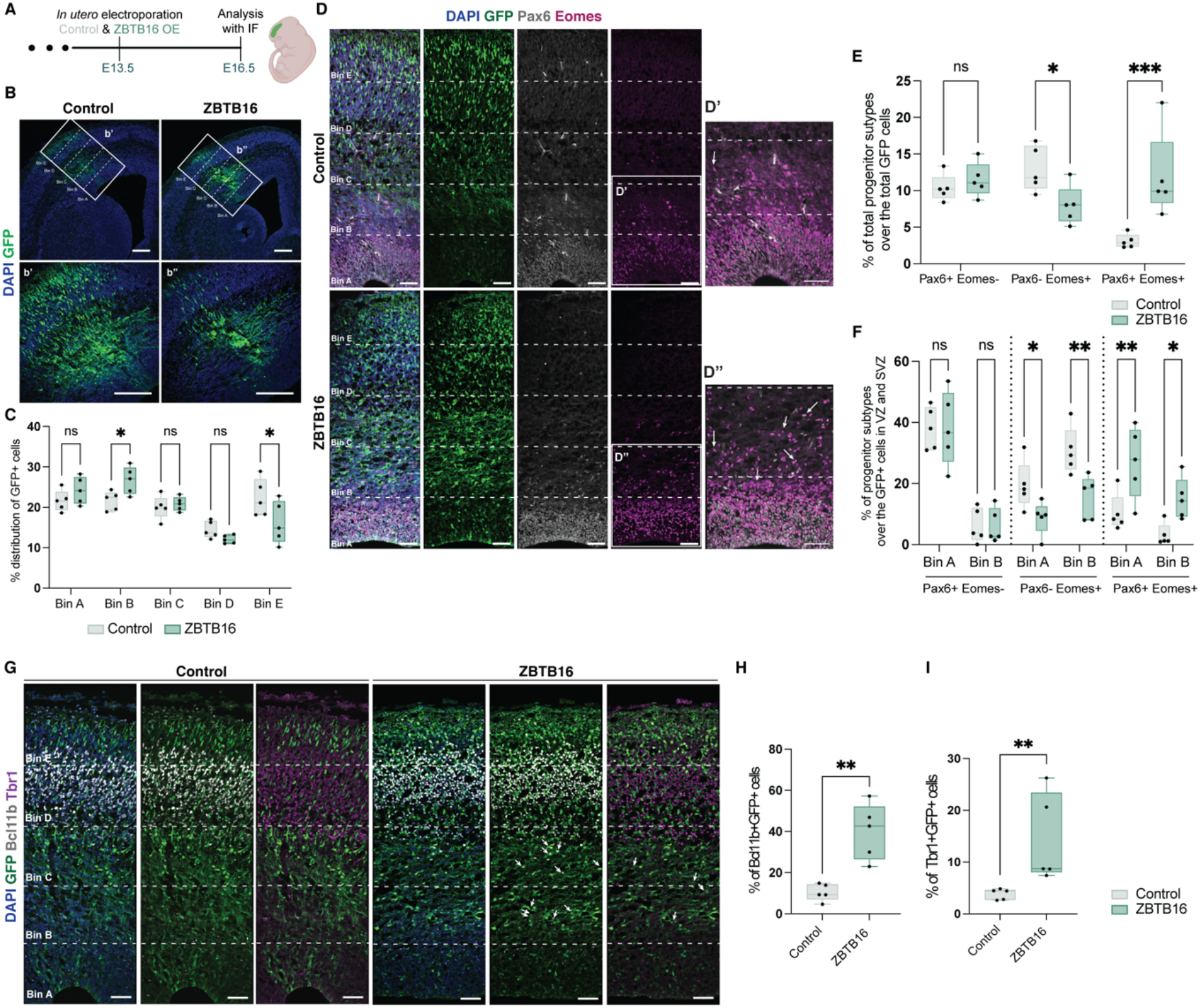
| *ZBTB16* increases PAX6+EOMES+ basal progenitors and deep layer neurons in a lissencephalic species. **A**, Workflow of the *in utero* electroporations of *ZBTB16* and analysis in fetal mice. **B,** Representative images of E16.5 fetal mouse brains at control and ZBTB16 OE conditions stained for GFP and DAPI. Box indicates the electroporation areas and **b’, b’’** are zoom-ins; Scale bars, 100μm. **C,** Quantification of the distribution of GFP cells in each bin normalized by the total number of GFP cells. **D,** Representative images of E16.5 fetal mouse brains at control and ZBTB16 OE conditions stained for Pax6, Eomes, GFP and DAPI. **d’ and d’’** are zoom-ins. Arrows indicate GFP cells that co-express Pax6 and Eomes; Scale bars, 50μm. **E,** Quantification of the GFP cells belonging in the different progenitor subtypes normalized by total GFP cells. **F,** Quantification of the GFP cells belonging in the different positive progenitor subtypes in each bin normalized by total GFP cells of each bin. **G,** Representative images of E16.5 fetal mouse brains at control and ZBTB16 OE conditions stained for the layer V neuronal marker Bcl11b, layer VI neuronal marker Tbr1, GFP and DAPI. Arrows indicate GFP cells that express Bcl11b or that express Tbr1; Scale bars, 50μm. **H,** Quantification of the GFP cells that are Bcl11b+ normalized by total GFP cells. **I,** Quantification of the GFP cells that are Tbr1+ normalized by total GFP cells. OE, overexpression; IF, Immunofluorescence. For **H & I** significance was tested with two tailed Mann-Whitney comparison between ZBTB16-overexpression and control plasmid. For **C, E & F** significance was tested with two-way ANOVA with Benjamini, Krieger and Yekutieli multiple testing correction (C: p.interaction = 0.003, F= 4.7, DF= 4/ E: p.interaction = 0.0003, F= 11.84, DF= 2/ F: p.interaction < 0.0001, F= 6.58, DF= 5). Mann-Whitney p-values for H or post-hoc p-values for C,E &F: *** <=0.001, ** <=0.01, * <=0.05, ns >0.05

In mice, ZBTB16 overexpression significantly changed the distribution of the GFP+ electroporated cells. ZBTB16+GFP+ cells accumulated in the SVZ with less cells reaching the outer-most part of the cortical plate (Figure 5B,C), indicating possible identity changes and/or altered timing of differentiation. Similar to hCOs, we found more Pax6+Eomes+ BPs (Figure 5D,E, 8.8% overall increase) in bin A and bin B (Figure 5F, 16.5% increase in bin A and 11.8% increase in bin B). Interestingly, in mice, Pax6-Eomes+ IPs, which are the vast majority of endogenous BPs of lissencephalic species^19^, were significantly decreased after ZBTB16 overexpression (Figure 5E,F, 10.6% decrease in bin A and 15.1% decrease in bin B). In addition, ZBTB16 seems to increase the self-renewing capacity of the cells it is overexpressed in as we observed a significant increase of BRDU+Ki67+ cells in bins A and B compared to control electroporations (cell cycle re-entry analysis with BRDU for 24 hours, Figure S9, see Supplemental Results for more details). Our results thus suggest that ZBTB16 overexpression in lissencephalic neurogenesis leads to increased gyrified species-enriched BPs at the expense of the endogenous neurogenic progenitors.

To further study what this means for neurogenesis we analyzed the neuronal output. Here, we found that ZBTB16 overexpression led to increased numbers of layer V neurons expressing Bcl11b (33.3% significant increase, Figure 5G,H) and layer VI neurons expressing Tbr1 (4.3% significant increase, Figure 5G,I) but not of upper layer IV neurons that express Satb2 (Figure S10C,D). Given that in mice it is known that upper layer neurons are mainly born starting at E14.5 and not before^32^, we repeated the same experiments but performing the IUEs a day later, at E14.5, and analyzing neuronal output at E17.5. Indeed, at this developmental stage ZBTB16 overexpression leads to increased numbers of Satb2+ cells compared to the control plasmid (14% significant increase, Figure S10F,G). We also still find increased numbers of Bcl11b+ cells (26% significant increase, Figure S10G). Thus, in mice ZBTB16-overexpression leads to increased numbers of both deep– and upper-layer neurons whereas in hCOs the main effect of both dex and ZBTB16-overexpression is on upper-layer neurons, potentially highlighting differences in the plasticity of these cells in a gyrified-or lissencephalic-context. When examining the distribution of neurons, we found an accumulation in the SVZ– and IZ– areas with less neurons having reached the CP after ZBTB16 overexpression (Figure S10A,B,E). This finding again lends support to the fact that the higher proliferative capacity of the PAX6+EOMES+ BPs associates with a potentially longer neurogenic period similar to what we observed following dex treatment in hCOs. In fact, when analyzing the distribution of neurons across the five bins not 3 but 6 days post electroporation (i.e. at day E19.5/birth) there were no significant differences for the distribution of GFP+ cells with ZBTB16 overexpression (Figure S10H-L), indicating that migratory processes are probably not affected.

### *ZBTB16* directly induces *PAX6* expression

Considering that dex via *ZBTB16* seems to sustain *PAX6* expression in EOMES+ cells, even in a lissencephalic species where physiologically they are mutually exclusive^33–35^, and that *ZBTB16* is a TF, we analysed the activator capacity of ZBTB16 on *PAX6* human promoters. *PAX6* has three promoter regions that regulate tissue-specific expression and that are highly conserved between humans and rodents: the P0, P1 and Pa promoters ^36, 37^ (Figure S11A). Using luciferase assays, we found that ZBTB16 activates the P1 promoter of *PAX6* but not the P0 and Pa promoters (Figure S11B). This suggests that ZBTB16 could regulate *PAX6* expression via the P1 promoter which is active during neocortical development, in comparison to the P0 and Pa promoters which are minimally active^36^. This could be a potential mechanism responsible for sustaining *Pax6* expression in Eomes+ cells even in mice by ZBTB16 overexpression circumventing the negative feedback loop that physiologically ensures that *Pax6* and *Eomes* are not co-expressed in rodents^33–35^.

### Glucocorticoids interact with the *ZBTB16* genetic and epigenetic landscape

In order to analyse the mechanisms via which the physiological temporal expression pattern of *ZBTB16* is affected by GCs, we looked at the gene regulatory landscape of the human *ZBTB16* locus in response to GCs. ENCODE data indicate the existence of intronic GR-response elements (GREs) at the human *ZBTB16* locus (Figure 6A). We have previously shown that activation of GR leads to DNA methylation changes in GREs of target genes which associates with changes in gene transcription^38, 39^. To identify the molecular mechanism by which dex via GR induced *ZBTB16* expression, we treated day 30 hCOs with 100nM dex for 7 days and used HAM-TBS^40^ (Highly Accurate Method for Targeted Bisulfite Sequencing) and Pyrosequencing to measure DNA methylation of all GREs, as identified by public GR-ChIP sequencing datasets (Table S9) and additional non-GR-related regulatory elements (Figure 6A). Out of 55 CpGs covered with HAM-TBS and 8 covered with Pyrosequencing, 44 were located within GREs and 19 in enhancer elements lacking GR binding sites. While only 1 of the CpGs outside these GR-binding regions showed significant DNA methylation level changes following dex stimulation (Table S10), this was true for 18 of the 44 CpGs around GREs (Figure 6B and Table S10). All significantly altered GRE-CpGs are located in enhancer regions that loop to the transcriptional start site of *ZBTB16* (“GeneHancer” track in Figure 6A and Bothe *et al*., Life Science Alliance, 2021^41^). Our data thus support a model in which GR binds to these enhancer elements as evidenced by altered DNA methylation and increases *ZBTB16* transcription.

**Figure 6.**
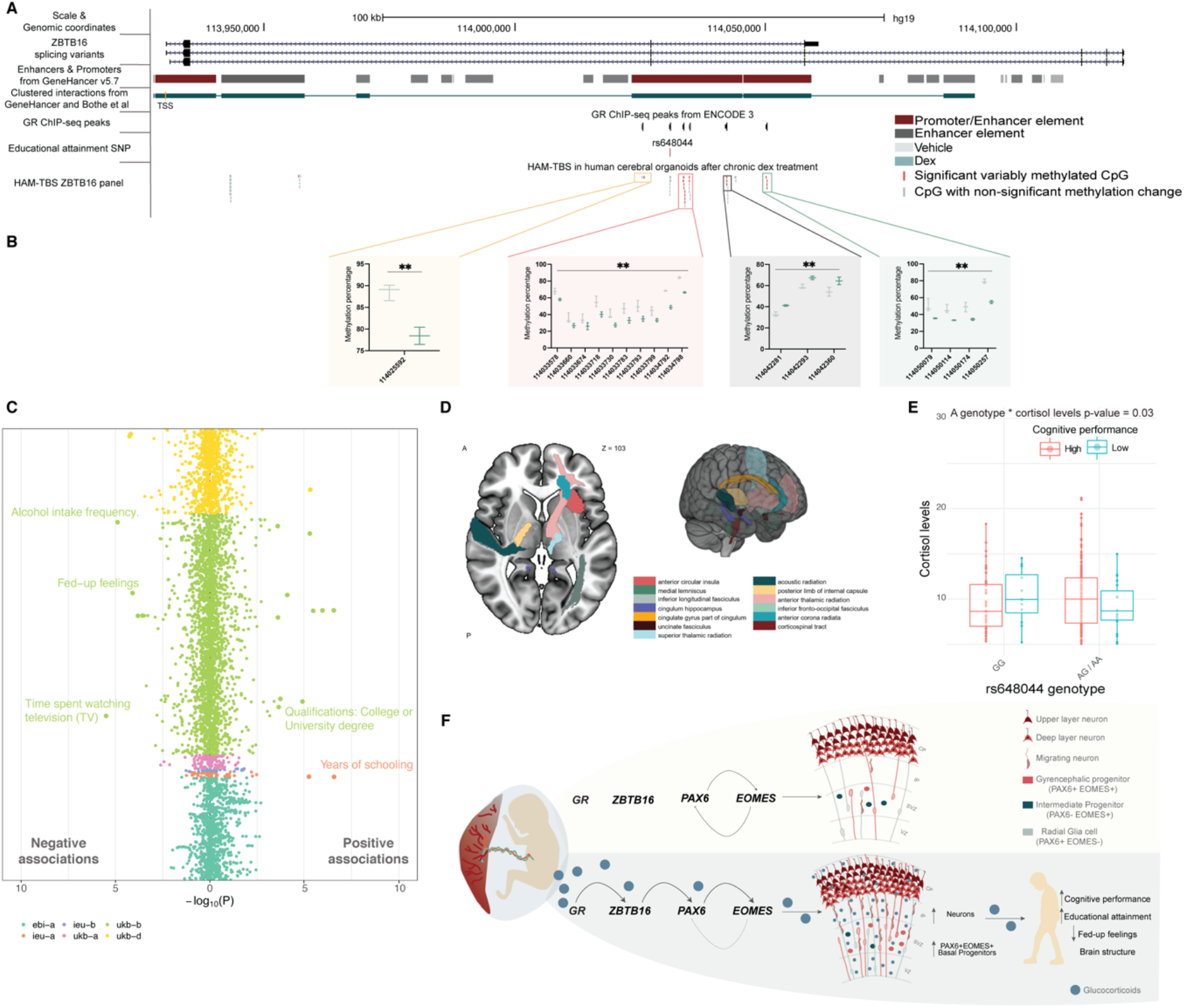
| Glucocorticoids interact with the genetic and epigenetic landscape of ZBTB16 to impact postnatal neurobehavioral and structural phenotypes. **A**, Graphical representation of the ZBTB16 locus including the position of the amplicon for HAM-TBS and Pyrosequencing. **B,** HAM-TBS results for CpGs with significantly altered DNA methylation levels following exposure to 7 days of 100nM dex vs. vehicle. **C,** Plot depicting Phe-WAS-MRa associations as –log of the p-value for rs648044 with various phenotypes from the UK Biobank. Single phenotypes are depicted as individual points. Associations are presented based on negative (negative MRa estimate.i.e. lower quantitative measures with A-allele effects) and positive (positive MRa estimate, i.e. higher quantitative measures with A-allele effects) effects. Color coding reflects the different sources of GWAS depicted in the legend. Traits that remain significant following Benjamini-Hochberg correction are shown with larger dots and are listed in Table S14. All neurobehavioral traits are labeled. **D,** Illustration of significant MRa associations between brain imaging phenotypes and rs648044 mapped onto the brain atlases. All effects are listed also in Table S15. **E,** Cortisol levels before GW28 according to rs648044 genotype and cognitive performance as assessed using the Bayley-III cognitive subscale in the data from the ITU cohort. Total cortisol output was estimated based on the area under the curve with respect to ground (AUCg) of all seven saliva samples using the trapezoid rule in 246 mothers. **F,** Summary of the effects of glucocorticoids on ZBTB16 expression, cortical cellular architecture during development and postnatal phenotypes. GR, glucocorticoid receptor; ChIP, chromatin immunoprecipitation; Seq, sequencing; SNP, single nucleotide polymorphism; HAM-TBS, highly accurate method for targeted bisulfite sequencing; Dex, dexamethasone; MRa, mendelian randomization analysis. For **B** significance was tested with two-way ANOVA with Benjamini, Krieger and Yekutieli multiple testing correction (GH11J114152 enhancer: p.treatment < 0.0001, F= 7.33, DF= 31/ GH11J114174 enhancer: p.treatment< 0.0001, F= 37.32, DF= 3). Post-hoc p-values: ** <=0.01

Knowing that environmental factors, including GCs, can interact with the genetic landscape to modulate their effects on expression^42^, we next analyzed the genetic landscape of *ZBTB16*. We catalogued the SNPs (single nucleotide polymorphisms) in the *ZBTB16* locus previously associated with neurobehavioral and brain structural outcomes (GWAS Catalogue & Table S11) and identified rs648044 as the only variant associated with both (Table S12). rs648044 has been associated with educational attainment in two GWAS (genome-wide association study) for this trait (Lee *et al*.^43^: N= 1,131,881, FDR = 9*10^-9^ & Okbay *et al*.^44^: N= 3,037,499, FDR = 2*10^-8^) and with generalized cortical thickness^45^ (N= 35,657 individuals, FDR = 6*10^-9^). In the latter GWAS on cortical morphology, gene-level analysis also identified the whole *ZBTB16* locus to be significantly associated with generalized cortical surface area (FDR = 7.2*10^-14^) and thickness (FDR = 1.9*10^-8^), thus suggesting relevance of *ZBTB16* for adult cortical morphology.

In our DNA methylation assays, CpGs surrounding this SNP showed low methylation levels supporting its regulatory activity (Figure S12A and Table S10). To further explore its role in gene regulation, we first analyzed whether this SNP moderates dex-induced activity of the surrounding 200 bp (base pair) enhancer element using a STARR-qPCR (Self Transcribing Active Regulatory Region-qPCR) approach. We identified that the sequence surrounding rs648044 indeed possesses enhancer activity that is increased with GR activation via dex (Figure S12B). The extent of the dex-induced activity increase was rs648044 allele-dependent, with the allele associated with higher educational attainment (A-allele), conferring a significantly stronger increase following dex (Figure S12C). Next, we used CRISPR-Cas9 to KO 400 nucleotides surrounding rs648044 in No.1– iPSCs in order to identify whether this enhancer affects *ZBTB16* transcription in hCOs. The rs648044 genotype of No.1-iPSCs is heterozygous (G/A), with the rarer A allele (allele frequency = 0.35) creating a degenerated partial GR binding site (described half site^46^: AGXACAG, rs648044 creates: AGC<u>A</u>GAG, Figure S12D). A KO of the A allele resulted in rs648044^G/-^ cells that only carry the G allele (Figure S12D). Using the edited cell line (rs648044^G/-^) and the control cell line carrying both alleles (rs648044^G/A^), we found that the absence of the A allele confers significantly less induction of *ZBTB16* following 100nM dex in day 30 hCOs treated for 7 days (Figure S12E), suggesting that the enhancer containing rs648044 modulates dex-mediated transcriptional effects on *ZBTB16* in hCOs.

### Glucocorticoids x rs648044 effects on *ZBTB16* relate to beneficial postnatal outcomes

Given the functional effects of rs648044 on GC-induced ZBTB16 expression in the developing cortex, we used phenome-wide mendelian randomization analysis (MRa-PheWAS) to identify causal effects of *ZBTB16* levels on 7,323 phenotypes from the UK Biobank and the NHGRI-EBI GWAS Catalog (Table S13) that include, among many others, neurobehavioral traits and adult neuroimaging data. We used rs648044 as exposure and the magnitude of the allele-specific expression changes following dex in the STARR-qPCR experiment as instrument to test for these associations. MRa-PheWAS provided strong evidence for associations with multiple outcomes as indicated by the QQ-plot (Figure S12G). MRa on various phenotypes, including endophenotypes and diseases (N= 4,360), showed significant associations of GC-altered *ZBTB16* expression with 22 phenotypes after multiple testing correction. These included positive associations with years of schooling and whether individuals had obtained a college or university degree (Figure 6C and Table S14), which are direct measures of educational attainment. This supports the idea that the functional effects of rs648044 on GC-induced *ZBTB16* transcription in the developing cortex, are putatively causally-related to educational attainment postnatally. In addition, GC-induced *ZBTB16* transcription was negatively associated with “fed-up feelings” a phenotype related to neuroticism^47^, “alcohol intake” and “time spent watching television” (Figure 6C), suggesting associations of higher *ZBTB16* levels with beneficial postnatal outcomes and decreased associations with negative outcomes. Given the previously published relationships of both educational attainment^48–51^ as well as rs648044^43–45^ with cortical volumes and white matter measures, we also ran an MRa-PheWAS on all neuroimaging phenotypes (N = 3,143) in the UK Biobank using the same instrument. We observed 21 significant associations after multiple testing correction (Figure 6D, Figure S12H and Table S15). Most evidence indicated that GCs x rs648044-mediated increases in *ZBTB16* expression were significantly associated with altered white matter measures and with higher anterior circular insula thickness (Figure 6D).

To further examine the importance of GC-altered *ZBTB16* levels during neurogenesis for postnatal phenotypes, we used data from the Intra-Uterine (ITU) sampling in early pregnancy prospective cohort study^52^ to test the association of the rs648044 A allele and cortisol with cognitive performance. The ITU cohort comprises of 944 mother-child dyads. Neuropsychological assessment of the children was performed at 35.6 months of age (range 24.5 to 42.5) using the Bayley-III screening test^53^ designed to identify infants and toddlers at risk for developmental delay. We defined neurodevelopmental cognitive delay, using the Bayley cognitive subscale, as one standard deviation (SD) below the sample mean (Figure S12I). For children testing below and above 1 SD of cognitive performance, a complete set of salivary cortisol data during pregnancy and fetal genotypes was available for 246 mother-child dyads before GW28 and for 221 mother-child dyads after GW28 (Table S16). Using logistic regressions, we tested the interaction of rs648044 A allele * mean cortisol levels before GW28, after GW28 or across pregnancy on cognitive performance at 3 years of age (see Methods for covariates). The interaction was not significantly associated with cognitive performance when using cortisol levels from across pregnancy (p-value 0.055, Table S17). Interestingly, when stratifying the data for cortisol levels before GW28, and thus during neurogenesis, the interaction of rs648044 A allele with cortisol was significantly associated with higher cognitive performance (p-value 0.035, Figure 6E, Table S17), an effect not found with cortisol data after GW28 (p-value = 0.46, Table S17) and thus after the end of neurogenesis.

Thus, higher GC-induced *ZBTB16* expression in rs648044 A allele carriers during neurogenesis might drive the association of GC treatment with higher cognitive performance, higher educational attainment, lower neuroticism measures and increased cortical thickness as well as altered white matter measures postnatally. This suggests that the genetic association of this variant with postnatal and adult neurobehavioral and brain structural measures could be mediated in parts by its effects on GC-induced *ZBTB16* levels in early brain development and in consequence their effects on neurogenesis.

## Discussion

With this work we sought to understand the effects of GCs during human cortical neurogenesis at the cellular and molecular levels. Our results provide a novel molecular and cellular mechanism for GC effects on neurogenic processes and also directly relate it to neurobehavioral, cognitive and brain structural outcomes after birth. We found that GCs increase a particular population of neural progenitor cells that co-express PAX6 and EOMES. These progenitor cells were found at the basal side of the germinal zones and are known to be enriched in species with gyrified brains and related to a higher proliferative capacity, contributing to increased neuronal production^19, 20, 54^. Indeed, we found these BPs to have higher self-renewing capacity while neurogenesis was also increased. We show that these effects of GCs are mediated by an alteration of the developmental expression profile of a TF called *ZBTB16*. *ZBTB16* activates a forebrain active promoter of *PAX6*, potentially explaining how GCs increase the numbers of PAX6+EOMES+ BPs. The increase in these specific BPs is followed by increased numbers of neurons, albeit cell lineage experiments would be needed to directly prove the association, and relates to beneficial postnatal outcomes at the neurobehavioral level, like increased cognitive performance, educational attainment and altered brain structure (Figure 6F).

Precise temporal and spatial regulation of gene expression by TFs is key for the proper unfolding of neurogenic processes^25^. Thus, we focused on GC-effects on TFs as the potential molecular mechanism responsible for their effects on neurogenesis. We showed that GCs alter the tightly regulated developmental expression profile of a TF called *ZBTB16* in hCOs which in turn mediates the GC-effects on neurogenesis. While ZBTB16 shows a dynamic expression pattern in both gyrencephalic and lissencephalic species, the expression among species differs according to developmental time-windows. In mice, *Zbtb16* appears at E7.5 in the neuroectoderm, increases until E10.5 and is subsequently downregulated and finally only expressed in specific areas of the hindbrain and the septum but not the forebrain^28^, so that in rodents *Zbtb16* is not expressed during cortical neurogenesis. This is contrary to the expression pattern observed in human fetal brain and hCOs (Figure S4), where *ZBTB16* is expressed during the initial stages of neurogenesis. This points to possible divergent actions of this TF in gyrencephalic and lissencephalic species neurodevelopment and highlights this protein as important for gyrified-species enriched neurogenic processes at baseline as well as in response to GCs.

Indeed, overexpressing ZBTB16 in the mouse fetal brain during the neurogenic period (E13-E16, when *Zbtb16* is not anymore expressed physiologically) leads to increased Pax6+Eomes+ BPs. In lissencephalic cortical development *Pax6* and *Eomes* create a positive feedforward cascade that self-regulates with direct negative feedback effects^33–35^, ensuring mutually exclusive expression of these proteins. This seems to be overridden by asynchronous overexpression of *ZBTB16*. We also show that *ZBTB16* directly activates the *PAX6* promoter P1^36^, that is functional in the forebrain, and sustains PAX6 expression potentially leading to the increased PAX6+EOMES+ BPs both in hCOs and in mice. Thus, via the action of *ZBTB16,* GCs have the ability to extend or open a sensitive developmental time-window for the production of gyrencephalic-enriched BPs in a lissencephalic species resulting in enhanced neurogenic potential and higher production of neurons. *ZBTB16* seems to specifically mediate the effects of GCs on neurogenesis since it is differentially regulated only in the progenitor clusters. Thus, it is unlikely that it would mediate other reported in the literature GC effects that are specific to neuronal function, as, for example, synaptic plasticity^55, 56^.

Given the high prevalence of premature births, GC excess during human neurodevelopment through administration of sGCs is a very common phenomenon^9^. In fact, in about ∼615,000 extreme preterm pregnancies per year, sGC treatments, if used, would take place in a period of active neurogenesis, before GW28^10, 16^. In fact, it was recently shown that sGCs at GW22 increase the survival rate of the offspring 3.5 times, with the authors further supporting and suggesting the use of SGCs in extremely premature births^11^. While endogenous GCs are important for the maturation and function of fetal organs^4^, prenatal excess of GCs has been extensively associated with long-term metabolic, endocrine and cardiovascular problems^2^ and risk for neurodevelopmental^57^ and mental^58^ disorders in the offspring. Evidence from a recent meta-analysis of studies including more than 1,25 million children re-affirms the association of exposure to sGCs with negative effects on cognitive and neuropsychiatric outcomes when administered to children with late-preterm or term birth. However, the authors also report a significantly lower risk for neurodevelopmental impairments in children with extremely preterm birth (< GW28), that were treated with sGCs between GW22 to GW27^14^. This meta-analysis could be pointing to potential differential effects of GCs on neurodevelopment depending on the developmental time-window they were administered in.

One process that is different among extremely preterm and term or adult brains and might contribute to these dichotomous effects is human cortical neurogenesis, which peaks ∼GW20 and is reduced but present until GW28 in the SVZ of the brain^16^. This means that extremely preterm born children are still treated within the time window of active cortical neurogenesis. Such differential effects are supported by a study in mice in which dex administration during neurogenesis was associated with anxiolytic and anti-depressive like behavior in the adult offspring^59^ while increased, prolonged exposure to GCs following completion of cortical neurogenesis has repeatedly been associated with increased anxiety and depressive-like behaviors and decreased cognitive ability in these animals^60^. With this work we highlight a potential molecular and cellular pathway for the lasting effects of prenatal sGC exposure during neurogenesis and pinpoint *ZBTB16* as an important hub gene mediating GC effect on cortical neurogenesis and beneficial postnatal phenotypes. The latter is, on the one hand, supported by our results using the ITU cohort where we find that that in carriers of the high *ZBTB16*-induction rs648044 allele, increased maternal cortisol levels were associated with better cognitive performance, but only when considering cortisol levels before GW28, i.e. during active neurogenesis, and not at later time points. It is additionally supported from the MRa-PheWAS where we find associations with higher educational attainment, lower neuroticism measures and altered cortical and white matter measures. Nevertheless, the association with postnatal behavioral and structural outcomes should be interpreted with caution as the increased number of neurons may ultimately not contribute to brain function postnatally.

Overall, our work provides a molecular and cellular mechanism of how GCs early in development, during the neurogenic period, affect the cytoarchitecture of the developing cortex which in turn associates with postnatal outcomes at the brain structural but also neurobehavioral level. This work highlights the importance of GCs for neurogenic processes in humans. In addition, the new knowledge offers a possible molecular and cellular pathway explaining the associations of early sGC use with beneficial behavioral and neurodevelopmental measures found in the literature and, thus, may help to refine sGC treatment guidelines according to the stage of pregnancy they are administered in.

## Supporting information

Supplemental Results

## Acknowledgments

LD acknowledges support by the Joachim Herz Foundation. We thank Mira Erhart, dr. Michael Czisch and dr. Benoit Boulat for informative discussions on the brain structural measures. We thank Nathalie Gerstner and dr. Janine Knauer-Arloth for discussion on data analysis. We thank dr. Nadine Gogolla for discussion on broader relevance of the findings. This work was supported by the Hope for Depression Research foundation and a NARSAD Distinguished Investigator award to EBB.

## Authors’ contributions

ACK conceived the idea, designed and carried out experiments, performed analyses, provided critical intellectual input, generated and revised the paper draft; CC designed and performed omics experiments, provided critical intellectual input and revised the paper draft; CK designed and performed electroporation experiments, provided critical intellectual input and revised the paper draft; LD performed analysis and revised the paper draft; MHL & CR performed cell biology experiments; NK & DP performed the phenome wide mendelian randomization analysis and revised the paper draft; SPG designed and provided critical intellectual input for the STARR experiments; AS, DC, MLP, SS, EW, KH and KR worked on creating and analysing the ITU cohort, provided data and analysis pipelines; SR performed analysis and revised the paper draft; VS, BW, MK & SS performed experiments; MRH supported project organization and experimental procedures; ML designed and performed CRISPR-Cas9 experiments, provided critical intellectual input and revised the paper draft; SC provided critical intellectual input and revised the paper draft; EBB conceived the idea, obtained funding, supervised the study, designed experiments and analysis pipelines, provided critical intellectual input and contributed to paper draft writing and revisions.

## Supplemental Information

Supplemental Results

Supplemental Tables 1-21

Supplemental Figures 1-12

## STAR Methods

### Resource availability

#### Lead contact

Further information and requests for resources and reagents should be directed to and will be fulfilled by the lead contact, Elisabeth Binder.

#### Materials availability

Plasmids generated in this study are available upon request. Human iPSC lines used in this study are subject to MTA approvals.

#### Data and code availability

Scripts and data for the MRa-PheWAS analyses are openly available via https://osf.io/4ud6q/ for full transparency. Data for the bulk RNA-seq are deposited as BioProject under accession code PRJNA865917.

### Method details

#### iPSCs

HPS0076 Human induced pluripotent stem cells (hiPSCs) were obtained from the RIKEN BRC cell bank and reprogrammed from skin fibroblasts of a female donor (HPS0076-409b2)^61, 62^. No.1 hiPSCs were reprogrammed from NuFF3-RQ newborn foreskin feeder fibroblasts of a male donor (GSC-3404, GlobalStem)^63^. MTA approvals were obtained for the use of both hiPSC lines. hiPSCs were cultured in Matrigel-coated (1:100 diluted in DMEM-F12 (Gibco^™^, 31330-038), Corning Incorporated, 354277) Costar^®^ 6-well cell culture plates (Corning Incorporated, 3516) in mTESR1 Basal Medium (STEMCELL Technologies, 85851) supplemented with 1× mTESR1 Supplement (STEMCELL Technologies, 85852) at 37°C with 5% CO_2_. Passaging was performed with Gentle Cell Dissociation Reagent (STEMCELL Technologies, 07174). RevitaCell Supplement (1:100 diluted, Gibco^™^, A2644501) was added the day of the dissociation for 24 h to increase cell survival.

#### Cerebral organoids

Human cerebral organoids (hCOs) were created as previously described by Lancaster *et al*^17^ with some modifications. Briefly, hiPSCs were dissociated in StemPro Accutase Cell Dissociation Reagent (Life Technologies, A1110501). Nine thousand single cells were plated into Ultra-low attachment 96-well plate round bottom wells (Corning Incorporated, 7007) in human embryonic stem cell medium (hESC, DMEM/F12-GlutaMAX (Gibco^™^, 31331-028) with 20% Knockout Serum Replacement (Gibco^™^, 10828-028), 3% FBS (Fetal Bovine Serum, Gibco^™^, 16141-061), 1% non-essential amino acids (Gibco^™^, 11140-035), 0.1 mM 2-mercaptoethanol (Gibco^™^, 31350-010)) supplemented with 4 ng/ml human recombinant FGF (Fibroblast Growth Factor, Peprotech, 100-18B) and 50 µM Rock inhibitor Y27632 (Millipore, SCM075) for 4 days and in hESC medium without bFGF and Rock inhibitor for an additional 2 days to form embryoid bodies (EBs). On day 6, the medium was changed to neural induction medium (NIM, DMEM/F12 GlutaMAX supplemented with 1:100 N2 supplement (Gibco^™^, 15502-048), 1% Non-essential amino acids and 1 µg/ml Heparin (Sigma, H3149)) and cultured for an additional 6 days. On day 12, the EBs were embedded in Matrigel (Corning Incorporated, 354234) drops and transferred to 10-cm cell culture plates (TPP, 93100) in neural differentiation medium without vitamin-A (NDM-A, DMEM/F12GlutaMAX and Neurobasal (Gibco^™^, 21103-049) in ratio 1:1 supplemented with 1:100 N2 supplement 1:100 B27 without Vitamin A (Gibco^™^, 12587-010), 0.5% non-essential amino acids, insulin 2.5 µg/ml (Gibco^™^, 19278), 1:100 Antibiotic-Antimycotic (Gibco^™^, 15240-062) and 50 µM 2-mercaptoethanol) for 4 days. On day 16, hCOs were transferred onto an orbital shaker in NDM+A medium (same composition as NDM-A with the addition of B27 with Vitamin A (Gibco^™^, 17504-044) in the place of B27 without Vitamin A) and were grown in these conditions at 37°C with 5% CO_2_. NDM+A medium was changed twice per week.

#### Bulk RNA sequencing

RNA was isolated from day 45 No.1–hCOs in triplicates with 2-3 organoids pooled per replicate, either treated with 100nM dex for 7 days or veh (DMSO). Sequencing libraries were prepared using the QuantSeq 3’ mRNA Fwd kit (Lexogen) following manufacturer’s instructions. Sequencing was performed on an Illumina HiSeq4000 sequencer generating 150 bp long single-end reads. Read quality was verified using FastQC version 11.4^64^. For adapter trimming and quality filtering the software cutadapt version 1.9.1^65^ was used. For read alignment and gene quantification salmon version 0.43.1^66^ was applied setting the parameters noLengthCorrection and perTranscriptPrior to account for the tag sequencing approach. Differential gene expression was assessed using the R package DESeq2^67^. Data are openly available as BioProject with accession number PRJNA865917.

#### *In utero* electroporations of mice

All experiments and protocols were performed in accordance with the European Communities’ Council Directive 2010/63/EU and were approved by the committee for the Care and Use of Laboratory Animals of the Government of Upper Bavaria. All mice were obtained from the in-house breeding facility of the Max Planck Institute of Psychiatry and kept in group housed conditions in individually ventilated cages (IVC; 30cm x 16 cm x 16 cm; 501 cm2) serviced by a central airflow system (Tecniplast, IVC Green Line – GM500). Animals had ad libitum access to water (tap water) and standard chow and were maintained under constant environmental conditions (12:12 hr light/dark cycle, 23 ± 2 °C and humidity of 55%). Time-pregnant female mice at stage E13.5 or E14.5 were anesthetized by intraperitoneal injection of saline solution containing fentanyl (0.05 mg per kg body weight), midazolam (5 mg per kg body weight), and medetomidine (0.5 mg per kg body weight) and embryos were electroporated as described in Nature Protocols by Saito *et al* ^68^. In brief, plasmids were mixed with Fast Green (2.5 mg/μL; Sigma F7252) and 1-2*μ*l were injected at a final concentration of 1*μ*g/μL using glass micropipettes (5-000-1001-X10, Drummond Scientific). The DNA was electroporated into the cells by delivering 5 pulses applied at 40V for 50ms in 1sec intervals. Anesthesia was terminated by injection of buprenorphine (0.1 mg per kg body weight), atipamezole (2.5 mg per kg body weight), and flumazenil (0.5 mg per kg body weight). The pups remained *in utero* for 3 or 6 more days, after which euthanizing occurred and the pups’ brains were isolated. Brains were fixed at 6dpe (days post electroporation) in 4% PFA (paraformaldehyde) for 16 h, cryo-preserved with 30% sucrose for at least 16 h and stored at −20 °C in OCT (optimal cutting temperature, Thermo Fischer Scientific, 23-730-571). For immunofluorescence, 12 µm cryosections were prepared on SuperFrost^TM^ slides. For the cell cycle re-entry experiment, embryos were electroporated at E13.5 and the mother received an intraperitoneal injection of 14mg/ 100g of body weight of BRDU diluted in saline at E15.5, 24 hours before sacrifice at E16.5.

#### Electroporations of human cerebral organoids

HPS0076-hCOs were electroporated as described in^69, 70^. In brief, hCOs were kept in antibiotic-free NDM+A medium for three h prior to electroporation. Electroporation was performed in hCOs at day 43 after the initial plating of the cells, and fixed at 7dpe. During electroporation, hCOs were placed in an electroporation chamber (Harvard Apparatus, Holliston, MA, USA) under a stereoscope. Using a glass micropipette (5-000-1001-X10, Drummond Scientific) 0.5 µL of plasmid DNA was injected together with Fast Green into different ventricles at a final concentration of 1*μ*g/μL. hCOs were subsequently electroporated with 5 pulses applied at 80 V for 50 ms each, at 500 ms intervals (ECM830, Harvard Apparatus). Following electroporation, hCOs were kept for an additional 24 h in antibiotics-free NDM+A media and then changed into the normal NDM+A media until fixation at 7dpe. hCOs were fixed using 4% PFA for 1 hour at 4 °C, cryo-preserved with 30% sucrose for 16 h and stored at −20 °C in OCT. For immunofluorescence, 16 µm cryosections were prepared on SuperFrost^TM^ slides.

#### Glucocorticoid treatment in cerebral organoids and mice

*hCOs*: Day 43 hCOs of both lines were treated for 7 days (chronic treatment) with 100nM of dexamethasone (dex). To achieve the concentration used, dex was diluted in DMSO (dimethyl sulfoxide) in a concentration of 100μM and subsequently diluted in NDM+A culture medium to a final concentration of 100nM. The DMSO control (vehicle-veh) underwent the same dilutions as described for dex. The medium was changed every two days. At the end of the treatment, hCOs were fixed using 4% PFA for 1 hour at 4 °C, cryo-preserved with 30% sucrose for 16 h and stored at −20 °C in OCT. For immunofluorescence, 16 µm cryosections were prepared on SuperFrost^TM^ slides.

*Fetal mice*: We injected fetal mouse brains at E13.5 (embryonic day 13.5) with dex diluted in DMSO and analysed the effects at E16.5. To achieve ∼100nM of dex concentration we measured the volume of E13.5 mouse brains by measuring the volume displaced when submerged in saline, to ∼25μl. Thus, we injected 1μl of 2.5μM dex or veh into an E13.5 mouse brain ventricle to achieve a final concentration of ∼100nM.

*Concentration and timing rationale:* The concentration of 100nM was chosen to mimic the antenatal corticosteroid therapy scheme used during pregnancy. The guidelines for sGC use are four doses of 6mg dex every 12 h given intramuscularly^8, 71^ when used in at risk for preterm birth pregnancies. A single dose of 6mg reaches a C_max_ of 65-95ng/μl at T_max_ 3 h^71^ which equals to a concentration of ∼162nM-245nM, whereas 1.5mg would equal to ∼40.4nM-61.25nM. The maternal to fetal steroid hormones ratio has been reported anywhere from 0.4 and higher, days after the treatment^71^. So, from a single 6mg dose we would expect at the very least ∼64.8nM-98nM reaching the fetus after 3 h post treatment. Similarly, from 1.5mg given each day we would expect at least ∼16.16nM-39.2nM reaching the fetus after 3 h post treatment. A previously published study that used 100nM of androgens, which are also steroid hormones, in hCOs measured an actual concentration of 16nM^72^. They attributed this phenomenon to the fact that steroid hormones have high affinity to plastic due to their lipophilic nature^73^. Since our primary aim was to study the role of sGC therapy schema, the use 100nM of dex should parallel the amounts reaching the embryo by the clinically used concentrations. As mentioned the schema of antenatal corticosteroids treatment is four doses during a two-days’ time. Dexamethasone is a long-acting corticosteroid with a biological half-life of 36 to 54/72 hours^74^. Considering the active administration time and the biological half-life of the steroid, we would expect dex effects on the fetus for 5-7 days, with peak exposure for 2-3 days and then a decrease. Thus, we chose to administer dex for 7 days.

#### Immunofluorescence

Sections were post-fixed in 4% PFA for 10 min at RT and permeabilized with 0.3% Triton in PBS for 5 min. Sections were subsequently blocked with 0.1% TWEEN, 10% Fetal Bovine Serum and 3% BSA. Primary and secondary antibodies (see Table S18) were diluted in blocking solution and nuclei were visualized using 0.5 µg/mL 4,6-diamidino-2-phenylindole (DAPI, Sigma Aldrich). Cover slips were mounted with Aqua Poly/Mount (Polysciences, 18606-20). For PAX6, EOMES, BCL11B, ZBTB16, SOX2, Ki-67, TBR1 and SATB2 antigen retrieval was performed before fixing with PFA. Briefly, sections were incubated with citric buffer (0.01 M, pH 6.0) for 1 min at 720 Watt and 10 min at 120 Watt, left to cool down at RT for 20 min and washed three times with PBS for 5 min.

#### Protein isolation and Western blot

Proteins were isolated on ice in RIPA buffer (Sigma-Aldrich, R0278) supplemented with protease (Sigma Aldrich, P8340) and phosphatase inhibitors (Sigma Aldrich, 4906845001). For each experiment three biological replicates were included each bearing the homogenate of three individual hCOs. 30µg of protein extracts were separated by SDS-PAGE with an 8% gel. Proteins were transferred to a PVDF membrane (Millipore, IPVH00010). For detection, membranes were incubated with primary antibodies (Table S18) for 16 h at 4°C and with horse-radish peroxidase-labeled secondary antibodies (Table S18) at RT for 1 hour. Subsequently, they were treated with Immobilon Western HRP Substrate Luminol Reagent and Solution (Millipore, WBKLS0500) to visualize the bands. The quantification was performed in the Bio-Rad Image Lab Software (Version 6.1). Relative protein expression levels were quantified and normalized with ACTIN as endogenous control.

#### RNA isolation and quantitative PCR

Total RNA was extracted from hCOs using the RNeasy Mini extraction kit (Qiagen, 74104) according to the manufacturer’s instructions. For each experiment three biological replicates were included each bearing the homogenate of three individual hCOs. Complementary DNA (cDNA) synthesis was performed using the Maxima H Minus Reverse Transcriptase (Thermo Fisher, EP0751) with oligo(dT)16 primers (Invitrogen, N8080128) and random hexamers (IDT DNA Technologies, 51-01-18-25) in a 1:1 ratio. Quantitative PCR (RT-qPCR) reactions were run in quadruplicate using PrimeTime qPCR Primer Assays (IDT DNA Technologies, Table S19) and PrimeTime® Gene Expression Master Mix (IDT DNA Technologies, 1055770) on a LightCycler 480 Instrument II (Roche). Relative gene expression levels were quantified using the relative quantification method and normalized with POLR2A and YWHAZ as endogenous control genes. Data shown are additionally normalized over the vehicle samples so that vehicle samples have values of approximately one.

*Dexamethasone effects on fetal mice*: For the effect of dex on ZBTB16 transcription at least five different fetal mouse brains were analyzed per condition. After brain isolation both cortices were excised, RNA was isolated and quantitative PCR was performed as already described.

#### Plasmids preparation

Multiple PCR inserts were simultaneously cloned by In-Fusion HD Cloning Plus into the linearized vector pCAG-DsRed2 (Addgene, #15777) to create the pCAG-ZBTB16-F2A-GFP plasmid. More specifically, the human ZBTB16 ORF (NM_006006.5, 2034bp) sequence was amplified from a plasmid delivered from GenScript and the F2A-GFP from the Snap25-LSL-2A-GFP vector (Addgene, #61575). PCR primers were designed for the sequence of interest with extensions that are complementary to the ends of the linearized vector or the corresponding fragment (Table S19). PCR was performed using the CloneAmp HiFi PCR Premix (Takara Bio, 639298) following manufacturer’s instructions

After cloning the fragments for 3 h at 37°C, the new construct was transformed into StellarTM competent cells and grown for 16 h on agar plates containing 100μg/ml ampicillin. Single colonies were picked, plasmidic DNA was isolated with Qiagen plasmid kits (12123, 12143) and the genotype was checked with Sanger sequencing (Table S19). The pCAG-F2A-GFP plasmid was created by cutting the ZBTB16 fragment out of the pCAG-ZBTB16-F2A-GFP plasmid using the BamhI and BglI restriction enzymes.

#### Luciferase reporter assays

Luciferase assays were designed to assess the activity of the three human *PAX6* promoters^75^, P0, P1 and Pa (Table S20) under ZBTB16 overexpression. The promoter sequences were cloned into the firefly luciferase (Luc2) reporter expression pRP vector by VectorBuilder. The human ZBTB16 expression plasmid was generated by VectorBuilder using the pRP backbone. 500ng of total plasmid DNA (75% of human ZBTB16 expression plasmid, 15% of reporter plasmid and 10% of the pCAG-F2A-GFP, as internal control of transfection efficiency) were transfected into 72,000 HeLa cells in a well of a 24 well plate using Lipofectamine 2000 (ThermoFisher Scientific) following manufacturer’s instructions. All transfections were carried out in triplicates. HeLa cells were cultured in DMEM-F12 medium (Gibco™, 11320033) supplemented with 10% FBS (Gibco™, 16141-061) and 1× Antibiotic-Antimycotic (Gibco™, 15240-062). The medium was refreshed the next day and 48 h later transfected cells were PBS washed and incubated for 15 min at RT in 1× passive lysis buffer (Biotium, 99821). Plates were kept at least 1 hour at –80°C. Next, the lysate was scraped and centrifuged at full speed for 30 sec at 4°C. 20μl of the supernatant was subjected to the luciferase assay with the addition of 50 ul D-luciferine (Beetle Juice luciferase assay, PJK, 102511-1) by using a Tristar multimode reader (Berthold). The luminescence measurement was done for 5 sec using 2 sec delay. In addition, 50μl of the lysate were assessed for GFP fluorescence. The luciferase reading was normalized over the GFP results for each well. Data is shown as fold changes over the control plasmid.

#### CRISPR-Cas9 editions of hiPSCs

CRISPR-Cas9 editing was used to create genomic deletions of the *ZBTB16* exon 2 and of approximately 400 bp of the regulatory element centered on rs648044 an intronic *ZBTB16* variant. Genome editing was done by electroporation of gRNA pairs (crRNA/tracrRNA duplexes, alt-CRISPR IDT) and recombinant S.P. HiFi Cas9 V3 nuclease (IDT DNA Technologies, 1081060). crRNAs were designed using the Benchling webtool and analysed for self– or heterodimers using the IDT OligoAnalyzer™ tool (Table S21). To delete the region of interest, 300,000 No.1 iPSCs were transfected with 35pmol of each gRNA (crRNA/tracrRNA duplex 1:1 in 1× Arci annealing buffer (STEMCELL Technologies, 76020), IDT DNA Technologies), 40pmol of Cas9 and 100pmol of electroporation enhancer (IDT DNA Technologies, 1075915) in 26.57μl of the P3 primary cell 4D_X Kit S (Lonza, V4XP-3032) using the 4D-Nucleofector X Unit (Lonza, AAF-1003X) with the CA-137 program. Edited cells were plated into one well of 24-well plate coated with Matrigel 1:100 and cultured in supplemented mTESR1 Basal Medium and, for the day of the edition, with RevitaCell (1:100)) at 37°C with 5% CO_2_. For control editions, cells were electroporated with Cas9 without the addition of gRNAs. The next day the medium was changed to supplemented mTESR1 and cells were propagated approximately for 2-3 days till they reached 80-90% confluence. Next, cells were passaged into a well of a 6-well plate coated with Matrigel using Gentle Cell Dissociation Reagent and propagated until confluence. Subsequently, 89.5% of the cells were expanded, 10% of them were taken for bulk genotyping analysis and 0.5% were plated in a well of a 6-well plate coated with Matrigel to generate single-cell-derived clonal cell lines. Bulk and single cell DNA extraction was done using 30µl of QuickExtract DNA Extraction Solution (Lucigen, QE09050). Briefly, cells were dissociated, pelleted, resuspended in the extraction solution and incubated at 65 °C for 10 min and 98 °C for 5 min. PCR was performed using the primers in Table S19, the Q5 high fidelity master mix (New England Biolabs, M0494S) and 40ng of cell extract in a total volume of 10 µl. The thermal cycling profile of the PCR was: 98 °C 30 s; 35 × (98° 10 s, 65 °C 15 s, 72 °C 60 s); 72 °C 2 min. Automated electrophoresis technique (DNA screen tape analysis, Agilent) and Sanger sequencing (Eurofins, primers in Table S19) were used to confirm the presence of CRISPR-Cas9-mediated knockout mutants in the bulk population and the single clones. For the *ZBTB16* exon 2 edition we selected a heterozygous KO cell line (termed *ZBTB16*^+/-^ and *ZBTB16*^+/+^ for the control edition) showing by western-blot analysis a ∼46% reduction in protein expression (Table S18). The rs648044 is a heterozygous SNP in the iPSC cell line used (rs648044^G/A^). From the edited single clones, we selected a heterozygous KO cell line of the enhancer element harboring the A genotype (termed rs648044^G/-^ and rs648044^G/A^ for the control edition). The effect of the KO on *ZBTB16* expression was assessed by RT-qPCR in veh and dex conditions and using *POLR2A* and *YWHAZ* for normalization (Table S19).

#### Flow Cytometry

No.1-ZBTB16^+/+^ and ZBTB16^+/-^ hCOs were collected for Flow Cytometry analysis (FCa) at day 50 after 7 days of treatment with veh or 100nM dex and/or 1μM of the GR antagonist RU-486 (Selleck, S2606). Three to four samples per batch were analysed and each sample contained two individual hCOs. hCOs were enzymatically dissociated with accutase supplemented with DNase I (ThermoFisher Scientific, EN0521) at 37°C for maximum 40 min. During incubation, every 10 min the hCOs were additionally manually dissociated with a P1000 pipette. Once dissociated, the hCOs were centrifuged for 5 min at 300g and the pellet was resuspended in PBS. Next, cells were centrifuged and the cell pellets were fixed with 70% EtOH at –20°C for one hour. Subsequently, After the addition of 5ml washing buffer (PBS + 1% FBS) fixed cells were centrifuged for 30 min, at 4°C and 500g. The cell pellet was resuspended in 200μl staining solution (wash buffer supplemented with anti-PAX6 and anti-EOMES, Table S18) and incubated for 30 min on ice. After the primary antibody incubation, 1ml of washing buffer was added and the stained cells were centrifuged for 30 min, at 4°C, at 500g. The cell pellet was resuspended in 200μl secondary antibody staining solution (wash buffer supplemented with anti-rabbit 488, anti-sheep 594 and DAPI, Table S18) and incubated for 30 min on ice. The stained cells were filtered through an 100μm cell strainer and diluted in additional 200μl wash buffer. FCa was performed at a FACS Melody (BD) in BD FACS Flow TM medium, with a nozzle diameter of 100 µm. For each run, 20,000 cells were analyzed. For the 488 fluorophore we used the 488nm laser coupled with the 530/30 filter and for the 594 fluorophore we used the 561nm laser coupled with the 613/18 filter. *Gating strategy*: SSC-A/FSC-A gates were used to exclude cell debris and FSC-H/ FSC-W to collect single cells. Gating for fluorophores was done using samples stained with secondary antibody only. The flow rate was set below 30 events/s. Further analysis was done using the software of the FACS Melody (BD) and the online free software Floreada.io.

#### Targeted Bisulfite Sequencing

Targeted bisulfite sequencing was performed following the original protocol^40^. DNA was isolated from day 50 hCOs that were treated with 100nM dex or veh for 7 days with the NucleoSpin Genomic DNA kit (Macherey-Nagel). Bisulfite treatments were performed in triplicate for 200ng of DNA from each sample with the EZ DNA Methylation Kit (Zymo Research, D5001) and then pooled to run one PCR amplification per amplicon in order to reduce cost and maximize the number of samples per sequencing run. Twenty nanograms of bisulfite-converted DNA and 49 amplification cycles were then used for each PCR amplification (Table S9) with the Takara EpiTaq HS Polymerase (Clontech, R110A). PCR amplicons were then quantified with an automated electrophoresis technique (2200 DNA screen tape analysis, Agilent) and pooled in equimolar quantities for each sample. AMPure XP beads (Beckman Coulter, A63880) were used for a double size selection (200–500 bp) to remove primer dimers and high molecular DNA fragments. Libraries were generated using the TruSeq DNA PCR-Free HT Library Prep Kit (Illumina, 20015963) according to the manufacturer’s instructions. Each library was quantified with the Qubit® 1.0 (ThermoFisher Scientific), normalized to 4 nM and pooled. Library concentration and fragment sizes were checked with the Agilent’s 2100 Bioanalyzer (Agilent Technologies) and quantitative PCR using the Kapa HIFI Library quantification kit (Kapa Biosystems, KK4824). Paired-end sequencing was performed on an Illumina MiSeq Instrument (Illumina) with their MiSeq Reagent Kit v3 (MS-102-3001, 2Å∼ 300 cycles) with the addition of 15% of PhiX Control v3 library (Illumina, FC-110-3001) generating 300bp long paired-end reads. Reads were processed as described by Roeh *et al*.^40^. In brief, read quality was verified using FastQC^64^, and cutadapt v1.11^76^ was applied to trim reads. Subsequently, reads were aligned to a restricted reference consisting of the amplicon sites using Bismark v0.18.2^77^. Paired-end reads were stitched together using an in-house perl script. Using the R package methylKit v.1.6.3^78^ increasing Phred score quality cutoff to 30, methylation levels were extracted. Further filtering was conducted in R. We excluded artifacts on a per sample basis, including low-coverage amplicons (sequencing coverage < 1000, 0 samples excluded) and samples with bisulfite conversion efficiency lower 95% (0 samples excluded). To test for significance, individual CpGs of the same enhancer element were tested with two-way ANOVA and corrected for multiple comparisons with the two-stage step-up method of Benjamini, Krieger and Yekutieli^79^.

#### Methylation analysis by Bisulfite Pyrosequencing

DNA was isolated from day 50 hCOs that were treated with 100nM dex or veh for 7 days with the NucleoSpin Genomic DNA kit (Macherey-Nagel). Bisulfite treatments were performed in triplicate for 300ng of DNA from each sample with the EZ DNA Methylation Kit (Zymo Research, D5001) and then pooled to run one PCR amplification per amplicon in order to reduce cost and maximize the number of samples per sequencing run. Twenty nanograms of bisulfite-converted DNA and 45 amplification cycles were then used for each PCR amplification (Table S9) with the PyroMark PCR Kit (Qiagen, 978703). Pyrosequencing primers were designed with the MethMarker software and carried out on a PyroMark Q48 Autoprep using PyroMark Q48 Magnetic Beads (cat. no. 974203), PyroMark Q48 Discs (cat. no. 974901) and PyroMark Q48 Absorber Strips (cat.no. 974912), according to the manufacturer’s recommendations. To test for significance, individual CpGs were tested with two-way ANOVA and corrected for multiple comparisons with the two-stage step-up method of Benjamini, Krieger and Yekutieli^79^.

#### HPS0076 chronic single-cell dataset

We generated and analyzed a 10×3’ v2 scRNA-seq of 70 days old cerebral organoids (HPS0076 derived) treated with 100nM of dexamethasone for 10 days prior to collection dataset to validate our findings using single-cell transcriptomics. This dataset contained 3794 control cells and 8539 treated cells. We used the MAST R package to compute differential expression between treatment and control for each of the 8 cell-types in the dataset. For this, we only used cells from samples with more than 10 cells in the dataset. We furthermore removed any genes which were expressed in less than 5% of the cells and controlled for the number of expressed genes in the expression model. We used the likelihood ratio test to compute the statistical significance of differential expression and corrected for multiple testing using FDR correction. We adhered to the filtering and testing procedure when we computed differentially expressed genes between ZBTB16 positive and negative cells in the combined population of vehicle-treated Radial Glia and Intermediate Progenitor cells. For the differential testing procedure between treatment and control in cells positive for PAX6 and EOMES, we did not apply the filtering of samples which do not contain more than 10 cells due to low cell numbers in this case.

For Gene Set Enrichment Analysis, we used the R package Cluster Profiler (release 3.17)^80, 81^ and plotted the results using EnrichPlot^82^.

#### No1 acute single-cell dataset

Uniform Manifold Approximation and Projection plots (UMAP)^83^ were used to visualize the expression of five genes (*ZBTB16, PAX6, TSC22D3, KLF9, HEYL*) in a previously published single-cell RNA seq dataset (Cruceanu et al., AJP, 2022^18^). Specifically, gene expression was plotted in the day 30 subsets from cell-line 1 of the aforementioned dataset using the SCANPY^84^ python package. The software environment and data processing used to produce these Figure s was identical to the environment and processing steps used to produce expression-UMAPs in Cruceanu et al., AJP, 2022^18^. Using the same data subset, the number of cells positive (>0 raw counts) and negative (0 raw counts) for the five genes were computed across the cell-type clusters defined in the associated publication. The percentages of the positive, for each gene, cells in each cluster depicted in Figure 1j were calculated as a fraction of the total cells positive for each gene in all clusters using the Wilson/Brown method with 95% confidence intervals.

#### STARR (Self-Transcribing Active Regulatory Region sequencing)-qPCR

*rs648044 cloning into STARR reporter plasmid:* For the STARR-qPCR assay, candidate sequences are cloned in the 3’-UTR of the sgGFP reporter gene driven by the SCP1 minimal promoter^85^. Once transfected into GR expressing cells, active regulatory elements would modulate GFP expression under dex treatment^86^. 201bp long DNA inserts (gblock, IDT) containing 200 bp putative regulatory element centered on the rs648044 (reference and alternative allele) and flanked by 15bp sequence homologous to the STARR reporter construct (Table S19) were inserted by in-Fusion HD Cloning Plus into the human STARR-seq vector digested with SalI and AgeI following the manufacturer’s instructions (Takara Bio, 102518). The inserts had additionally 2bp to reconstitute the AgeI and SalI restriction sites lost during cloning. Subsequently, the constructs were transformed into Stellar^TM^ competent cells (Takara Bio, 636763) and grown for 16 h on agar plates containing 100μg/ml ampicillin. Single colonies were picked, plasmidic DNA was isolated with Qiagen plasmid kits (12123, 12143) and the genotype was checked with Sanger sequencing (Table S19).

*U2OS-GR18 cells transfection:* U2OS cells stably transfected with rat GRα (GR18 cells)^87^ were cultured in Dulbecco’s Modified Eagle Medium-high glucose supplemented (Gibco^TM^, 11965084) with 10% FBS (Gibco^™^, 16141-061) and 1× Antibiotic-Antimycotic (Gibco™, 15240-062). Two million cells were transfected with 2μg of plasmid in triplicates using the Amaxa Nucleofector II Kit V and program X-001 (Lonza Bioscience). After 16 h cells were treated with 100nM dex or veh for four h. RNA was isolated with the RNeasy Mini extraction kit (Qiagen, 74104) according to the manufacturer’s instructions.

*cDNA conversion and qPCR:* cDNA was generated using two gene specific primers for plasmid GFP and RPL19 as endogenous control (Table S19) and the Quantitect Reverse Transcriptase (Qiagen) kit following manufacturer’s instructions. Regulatory elements activity was assessed with qPCR using primers for RPL19 and GFP (Table S19). The qPCR was analysed with the relative quantification method and GFP expression was normalized over the RPL19 expression. Data are shown both as expression values (2^-DCt^) and as fold changes of the expression values (dex/veh).

#### Mendelian randomization analysis (MRa) and Phenome-wide association (PheWAS) study

All processing and analysis were conducted using R software^88^. To assess the potentially causal effect of ZBTB16 on a range of phenotypes, rs648044 was used as genetic proxy with log fold-changes of the effect allele (A), averaged over STARR-qPCR experiments, (β= 1.475794, SE= 0.1015) used as exposure effect and variance estimates.

*Outcome phenotype selection:* Outcome phenotypes were selected from the MRC IEU OpenGWAS platform focusing on phenotype batches originating from the UK Biobank study, the NHGRI-EBI GWAS Catalog, and a GWAS on brain imaging phenotypes based on UK Biobank data (Table S13)^89–92^. From the initial phenotype list originating from these batches, 8979 phenotypes from GWAS of European populations were selected after filtering out duplicates and phenotypes not of interest to this study (e.g., *“*Patient Care Technician r*esponsible for patient data” or “Day-of-week questionnaire completion requested”).* Duplicates were filtered using a semi-automated procedure including deletion of phenotypes with identical names and smaller GWAS sample size as well as manual filtering of phenotypes with high similarity in trait names (quantified using the restricted Damerau-Levenshtein distance >0.8 implemented in the stringdist package)^93^. This procedure resulted in a final phenotype list of 7,503 phenotypes.

*Mendelian randomisation analysis (MRa):* The *TwoSampleMR* package was used for MRa^94^. Outcome data for rs648044 were extracted from phenotype summary data, which were available for 7,323 outcomes, and effect and reference alleles were harmonised with exposure data. Wald ratio MRa estimation was applied as method of choice for single-SNP MRa for all remaining outcome phenotypes^95^. To account for the multiple comparisons, P-values from all 7,323 comparisons were corrected using the Benjamini-Hochberg method^96^. In results visualizations, we also provide the Bonferroni-corrected multiple comparison threshold as reference^97^.

*Illustration of PheWAS associations*: Significant phenome-wide associations with brain region phenotypes were illustrated by overlaying human brain atlas regions as proxies of the regions of interest onto the MNI template in MRIcroGl (version <u>v2.1.58-0</u>, https://www.nitrc.org/projects/mricrogl/). Due to the different analysis streams and granularities of the significant brain phenotypes, different atlases were used to portray the results. For illustration purposes, regions taken from probabilistic atlases were thresholded at 10%. As a proxy for the circular anterior insular cortex thickness, we used the anterior insula of the “Hammersmith atlas”^98^. Medial lemniscus, cingulate gyrus part of cingulum, cingulum hippocampus, uncinate fasciculus, posterior limb of internal capsule, inferior fronto-occipital fasciculus, uncinate fasciculus, anterior corona radiata and the corticospinal tract were portrayed using the JHU ICBM DTI 81 white matter labels^99^, the acoustic radiation was visualized using the Juelich histological atlas^99^, the inferior longitudinal fasciculus, and the thalamic radiation by the JHU White Matter Tractography^100^ and the superior thalamic radiation by the XTRACT HCP Probabilistic tract atlas^101^.

Scripts and data for the MRa-PheWAS analyses are openly available via https://osf.io/4ud6q/ for full transparency.

#### Intra-Uterine prospective pregnancy cohort study

*Study Cohort:* The Intra-Uterine sampling in early pregnancy (ITU) is a prospective pregnancy cohort study with the overarching aim to unravel maternal-placental-fetal mechanisms involved in the programming of health and disease^52^. It comprises 943 women and their singleton children born alive in Finland between 2012-2017. The women were recruited at the voluntary national 21-trisomy screen, offered to all pregnant women at gestational weeks 9 to 21. Of these women, 543 (57.6%) were referred for fetal chromosomal testing at Helsinki and Uusimaa Hospital District Fetomaternal Medical Center (FMC) and thereafter cleared for fetal chromosomal abnormality. The rest, 400 (42.4%) women, had a negative 21-trisomy screen result and were not referred for fetal chromosomal testing.

*Ethics Approval:* The ITU study was conducted according to the World Medical Association’s Declaration of Helsinki and the research protocol was approved by the ethics committee of the Helsinki and Uusimaa Hospital District (approval date: 06.01.2015, reference number: 269/13/03/00/09). All women provided written informed consent. In compliance with the General Data Protection Regulation of the European Union, the personal data of all participants were de-identified, protected at all times, and confidentiality agreements were signed by all personnel with ITU data access.

*Data availability:* A complete set of salivary cortisol, fetal rs648044 genotype (genotype frequencies of AA = 0.234, GG = 0.284, AG = 0.482, Hardy-Weinberg-Equilibrium [HWE] p = .630), and child neuropsychological assessment at 35.6 months of age (range 24.5 to 42.5) was available for 246 mother-child dyads (46.7% female sex) before and for 221 mother-child dyads (49.1% female sex) after gestational week 28.

*Salivary Cortisol Assessment:* Salivary cortisol sample collection, storage, and competitive enzyme immunoassay were previously described^52^. Briefly, mothers collected saliva in early (<22 weeks, T1), mid-(22-35 weeks, T2), and late pregnancy (≥36 weeks, T3) when waking up (S1), 15 (S2) and 30 min (S3) thereafter; at 10 am (S4), noon (S5), 5 pm (S6), and when going to sleep (“lights out”, S7). Gestational week at assessment was computed using the ultrasound-based date of conception and self-reported date of cortisol assessment.

Salivary cortisol samples were available for 690 women (307 with and 383 without chromosomal testing) of which 15 were excluded due to corticosteroid treatment^102^ resulting in a total of 9,992 (70.49%) out of the maximum number of 14,175 samples in 675 women. Samples measured in duplicate were averaged and those with a coefficient of variation greater than 0.25 (1.6%, 164/9,992) excluded. Cortisol values below the lower limit of the assay range were truncated at 0.05 µg/L (2.4%, 240/9,992, no values fell above the upper limit). For some women, multiple assessment days (i.e., 24) fell into the same pregnancy stage and only the one with most complete observations, protocol adherence, or absence of day-specific outliers (i.e., >2SD of the day-specific mean) was kept, leading to the exclusion of 159 samples. For 6 women, date records were missing, and their 89 samples were excluded. Further exclusion criteria were self-reported illness on the day of sampling (89 samples, leading to the exclusion of all samples of 2 women), getting up before the first sample (108 samples), sampling of S2 or S3 more than 60 min after awakening (10 samples), or S4 sampling within the first 60 min after awakening (17 samples). Missing sampling times (0.1%, 19/9,992) were imputed according to sampling protocol for S2 to S6 or sample median for S1 and S7. The distribution of the cortisol samples was positively skewed, and thus natural log+1 transformed. To further reduce skewness, outliers (i.e., >4 SD pregnancy stage-specific mean) were winsorized (9 samples).

The resulting dataset consisted of 9,356 samples from 667 women (292 with and 375 without chromosomal testing). Of these, 589 provided salivary samples before gestational week 28 (range 12.3 to 27.9 weeks) and 499 after gestational week 28 (range 28.14 to 41.3 weeks). Total cortisol output per pregnancy stage was estimated based on the area under the curve with respect to ground (AUCg) of all seven saliva samples using the trapezoid rule^103^. When multiple occasions fell into the same gestational interval (i.e., gestational week <28 or >28 weeks), the total cortisol output and gestational week at assessment were averaged.

*Fetal Genotypes:* DNA was extracted from cord blood leucocytes using a bead-based method optimised by tissue type (Chemagic 360, Perkin Elmer) and genotyping performed on Illumina GSA-24v2-0 A1 arrays according to the manufacturer’s guidelines. Genotyping quality control and imputation were previously described in Kvist et al., 2022^52^. In total, the rs648044 genotype was available for 446 (genotype frequencies of AA = 0.226, GG = 0.285, AG = 0.489, HWE p = .704) out of a maximum of 944 samples.

*Child Neuropsychological Assessment*: At 35.48 months (range 24.5 to 42.5 months) follow-up, children’s cognitive skills were assessed by trained psychology students (supervised by a clinical pediatric neuropsychologist) using the Bayley-III screening test^53^. The Bayley-III is a norm-referenced test designed to identify infants and toddlers at risk for developmental delay. Scores were calculated using normative data tailored for exact age, with higher scores indicating better performance. The Bayley cognitive assessment was available for n = 618 (50.5% female) out of a maximum of 944 children. Neurodevelopmental delay was defined as 1SD below the sample mean.

*Maternal and child covariates:* All mothers consented to link their data to Finland’s detailed nationwide registers. Maternal sociodemographic and pregnancy characteristics used in this study included maternal age at delivery [years], parity [nulli-/multiparous], whether the mother was referred for fetal chromosomal testing [yes vs no], maternal education [primary/applied university/university education], which was self-reported in early pregnancy, and pre-pregnancy Body Mass Index [BMI, weight (kg)/height^2 (m^2)] verified by measurement in the first antenatal clinic visit between 7-10 gestational weeks. Child characteristics included gestational age at delivery and child sex.

*Logistic regressions: Binomial* logistic regressions were run using the “glm” function of the “stats” package in R (version 3.6.1)^88^. We tested the association of the interaction effect of mean cortisol * rs648044 A allele on delay in cognitive performance as defined by the Bayley cognitive subscale, using the following covariates: gestational week at cortisol assessment, child sex, child age when the Bayley questionnaire was done, gestational age at birth, maternal age and education, parity, maternal body mass index and case vs control as women were either recruited at routine care (controls) or after referral for chromosomal testing (cases). Cases all tested negative for chromosomal abnormality). The plot on Figure 6e was done using the “ggplot2” package^104^ in R.

### Quantification and Statistical analysis

#### Image analysis and Quantifications

Immunostained fluorescent staining was visualized using a Leica laser-scanning microscope and analyzed with FIJI ((Fiji Is Just) ImageJ 2. 1. 0/1.53c; Java 1. 8. 0_172[64 bit])^105^. For analysis of the hCOs, we included ventricles that fulfilled the following criteria: clear ventricular structure with elongated, radially-organized cells surrounding the ventricular zone (VZ-determined with DAPI staining), at least one cell electroporated in the VZ (for electroporation experiments) and expression of PAX6 and EOMES to define dorsal cortical ventricles. Cell counting was performed in one representative plane of a z stack using the cell counter tool in FIJI. For electroporations binning analysis was done.

*Electroporations on hCOs*: For each experiment, at least seven independent ventricles from five different HPS0076-hCOs generated in three independent preparations grown in different times were analyzed. Analysis of the ZBTB16 phenotype in the electroporation experiments was performed by always comparing hCOs electroporated with ZBTB16-F2A-GFP plasmid vs F2A-GFP control plasmid in the same batch. Throughout the area of the electroporation, bins were set as follows: the maximal distance between the most migrated GFP-positive cell and the apical surface of the ventricle with the first GFP-positive cells was measured and divided into three equally-heighted bins. Bin A is mainly comprised of the VZ, bin B of the outer-most basal part of the VZ and the SVZ and bin C of the CP. As normalization we used the number of GFP-positive cells in total or per bin as specified in each section.

*Dexamethasone effects on hCOs*: For each experiment, at least twelve independent ventricles from six different HPS0076-hCOs and at least five independent ventricles from three different No.1-hCOs, generated in two independent preparations for Line-1 and in one preparation from Line-2 iPSCs grown in different times were analyzed. Analysis of the dex phenotype was performed by always comparing hCOs grown in parallel in the same batch. For analysis of the dex effects in hCOs, VZ and SVZ (subventricular like-zone) were defined by the cell shape and proximity to the apical zone. The VZ area presented elongated, radially-organized cells positive for radial glia markers (PAX6, SOX2) but not for basal progenitors’ markers (EOMES). The area on top, assigned as SVZ, was positive for basal progenitor markers. Areas were defined and measured in FIJI using the ROI Manager tool. As normalization we used the measured area surface.

*In utero electroporations of fetal mice*: For each experiment and condition, at least eight mouse cortical sections from five different embryos collected from two littermate mothers were analyzed. For analysis of the *in utero* electroporations in mice, we used binning analysis. We chose a cortical column that had the majority of electroporated cells and we set the bins as follows: the maximal distance between the most migrated GFP-positive cell in the cortical plate and the apical surface of the ventricle was measured and divided by five. The width of the bin was the width of the 40× lens image and it was the same for all sections and mice. As normalization we used the number of GFP-positive cells in total or per bin as specified in each section.

#### Statistics & Plots

The statistical analysis was performed in GraphPad Prism (Version 9. 1.0 (2021)). Datasets were tested for normality with a D’Agostino & Pearson K2 test^106^. Groups were then compared with a two-tailed Mann-Whitney test or a one-way or a two-way ANOVA with multiple comparisons corrected with the two-stage step-up method of Benjamini, Krieger and Yekutieli^79^, according to the type of data and their distribution. More specifically, p-values for Figures: 1f, 3e&f, 4c,e&g, 5c,e&i, 6b and for Supplemental Figures: 1a&b, 3c&e, 7c, 9b, 10a,b,e,g,j&l, 11b, 12a,d&e were calculated using two-way ANOVA with Benjamini, Krieger and Yekutieli multiple testing correction. P-value for Figure 2f was calculated using one-way ANOVA with Benjamini, Krieger and Yekutieli multiple testing correction. P-values for Figures: 1c, 2e, 3c, 5h&i and for Supplemental Figures: 3b,d&e, 5b&d, 8b&d, 10d,i&k, 12b were calculated using Mann–Whitney (two-tailed) comparison between the two treatment/electroporation groups (veh-dex or ZBTB16 overexpression plasmid-control plasmid). On the plots, the p-values depicted are either the Welch’s t-test p-value, the Mann-Whitney comparison p-value or for the ANOVAS the corrected post-hoc p-values. For each statistical test the N equals the dots depicted in each plot.

Dots in plots of Figures: 1c&f, 3c,e&f and of Supplemental Figures: 1a&b, 5d, 7c, 8b&d represent individual ventricles. Specifically, for HPS0076 hCOs results in Figure 1c&f and for Supplemental Figure 1a,b dots represent individual ventricles from six different HPS0076-hCOs generated in two independent preparations grown in different times. For No1 hCOs results in Figure 1c,f dots represent individual ventricles from three different No.1-hCOs generated in one preparation. For Figure 3c,e&f and for Supplemental Figure 7c dots represent individual ventricles from at least five different HPS0076-hCOs generated in three independent preparations grown at different times. For Supplemental Figure 5d dots represent individual ventricles from three different hCOs generated in two independent preparations per iPSC line grown at different times. For Supplemental Figure 8 dots represent individual ventricles from 4 different hCOs per condition and per genotype generated in one preparation for each genotype.

Dots in plots of Figures: 2e,f, 4e,g, 6b and of Supplemental Figures: 2c,e, 3b, 5b, 12a,e,f represent replicates each containing RNA/DNA/protein/cells extracted from a pool of two to three organoids each, i.e, 3 pools each containing 2 to 3 organoids so 6 to 9 organoids in total. More specifically, dots and western lanes in Figures 2e,f and 4b represent protein and RNA expression values from an isolate of a pool of three hCOs each. The hCOs were generated in two independent preparations grown in different times per iPSC line. For Figure 4e,g dots represent FCa results from an isolate of a pool of two hCOs each. The hCOs were generated in two independent preparations per iPSC line grown at different times. For Figure 6b and for Supplemental Figure 12a results of two or three replicates for dex respectively and 3 for dmso are depicted and they represent methylation levels from an isolate of a pool of two hCOs each. For Supplemental Figures 2c,e and 3b dots represent FCa results from an isolate of a pool of two hCOs each. The hCOs were generated in one preparation for panel 2e and in two preparations for panel 3b and 2c. For Supplemental Figure 5b dots represent RNA expression values from an isolate of a pool of two hCOs each. The hCOs were generated in two independent preparations grown in different times per iPSC line. For Supplemental Figure 12e,f dots represent RNA expression values or fold changes from an isolate of a pool of two hCOs each. The hCOs were generated in one preparation per iPSC line.

Dots in plots of Figure 5 and Figure S10 and S11 represent individual embryos. Each dot represents average counts from at least two cortical sections from one embryo. Embryos collected from two independent mothers were analysed for each staining.

For Figure S11 dots represent luciferase activity values of cells from independent cell culture wells for each condition and promoter.

Dots in Figure S12a,b represent RNA expression values or fold changes from isolates of independent cell culture wells for each genotype and treatment.

Box and whisker plots represent 25th to 75th percentile of the data with the center line representing the median and whiskers representing minima and maxima. Bar plots with error bars showing standard error of the mean (SEM). Plots and statistics for all Figures were generated using the GraphPad Prism 9 software. Significance: ****: p <=0.0001, ***: p <=0.001, **: p <=0.01, *: p <=0.05, ns: p >0.05.

## References

1. Barker, D.J.P. (2004). The developmental origins of chronic adult disease. Acta Paediatr Suppl, 26–33. 10.1080/08035320410022730.

2. Monk, C., Lugo-Candelas, C., and Trumpff, C. (2019). Prenatal Developmental Origins of Future Psychopathology: Mechanisms and Pathways. Annu Rev Clin Psychol 15. 10.1146/annurev-clinpsy-050718-095539.

3. Krontira, A.C., Cruceanu, C., and Binder, E.B. (2020). Glucocorticoids as Mediators of Adverse Outcomes of Prenatal Stress. Trends Neurosci 43, 394–405. 10.1016/j.tins.2020.03.008.

4. Carson, R., Monaghan-Nichols, A.P., DeFranco, D.B., and Rudine, A.C. (2016). Effects of antenatal glucocorticoids on the developing brain. Steroids 114, 25–32. 10.1016/j.steroids.2016.05.012.

5. Edwards, P.D., and Boonstra, R. (2018). Glucocorticoids and CBG during pregnancy in mammals: diversity, pattern, and function. Gen Comp Endocrinol 259, 122–130. 10.1016/j.ygcen.2017.11.012.

6. Harris, A., and Seckl, J. (2011). Glucocorticoids, prenatal stress and the programming of disease. Horm Behav 59, 279–289. 10.1016/j.yhbeh.2010.06.007.

7. Lajic, S., Karlsson, L., and Nordenström, A. (2018). Prenatal Treatment of Congenital Adrenal Hyperplasia: Long-Term Effects of Excess Glucocorticoid Exposure. Hormone Research in Pediatrics. 10.1159/000485100.

8. Committee on Obstetric Practice (2017). Antenatal Corticosteroid Therapy for Fetal Maturation. Obstetrics and Gynecology 130, 102–109.

9. Cao, G., Liu, J., and Liu, M. (2022). Global, Regional, and National Incidence and Mortality of Neonatal Preterm Birth, 1990-2019. JAMA Pediatr, 1–10. 10.1001/jamapediatrics.2022.1622.

10. Chawanpaiboon, S., Vogel, J.P., Moller, A.B., Lumbiganon, P., Petzold, M., Hogan, D., Landoulsi, S., Jampathong, N., Kongwattanakul, K., Laopaiboon, M., et al. (2019). Global, regional, and national estimates of levels of preterm birth in 2014: a systematic review and modelling analysis. Lancet Glob Health 7, e37– e46. 10.1016/S2214-109X(18)30451-0.

11. Vidavalur, R., Hussain, Z., and Naveed, H. (2022). Association of Survival at 22 Weeks’ Gestation With Use of Antenatal Corticosteroids and Mode of Delivery in the United States. JAMA Pediatr, 10–12.

12. The RECOVERY Collaborative Group (2020). Dexamethasone in Hospitalized Patients with Covid-19 — Preliminary Report. N Engl J Med, 1–11. 10.1056/NEJMoa2021436.

13. Mcintosh, J.J. (2020). Corticosteroid Guidance for Pregnancy during COVID-19 Pandemic. Americal Journal of Perinatology, 809–812.

14. Ninan, K., Liyanage, S.K., Murphy, K.E., Asztalos, E. V, and McDonald, S.D. (2022). Evaluation of Long-term Outcomes Associated With Preterm Exposure to Antenatal Corticosteroids: A Systematic Review and Meta-analysis. JAMA Pediatr, e220483. 10.1001/jamapediatrics.2022.0483.

15. Mcewen, B.S., Bowles, N.P., Gray, J.D., Hill, M.N., Hunter, R.G., Karatsoreos, I.N., and Nasca, C. (2015). Mechanisms of stress in the brain. Nat Neurosci 18. 10.1038/nn.4086.

16. Malik, S., Vinukonda, G., Vose, L.R., Diamond, D., Bhimavarapu, B.B.R., Hu, F., Zia, M.T., Hevner, R., Zecevic, N., and Ballabh, P. (2013). Neurogenesis continues in the third trimester of pregnancy and is suppressed by premature birth. Journal of Neuroscience 33, 411–423. 10.1523/JNEUROSCI.4445-12.2013.

17. Lancaster, M.A., and Knoblich, J.A. (2014). Generation of cerebral organoids from human pluripotent stem cells. Nat Protoc 9, 2329–2340. 10.1038/nprot.2014.158.

18. Cruceanu, C., Dony, L., Krontira, A.C., Fischer, D.S., Roeh, S., Di Giaimo, R., Kyrousi, C., Kaspar, L., Arloth, J., Czamara, D., et al. (2022). Cell-Type-Specific Impact of Glucocorticoid Receptor Activation on the Developing Brain: A Cerebral Organoid Study. American Journal of Psychiatry 179, 375–387. 10.1176/appi.ajp.2021.21010095.

19. Dehay, C., Kennedy, H., and Kosik, K.S. (2015). The Outer Subventricular Zone and Primate-Specific Cortical Complexification. Neuron 85, 683–694. 10.1016/j.neuron.2014.12.060.

20. Betizeau, M., Cortay, V., Patti, D., Pfister, S., Gautier, E., Bellemin-Ménard, A., Afanassieff, M., Huissoud, C., Douglas, R.J., Kennedy, H., et al. (2013). Precursor Diversity and Complexity of Lineage Relationships in the Outer Subventricular Zone of the Primate. Neuron 80, 442–457. 10.1016/j.neuron.2013.09.032.

21. Romero, C.D.J., Bruder, C., Tomasello, U., Sanz-anquela, J.M., and Borrell, V. (2015). Discrete domains of gene expression in germinal layers distinguish the development of gyrencephaly. EMBO J 34, 1859– 1874.

22. Florio, M., and Huttner, W.B. (2014). Neural progenitors, neurogenesis and the evolution of the neocortex. Development (Cambridge) 141, 2182–2194. 10.1242/dev.090571.

23. Matsumoto, N., Tanaka, S., Horiike, T., and Shinmyo, Y. (2020). A discrete subtype of neural progenitor crucial for cortical folding in the gyrencephalic mammalian brain. Elife, 1–26.

24. Borrell, V., and Götz, M. (2014). Role of radial glial cells in cerebral cortex folding. Curr Opin Neurobiol 27, 39–46. 10.1016/j.conb.2014.02.007.

25. Silbereis, J.C., Pochareddy, S., Zhu, Y., Li, M., and Sestan, N. (2016). The Cellular and Molecular Landscapes of the Developing Human Central Nervous System. Neuron 89, 248. 10.1016/j.neuron.2015.12.008.

26. Cruceanu, C., Ph, D., Dony, L., Sc, M., Krontira, A.C., Sc, M., Fischer, D.S., Sc, M., Roeh, S., and Sc, B. (2021). Cell-Type-Specific Impact of Glucocorticoid Receptor Activation on the Developing Brain: A Cerebral Organoid Study. American Journal of Psychiatry, 1–13. 10.1176/appi.ajp.2021.21010095.

27. Liu, T.M., Lee, E.H., Lim, B., and Shyh-Chang, N. (2015). Balancing Stem Cell Self-renewal and Differentiation with PLZF. Stem Cells, 277–287. 10.1002/stem.2270.

28. Avantaggiato, V., Pandolfi, P.P., Ruthardt, M., Hawe, N., Acampora, D., Pelicci, P.G., and Simeone, A. (1995). Developmental analysis of murine Promyelocyte Leukemia Zinc Finger (PLZF) gene expression: implications for the neuromeric model of the forebrain organization. The Journal of Neuroscience 15, 4927– 4942.

29. Pebworth, M.P., Ross, J., Andrews, M., Bhaduri, A., and Kriegstein, A.R. (2021). Human intermediate progenitor diversity during cortical development. Proc Natl Acad Sci U S A 118, 1–10. 10.1073/pnas.2019415118.

30. Hai, L., Szwarc, M.M., Lanza, D.G., Heaney, J.D., and Lydon, J.P. (2019). Using CRISPR / Cas9 engineering to generate a mouse with a conditional knockout allele for the promyelocytic leukemia zinc finger transcription factor. 1–8. 10.1002/dvg.23281.

31. Stepien, B.K., Vaid, S., and Huttner, W.B. (2021). Length of the Neurogenic Period—A Key Determinant for the Generation of Upper-Layer Neurons During Neocortex Development and Evolution. Front Cell Dev Biol 9, 1–20. 10.3389/fcell.2021.676911.

32. Magrinelli, E., Baumann, N., Wagener, R.J., Glangetas, C., Bellone, C., Jabaudon, D., and Klingler, E. (2022). Heterogeneous fates of simultaneously-born neurons in the cortical ventricular zone. Sci Rep 12, 1–11. 10.1038/s41598-022-09740-6.

33. Manuel, M., Georgala, P.A., Carr, C.B., Chanas, S., Kleinjan, D.A., Martynoga, B., Mason, J.O., Molinek, M., Pinson, J., Pratt, T., et al. (2007). Controlled overexpression of Pax6 in vivo negatively auto-regulates the Pax6 locus, causing cell-autonomous defects of late cortical progenitor proliferation with little effect on cortical arealization. Development 555, 545–555. 10.1242/dev.02764.

34. Sansom, S.N., Griffiths, D.S., Faedo, A., Kleinjan, D., Ruan, Y., Heyningen, V. Van, Rubenstein, J.L., and Livesey, F.J. (2009). The Level of the Transcription Factor Pax6 Is Essential for Controlling the Balance between Neural Stem Cell Self-Renewal and Neurogenesis. PLoS Genet 5, 20–23. 10.1371/journal.pgen.1000511.

35. Elsen, G.E., Bedogni, F., Hodge, R.D., and Bammler, T.K. (2018). The Epigenetic Factor Landscape of Developing Neocortex Is Regulated by Transcription Factors. Front Neurosci 12. 10.3389/fnins.2018.00571.

36. Anderson, T.R., Hedlund, E., and Carpenter, E.M. (2002). Differential Pax6 promoter activity and transcript expression during forebrain development. Mech Dev 114, 171–175.

37. Tyas, D.A., Simpson, T.I., Carr, C.B., Kleinjan, D.A., Van Heyningen, V., Mason, J.O., and Price, D.J. (2006). Functional conservation of Pax6 regulatory elements in humans and mice demonstrated with a novel transgenic reporter mouse. BMC Dev Biol 6, 1–11. 10.1186/1471-213X-6-21.

38. Provençal, N., Arloth, J., Cattaneo, A., Anacker, C., Cattane, N., Wiechmann, T., Röh, S., Ködel, M., Klengel, T., Czamara, D., et al. (2019). Glucocorticoid exposure during hippocampal neurogenesis primes future stress response by inducing changes in DNA methylation. Proceedings of the National Academy of Sciences 117, 201820842. 10.1073/pnas.1820842116.

39. Wiechmann, T., Röh, S., Sauer, S., Czamara, D., Arloth, J., Ködel, M., Beintner, M., Knop, L., Menke, A., Binder, E.B., et al. (2019). Identification of dynamic glucocorticoid-induced methylation changes at the FKBP5 locus. Clin Epigenetics, 1–14.

40. Roeh, S., Wiechmann, T., Sauer, S., Ködel, M., Binder, E.B., and Provençal, N. (2018). HAM-TBS: High-accuracy methylation measurements via targeted bisulfite sequencing. Epigenetics Chromatin 11, 1–10. 10.1186/s13072-018-0209-x.

41. Bothe, M., Buschow, R., and Meijsing, S.H. (2021). Glucocorticoid signaling induces transcriptional memory and universally reversible chromatin changes. Life Sci Alliance 4, 1–17. 10.26508/lsa.202101080.

42. Klengel, T., and Binder, E.B. (2015). Epigenetics of Stress-Related Psychiatric Disorders and Gene × Environment Interactions. Neuron 86, 1343–1357. 10.1016/j.neuron.2015.05.036.

43. Lee, J.J., Wedow, R., Okbay, A., Kong, E., Maghzian, O., Zacher, M., Nguyen-Viet, T.A., Bowers, P., Sidorenko, J., Linnér, R.K., et al. (2018). Gene discovery and polygenic prediction from a genome-wide association study of educational attainment in 1.1 million individuals. Nat Genet 50. 10.1038/s41588-018-0147-3.

44. Okbay, A., Wu, Y., Wang, N., Jayashankar, H., Bennett, M., Nehzati, S.M., Sidorenko, J., Kweon, H., Goldman, G., Gjorgjieva, T., et al. (2022). Polygenic prediction of educational attainment within and between families from genome-wide association analyses in 3 million individuals. Nat Genet 54. 10.1038/s41588-022-01016-z.

45. Makowski, C., Loughnan, R., Jernigan, T.L., Seibert, T.M., Hagler, D.J., Smeland, O.B., Motazedi, E., Chu, Y., Lin, A., Hindley, G., et al. (2022). Vertex-wise multivariate genome-wide association study identifies 780 unique genetic loci associated with cortical morphology. Neuroimage, 1–19. 10.1016/j.neuroimage.2021.118603.Vertex-wise.

46. Schiller, B.J., Chodankar, R., Watson, L.C., Stallcup, M.R., and Yamamoto, K.R. (2014). Glucocorticoid receptor binds half sites as a monomer and regulates specific target genes. Genome Biol 15, 1–16. 10.1186/s13059-014-0418-y.

47. Smith, D.J., Nicholl, B.I., Cullen, B., Martin, D., Ul-haq, Z., Evans, J., Gill, J.M.R., Roberts, B., Gallacher, J., Mackay, D., et al. (2013). Prevalence and Characteristics of Probable Major Depression and Bipolar Disorder within UK Biobank: Cross-Sectional Study of 172, 751 Participants. 8, 1–7. 10.1371/journal.pone.0075362.

48. Bartrés-Faz, D., González-Escamilla, G., Vaqué-Alcázar, L., Abellaneda-Pérez, K., Valls-Pedret, C., Ros, E., and Grothe, M.J. (2019). Characterizing the molecular architecture of cortical regions associated with high educational attainment in older individuals. Journal of Neuroscience 39, 4566–4575. 10.1523/JNEUROSCI.2370-18.2019.

49. Ge, T., Chen, C.Y., Doyle, A.E., Vettermann, R., Tuominen, L.J., Holt, D.J., Sabuncu, M.R., and Smoller, J.W. (2019). The Shared Genetic Basis of Educational Attainment and Cerebral Cortical Morphology. Cerebral Cortex 29, 3471–3481. 10.1093/cercor/bhy216.

50. Kim, J.P., Seo, S.W., Shin, H.Y., Ye, B.S., Yang, J.J., Kim, C., Kang, M., Jeon, S., Kim, H.J., Cho, H., et al. (2015). Effects of education on aging-related cortical thinning among cognitively normal individuals. Neurology 85, 806–812. 10.1212/WNL.0000000000001884.

51. Vaqué-alcázar, L., Sala-llonch, R., and Valls-pedret, C. (2017). Differential age-related gray and white matter impact mediates educational influence on elders ’ cognition. Brain Imaging Behav 11, 318–332. 10.1007/s11682-016-9584-8.

52. Kvist, T., Sammallahti, S., Lahti-Pulkkinen, M., Cruceanu, C., Czamara, D., Dieckmann, L., Tontsch, A., Röh, S., Rex-Haffner, M., Wolford, E., et al. (2022). Cohort profile: InTraUterine sampling in early pregnancy (ITU), a prospective pregnancy cohort study in Finland: Study design and baseline characteristics. BMJ Open 12, 1–11. 10.1136/bmjopen-2021-049231.

53. N, B. (2006). Bayley scales of infant and toddler development (Pearson: PsychCorp) https://doi.org/10.1037/t14978-000.

54. Garcia, M.T., Chang, Y., Arai, Y., and Huttner, W.B. (2016). S-Phase Duration Is the Main Target of Cell Cycle Regulation in Neural Progenitors of Developing Ferret Neocortex. J Comp Neurol 470, 456–470. 10.1002/cne.23801.

55. McEwen, B.S., Nasca, C., and Gray, J.D. (2016). Stress Effects on Neuronal Structure: Hippocampus, Amygdala, and Prefrontal Cortex. Neuropsychopharmacology 41, 3–23. 10.1038/npp.2015.171.

56. Popoli, M., Yan, Z., McEwen, B.S., and Sanacora, G. (2012). The stressed synapse: The impact of stress and glucocorticoids on glutamate transmission. Nat Rev Neurosci 13, 22–37. 10.1038/nrn3138.

57. Melamed, N., Asztalos, E., Murphy, K., Zaltz, A., Redelmeier, D., Shah, B.R., and Barrett, J. (2019). Neurodevelopmental disorders among term infants exposed to antenatal corticosteroids during pregnancy: a population-based study. 3–10. 10.1136/bmjopen-2019-031197.

58. Räikkönen, K., Gissler, M., and Kajantie, E. (2020). Associations Between Maternal Antenatal Corticosteroid Treatment and Mental and Behavioral Disorders in Children Supplemental content. JAMA 323, 1924–1933. 10.1001/jama.2020.3937.

59. Tsiarli, M.A., Rudine, A., Kendall, N., Pratt, M.O., Krall, R., Thiels, E., Defranco, D.B., and Monaghan, A.P. (2017). Antenatal dexamethasone exposure differentially affects distinct cortical neural progenitor cells and triggers long-term changes in murine cerebral architecture and behavior. Transl Psychiatry. 10.1038/tp.2017.65.

60. Mcewen, B.S. (2005). Glucocorticoids, depression, and mood disorders: structural remodeling in the brain. Metabolism 54, 20–23. 10.1016/j.metabol.2005.01.008.

61. Koyanagi-Aoi, M., Ohnuki, M., Takahashi, K., Okita, K., Noma, H., Sawamura, Y., Teramoto, I., Narita, M., Sato, Y., Ichisaka, T., et al. (2013). Differentiation-defective phenotypes revealed by large-scale analyses of human pluripotent stem cells. Proc Natl Acad Sci U S A 110, 20569–20574. 10.1073/pnas.1319061110.

62. Okita, K., Matsumura, Y., Sato, Y., Okada, A., Morizane, A., Okamoto, S., Hong, H., Nakagawa, M., Tanabe, K., Tezuka, K.I., et al. (2011). A more efficient method to generate integration-free human iPS cells. Nat Methods 8, 409–412. 10.1038/nmeth.1591.

63. Cárdenas, A., Villalba, A., de Juan Romero, C., Picó, E., Kyrousi, C., Tzika, A.C., Tessier-Lavigne, M., Ma, L., Drukker, M., Cappello, S., et al. (2018). Evolution of Cortical Neurogenesis in Amniotes Controlled by Robo Signaling Levels. Cell 174, 590–606.e21. 10.1016/j.cell.2018.06.007.

64. Andrews Simon, Krueger Felix, Segonds-Pichon Anne, Biggins Laura, Krueger Christel, M.J. (2019). fastQC. https://www.bioinformatics.babraham.ac.uk/projects/fastqc/.

65. Martin, M. (2011). Cutadapt removes adapter sequences from high-throughput sequencing reads. EMBnet J 17, 10–12.

66. Patro, R., Duggal, G., Love, M.I., Irizarry, R.A., and Kingsford, C. (2017). Salmon provides fast and bias-aware quantification of transcript expression. Nat Methods 14, 417–419. 10.1038/nmeth.4197.

67. Love, M.I., Huber, W., and Anders, S. (2014). Moderated estimation of fold change and dispersion for RNA-seq data with DESeq2. Genome Biol 15, 1–21. 10.1186/s13059-014-0550-8.

68. Saito, T. (2006). In vivo electroporation in the embryonic mouse central nervous system. Nat Protoc 1, 1552–1558. 10.1038/nprot.2006.276.

69. Buchsbaum, I.Y., Kielkowski, P., Giorgio, G., Neill, A.C.O., Giaimo, R. Di Kyrousi, C., Khattak, S., Sieber, S.A., Robertson, S.P., and Cappello, S. (2020). ECE2 regulates neurogenesis and neuronal migration during human cortical development. EMBO Rep 49, 1–24. 10.15252/embr.201948204.

70. Klaus, J., Kanton, S., Kyrousi, C., Ayo-Martin, A.C., Di Giaimo, R., Riesenberg, S., O’Neill, A.C., Camp, J.G., Tocco, C., Santel, M., et al. (2019). Altered neuronal migratory trajectories in human cerebral organoids derived from individuals with neuronal heterotopia. Nat Med 25, 561–568. 10.1038/s41591-019-0371-0.

71. Jobe, A.H., Kemp, M., Schmidt, A., Takahashi, T., Newnham, J., and Milad, M. (2021). Antenatal corticosteroids: a reappraisal of the drug formulation and dose. Pediatr Res. 10.1038/s41390-020-01249-w.

72. Kelava, I., Chiaradia, I., Pellegrini, L., Kalinka, A.T., and Lancaster, M.A. (2021). Androgens increase excitatory neurogenic potential in human brain organoids. 10.1038/s41586-021-04330-4.

73. McManus, J.M., and Sharifi, N. (2020). Structure-dependent retention of steroid hormones by common laboratory materials. Journal of Steroid Biochemistry and Molecular Biology 198, 1–19. 10.1016/j.jsbmb.2019.105572.

74. Spoelhof, B., Ray, S.D., and Wayne, F. (2014). Fludrocortisone Cortisol/Hydrocortisone. In Encyclopedia of Toxicology, pp. 1038–1042. 10.1016/B978-0-12-386454-3.00293-1.

75. Kashiwagi, Y., Kato, N., Sassa, T., Nishitsuka, K., Yamamoto, T., Takamura, H., and Yamashita, H. (2010). Cotylenin a inhibits cell proliferation and induces apoptosis and PAX6 mRNA transcripts in retinoblastoma cell lines. Mol Vis 16, 970–982.

76. Martin, M. Cutadapt removes adapter sequences from high-throughput sequencing reads.

77. Krueger, F., and Andrews, S.R. (2011). Bismark: A flexible aligner and methylation caller for Bisulfite-Seq applications. Bioinformatics 27, 1571–1572. 10.1093/bioinformatics/btr167.

78. Akalin, A., Kormaksson, M., Li, S., Garrett-Bakelman, F.E., Figueroa, M.E., Melnick, A., and Mason, C.E. (2012). MethylKit: a comprehensive R package for the analysis of genome-wide DNA methylation profiles. Genome Biol 13. 10.1186/gb-2012-13-10-R87.

79. Benjamini, Y., Krieger, A.M., and Yekutieli, D. (2006). Adaptive linear step-up procedures that control the false discovery rate. Biometrika 93, 491–507. 10.1093/biomet/93.3.491.

80. Wu, T., Hu, E., Xu, S., Chen, M., Guo, P., Dai, Z., Feng, T., Zhou, L., Tang, W., Zhan, L., et al. (2021). clusterProfiler 4.0: A universal enrichment tool for interpreting omics data. The Innovation 2. 10.1016/j.xinn.2021.100141.

81. Yu, G., Wang, L.-G., Han, Y., and He, Q.-Y. (2012). clusterProfiler: an R Package for Comparing Biological Themes Among Gene Clusters. OMICS 16, 284–287. 10.1089/omi.2011.0118.

82. G, Y. (2023). Enrichplot: Visualization of Functional Enrichment Result. R package version 1.20.0.

83. Becht, E., McInnes, L., Healy, J., Dutertre, C.A., Kwok, I.W.H., Ng, L.G., Ginhoux, F., and Newell, E.W. (2019). Dimensionality reduction for visualizing single-cell data using UMAP. Nat Biotechnol 37, 38–47. 10.1038/nbt.4314.

84. Wolf, F.A., Angerer, P., and Theis, F.J. (2018). Open Access SCANPY: large-scale single-cell gene expression data analysis. Genome Biol, 1–5.

85. Arnold, C.D., Gerlach, D., Stelzer, C., Boryń, Ł.M., Rath, M., and Stark, A. (2013). Genome-wide quantitative enhancer activity maps identified by STARR-seq. Science (1979) 339, 1074–1077. 10.1126/science.1232542.

86. Schöne, S., Bothe, M., Einfeldt, E., Borschiwer, M., Benner, P., Vingron, M., Thomas-Chollier, M., and Meijsing, S.H. (2018). Synthetic STARR-seq reveals how DNA shape and sequence modulate transcriptional output and noise. PLoS Genet 14, 1–24. 10.1371/journal.pgen.1007793.

87. Rogatsky, I., Trowbridge, J.M., and Garabedian, M.J. (1997). Glucocorticoid receptor-mediated cell cycle arrest is achieved through distinct cell-specific transcriptional regulatory mechanisms. Mol Cell Biol 17, 3181–3193. 10.1128/mcb.17.6.3181.

88. Team, R.C. (2017). R: A language and environment for statistical computing. https://www.r-project.org/.

89. Elsworth, B., Lyon, M., Alexander, T., Liu, Y., Matthews, P., Hallett, J., Bates, P., Palmer, T., Haberland, V., Davey, G., et al. (2020). The MRC IEU OpenGWAS data infrastructure. bioRxiv, 2020.08.10.244293.

90. Bycroft, C., Freeman, C., Petkova, D., Band, G., Elliott, L.T., Sharp, K., Motyer, A., Vukcevic, D., Delaneau, O., O’Connell, J., et al. (2018). The UK Biobank resource with deep phenotyping and genomic data. Nature 562, 203–209. 10.1038/s41586-018-0579-z.

91. Buniello, A., Macarthur, J.A.L., Cerezo, M., Harris, L.W., Hayhurst, J., Malangone, C., McMahon, A., Morales, J., Mountjoy, E., Sollis, E., et al. (2019). The NHGRI-EBI GWAS Catalog of published genome-wide association studies, targeted arrays and summary statistics 2019. Nucleic Acids Res 47, D1005– D1012. 10.1093/nar/gky1120.

92. Elliott, L.T., Sharp, K., Alfaro-Almagro, F., Shi, S., Miller, K.L., Douaud, G., Marchini, J., and Smith, S.M. (2018). Genome-wide association studies of brain imaging phenotypes in UK Biobank. Nature 562, 210–216. 10.1038/s41586-018-0571-7.

93. van der Loo, M.P.J. (2014). The stringdist package for approximate string matching. R Journal 6, 111–122. 10.32614/rj-2014-011.

94. Hemani, G., Zheng, J., Elsworth, B., Wade, K.H., Haberland, V., Baird, D., Laurin, C., Burgess, S., Bowden, J., Langdon, R., et al. (2018). The MR-base platform supports systematic causal inference across the human phenome. Elife 7, 1–29. 10.7554/eLife.34408.

95. Palmer, T.M., Sterne, J.A.C., Harbord, R.M., Lawlor, D.A., Sheehan, N.A., Meng, S., Granell, R., Smith, G.D., and Didelez, V. (2011). Instrumental variable estimation of causal risk ratios and causal odds ratios in mendelian randomization analyses. Am J Epidemiol 173, 1392–1403. 10.1093/aje/kwr026.

96. Benjamini, Y., and Hochberg, Y. (1995). Controlling the False Discovery Rate: A Practical and Powerful Approach to Multiple Testing. Journal of the Royal Statistical Society: Series B (Methodological) 57, 289–300. 10.1111/j.2517-6161.1995.tb02031.x.

97. Bland, j. M., and Altman, D.G. (1995). Multiple significance tests: The Bonferroni method. Bmj 310, 170. 10.1136/bmj.310.6973.170.

98. Faillenot, I., Heckemann, R.A., Frot, M., and Hammers, A. (2017). Macroanatomy and 3D probabilistic atlas of the human insula. Neuroimage 150, 88–98. 10.1016/j.neuroimage.2017.01.073.

99. Mori, S., Oishi, K., Jiang, H., Jiang, L., Li, X., Akhter, K., Hua, K., Faria, A. V., Mahmood, A., Woods, R., et al. (2008). Stereotaxic white matter atlas based on diffusion tensor imaging in an ICBM template. Neuroimage 40, 570–582. 10.1016/j.neuroimage.2007.12.035.

100. Hua, K., Zhang, J., Wakana, S., Jiang, H., Li, X., Reich, D.S., Calabresi, P.A., Pekar, J.J., van Zijl, P.C.M., and Mori, S. (2008). Tract Probability Maps in Stereotaxic Spaces: Analyses of White Matter Anatomy and Tract-Specific Quantification. Neuroimage 32, 736–740. 10.1016/j.neuroimage.2007.07.053.Tract.

101. Warrington, S., Bryant, K.L., Khrapitchev, A.A., Sallet, J., Charquero-Ballester, M., Douaud, G., Jbabdi, S., Mars, R.B., and Sotiropoulos, S.N. (2020). XTRACT – Standardised protocols for automated tractography in the human and macaque brain. Neuroimage 217, 1–15. 10.1016/j.neuroimage.2020.116923.

102. Masharani, U., Shiboski, S., Eisner, M.D., Katz, P.P., Janson, S.L., Granger, D.A., and Blanc, P.D. (2005). Impact of exogenous glucocorticoid use on salivary cortisol measurements among adults with asthma and rhinitis. Psychoneuroendocrinology 30, 744–752. 10.1016/j.psyneuen.2005.03.003.

103. Pruessner, J.C., Kirschbaum, C., Meinlschmid, G., and Hellhammer, D.H. (2003). Two formulas for computation of the area under the curve represent measures of total hormone concentration versus time-dependent change. Psychoneuroendocrinology 28, 916–931. 10.1016/S0306-4530(02)00108-7.

104. Wickham, H. (2016). ggplot2: Elegant Graphics for Data Analysis. Springer-Verlag New York.

105. Schindelin, J., Arganda-Carreras, I., Frise, E., Kaynig, V., Longair, M., Pietzsch, T., Preibisch, S., Rueden, C., Saalfeld, S., Schmid, B., et al. (2012). Fiji: an open-source platform for biological-image analysis. Nature Methods 2012 9:79, 676–682. 10.1038/nmeth.2019.

106. Trujillo-Ortiz, A., and Hernandez-Walls, R. (2003). D’Agostino-Pearson’s K2 test for assessing normality of data using skewness and kurtosis. 10.13140/RG.2.2.19734.98889.

