## Supplemental Results for "Human neurogenesis is altered via glucocorticoid-mediated regulation of *ZBTB16* expression"

Anthi C. Krontira *et al.*

### Supplemental Results

#### Morphology of the PAX6+EOMES+ basal progenitors

We studied the morphology of the PAX6+EOMES+ BPs. In macaques, these cells were shown to have both IP-related morphologies with no processes and bRG- related morphologies with a basal, an apical or both processes[1]. In addition, the transcriptional and morphological diversity of human EOMES-expressing cells was studied recently. The authors found, among others, one cluster of EOMES+ cells that co-express SOX2, a co-factor of PAX6, and termed them RG-like-nIPs. Interestingly, they show that although these cells are biased to have an RG-like phenotype with an apical, basal or both processes, they are also found in all other morphologies which include short multipolar, spherical and monopolar/bipolar cells with horizontal orientations[2]. To answer the question of cell morphology we performed GFP cell morphology reconstructions of PAX6+EOMES+ BPs in the electroporated hCOs. We indeed find all types of morphologies for these cells, cells with no processes, with an apical process, a basal process, bipolar cells and monopolar/bipolar horizontal cells, thus exhibiting all physiological morphologies found in these cells in species with a gyrified brain (Figure S6).

#### Self-renewing capacity of ZBTB16-overexpressing cells

A distinctive trait differentiating between IPs and bRGs is their capacity of self-renewal, with the latter cell type undergoing more self-renewing and proliferative divisions. Interestingly, it has been shown that the cell-cycle characteristics of the PAX6+EOMES+ cells lie in-between the ones of bRGs and IPs, suggesting that PAX6+EOMES+ cells may also have an intermediate phenotype in relation to division patterns being both self-renewing and neurogenic[3]. We show that there is a significant increase of Ki67+ cells in ZBTB16-electroporated organoids (Figure 3B & C), suggesting an increase of proliferation capacity. To further support this, we performed cell cycle re-entry analysis in mice, as cell cycle dynamics are well known for this species. Mice electroporated at E13.5 with a control or ZBTB16-overexpressing plasmid received a BRDU injection at E15.5, 24 hours before they were sacrificed at E16.5. We then stained for Ki67, BRDU and GFP for the plasmid and quantified BRDU+Ki67+ GFP cells to label cells that re-entered the cell cycle. Indeed, we find more BRDU+Ki67+ cells in bins A and B, the same bins where we find the increase in Pax6+Eomes+ cells, in GFP cells overexpressing ZBTB16 as compared to control plasmids (Figure S8). This indicates that ZBTB16 prompts the cells to re-enter the cell cycle.

#### Dex effects on the PAX6+EOMES+ progenitors' transcriptional landscape

To study whether dex treatment affected the transcriptomic profile of PAX6+EOMES+ BPs, we used a single-cell RNA sequencing (sc-RNA seq) dataset of HPS0076 hCOs treated with 100nM of dex for 10 days (treatment initiation at day 60). Indeed, even if their total numbers are low (Table S1), the PAX6+EOMES+ progenitor cells found after dex treatment are 3.38 times more compared to vehicle even when we account for the difference in total cell numbers between the treatment groups, a similar fold change to the ones we find with our immunostainings in Figure 1C (Figure S1C), thus further supporting the effects of GCs in increasing the numbers of these double positive progenitors. We sub-setted our dataset to the PAX6+EOMES+ cells and performed differential expression (DE) analysis for treatment effects on those. We found only two DE genes significant at an FDR (false discovery rate) threshold of 10%, *MEST* and *RSPO3* (Table S2) which are both WNT signaling regulators and thus very important for neurodevelopment[19,20]. In addition, there were 221 DE genes significant at the nominal p-value level of < 0.05. We performed Gene Set Enrichment analysis (GSEA) to identify gene sets overrepresented in the PAX6+EOMES+ cells after treatment. We uncovered many gene sets associated with cell fate commitment and glia cell proliferation and differentiation (Figure S1D & Table S3), highlighting the importance of these cells in regulating neurogenic processes and possibly pointing to the fact that they belong in the RG population, as has already been shown previously[21].

### Supplemental Figures

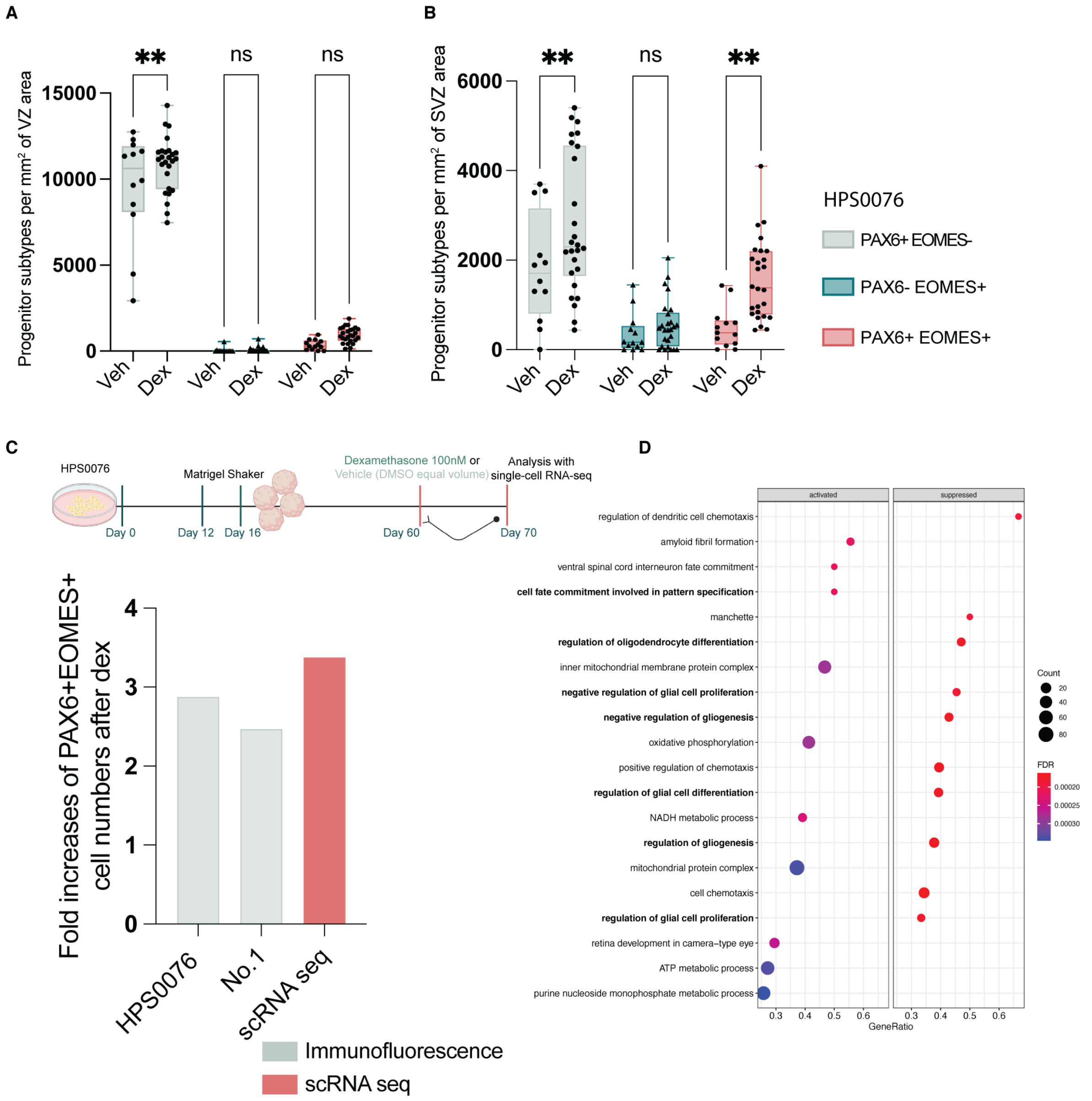


**Figure S1I Glucocorticoids increase PAX6+EOMES+ basal progenitors. A,** Quantification of the progenitor subtypes in each treatment condition normalized by $mm^2$ of quantified VZ area in hCOs produced from HPS0076 iPSCs. **B,** Quantification of the progenitor subtypes in each treatment condition normalized by $mm^2$ of quantified SVZ area in hCOs produced from HPS0076 iPSCs. **C,** Treatment paradigm of HPS0076 hCOs that were analysed with single-cell RNA sequencing. Fold increases of the PAX6+EOMES+ cell numbers after dex in the Progenitor clusters of the scRNA-seq dataset, and in the germinal zones of the Immunofluorescent quantifications shown in Figure 1B,C. **D,** Gene sets enriched at an FDR threshold of 5% in the PAX6+EOMES+ after dexamethasone as found with GSEA. hCOs, human cerebral organoids; VZ, ventricular zone; SVZ, subventricular zone; Veh, vehicle; Dex, dexamethasone; GSEA, gene set enrichment analysis; FDR, false discovery rate. For **A & B** significance was tested with two-way ANOVA with Benjamini, Krieger and Yekutieli multiple testing correction (A: p.treatment= 0.01, p.progenitor_type< 0.0001, p.interaction= 0.12, F.interaction= 2.09, F.treatment= 6.18, F.progenitor_type= 670.3, DF.interaction= 2, DF.treatment= 1, DF.progenitor_type= 2 / B: p.treatment= 0.0003, p.progenitor_type< 0.0001, p.interaction = 0.15, F.interaction = 1.91, F.treatment= 14.19, F.progenitor_type= 29.21, DF.interaction= 2, DF.treatment= 1, DF.progenitor_type= 2).

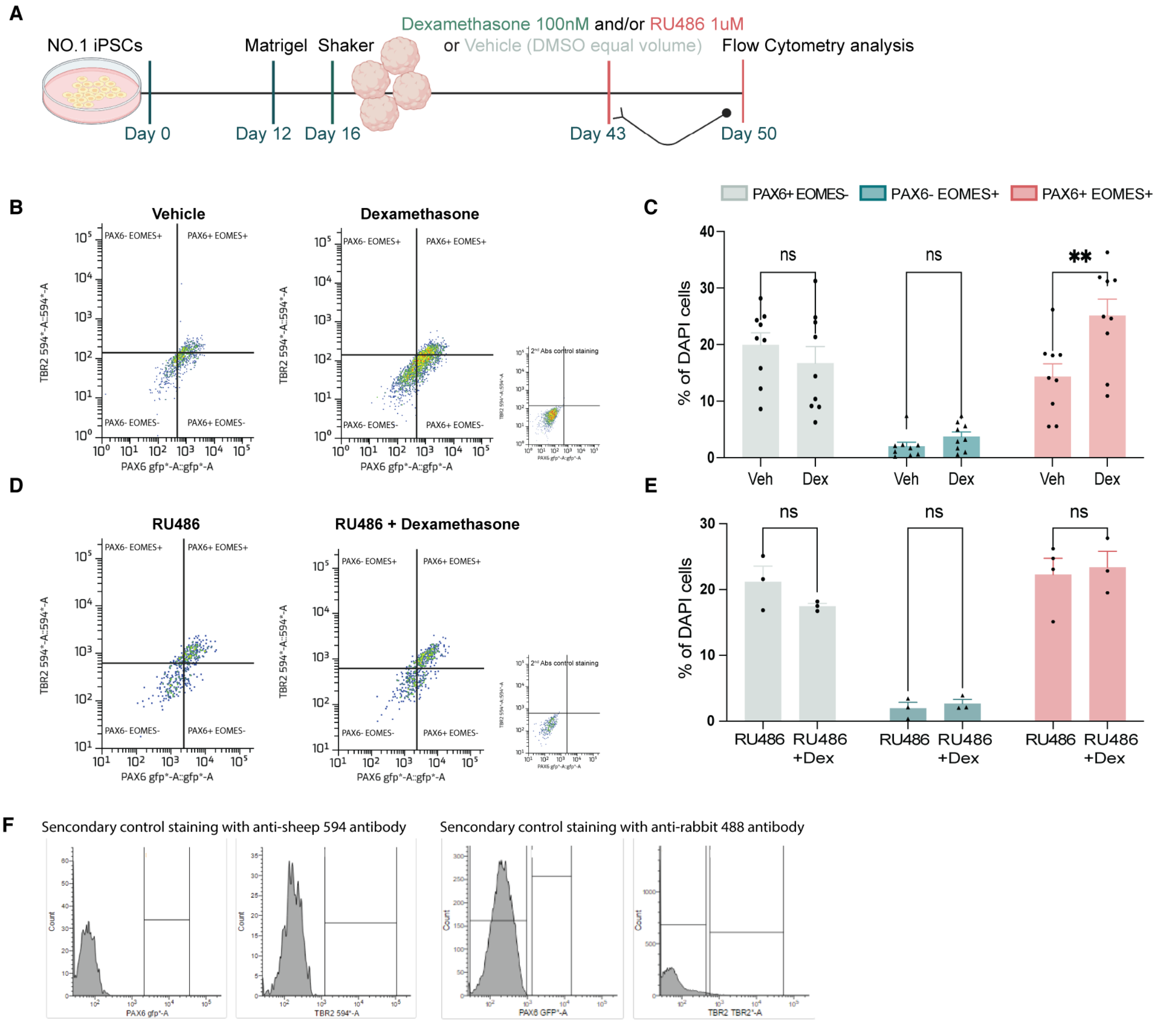


**Figure S2| The glucocorticoid receptor mediates the effects of glucocorticoids on basal progenitors. A,** Treatment and analysis workflow in hCOs derived from Line 2- iPSCs for co-treatment of dex and its antagonist RU-486. **B,** Representative images of FCa of hCOs per treatment condition. **C,** Quantification of the FCa results. Percentages of DAPI cells in each progenitor subtype and treatment condition. **D,** Representative images of FCa of hCOs per treatment condition. **E,** Quantification of the FCa results. Percentages of DAPI cells in each progenitor subtype and treatment condition. **F,** Representative images of secondary control stainings with anti-sheep 594 and anti-rabbit 488 antibodies. DMSO, dimethyl sulfoxide; RU486, mifepristone- GR antagonist; FCa, flow cytometry; hCOs, human cerebral organoids; Veh, vehicle; Dex, dexamethasone. Significance was tested with two-way ANOVA with Benjamini, Krieger and Yekutieli multiple testing correction (C: p= 0.0093, F= 3.62, DF= 4/ E: p= 0.77, F= 0.44, DF= 4). Post-hoc p-values: *** <=0.001, ns >0.05

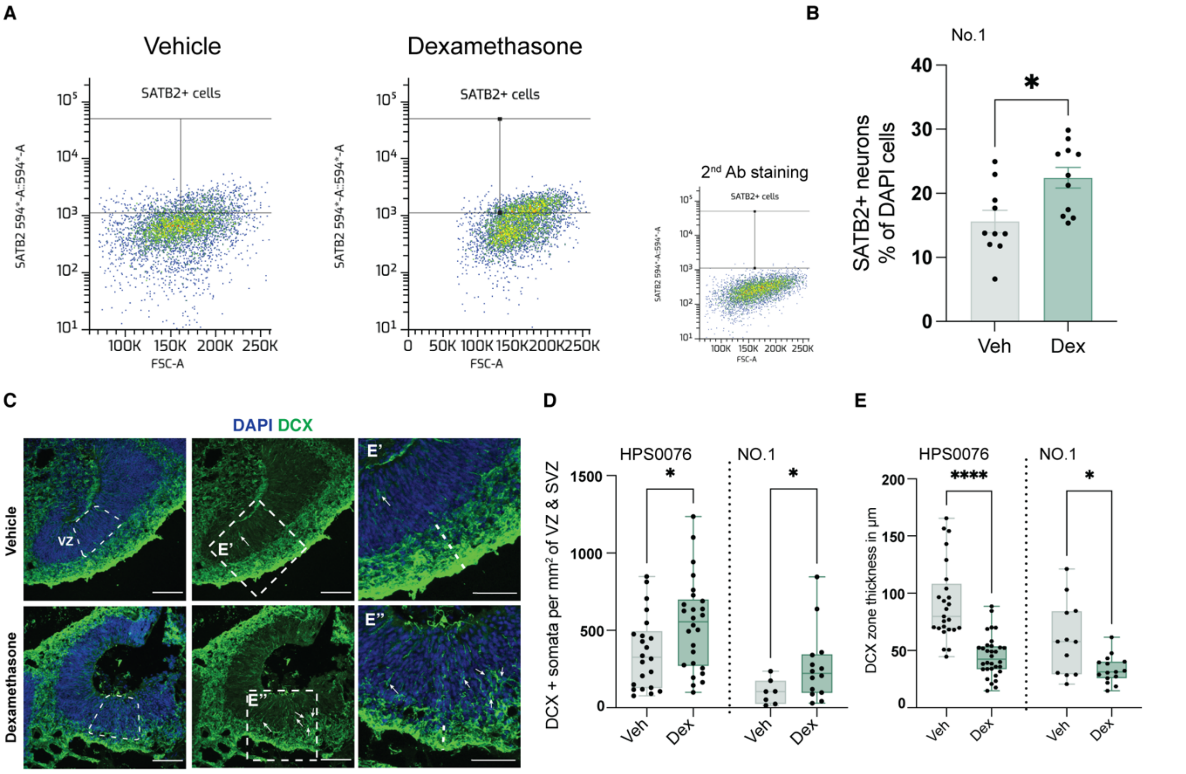


**Figure S3l Glucocorticoid effects on progenitors and neurons in hCOs. A,** Representative images of FCa of hCOs per treatment condition. **B,** Quantification of the FCa results. Percentages of DAPI cells that are positive for SATB2, a layer IV neuronal marker, in each treatment condition. Veh, vehicle; Dex, dexamethasone; FCa, flow cytometry analysis. **C,** Representative images of day 50 hCOs at vehicle and dex conditions stained for DCX and DAPI. Arrows indicate DCX positive somata in the VZ; Scale bars, 100µm. **D,** Quantification of DCX somata found in the VZ normalized per $mm^2$ of area. **E,** Quantification of DCX zone thickness in µm. For **D, E** significance was tested with Mann-Whitney comparison of dex versus dmso. p-values: **** <=0.0001, * <=0.05.

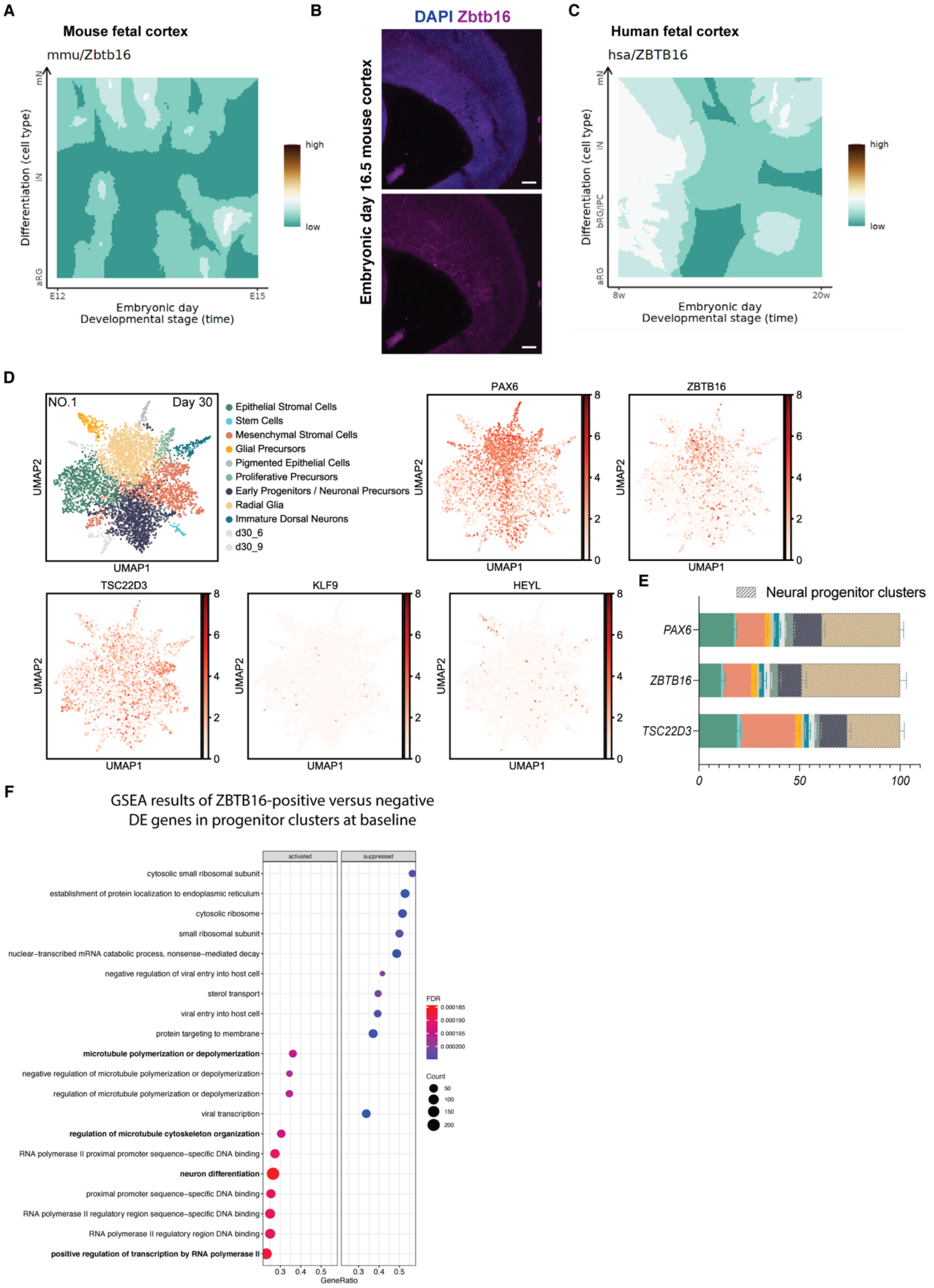


**Figure S4I *ZBTB16* expression profile in different species. A,** *ZBTB16* expression in the mouse developing cortex from Klinger et al., Science, 2021. **B,** Representative images of E16.5 mouse cortex stained for ZBTB16 and DAPI, note very low to absent staining. **C,** *ZBTB16* expression in the human developing cortex from Klinger et al., Science, 2021. Note the relative higher expression off *ZBTB16* in human than in mouse brain. **D,** UMAP plots of cell clusters and TFs in day 30 No.1 hCOs from Cruceanu et al., 2022. **E,** Percent of the fraction of total cells positive for each TF in each cluster. **F,** Gene sets enriched at an FDR threshold of 5% in the ZBTB16+ versus ZBTB16- progenitor cells as found with GSEA. Color scheme follows the one in Figure S4D. aRG, apical Radial Glia; bRG, basal Radial Glia; IPC, intermediate progenitor; iN, immature neuron; mN, mature neuron.

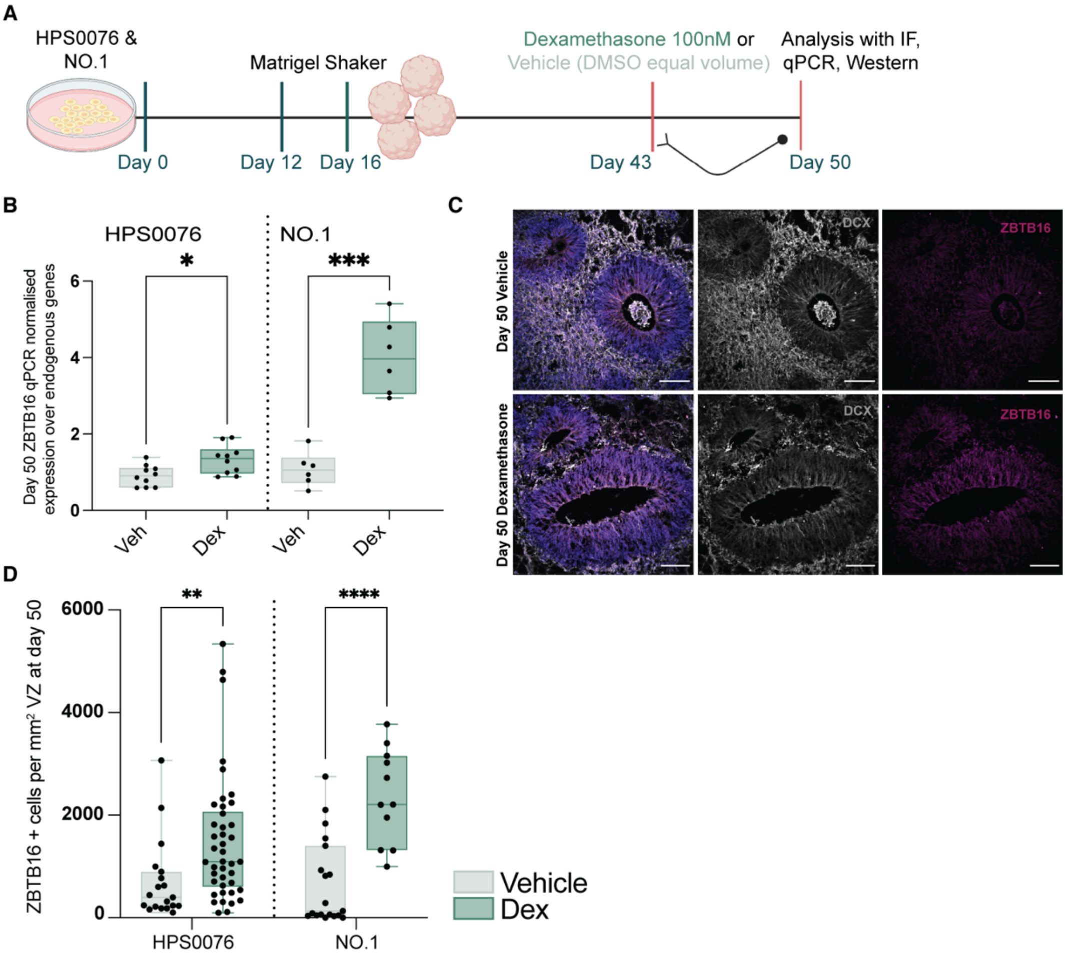


**Figure S5I Glucocorticoids increase *ZBTB16* expression. A,** Schematic of the dex-treatment paradigm and analysis workflow in hCOs from Line 1 and Line 2 iPSCs. **B,** Relative *ZBTB16* RNA expression in day 50 hCOs exposed to 7 days of 100nM dex or vehicle. **C**, Representative images of day 50 hCOs at vehicle and dex conditions stained for the immature neuronal marker DCX and ZBTB16. Scale bars, 100µm. **D,** Quantification of ZBTB16+ cells found in the VZ normalized per mm² of area. qPCR, quantitative polymerase chain reaction; IF, immunofluorescence; Veh, vehicle; Dex, dexamethasone; hCOs, human cerebral organoids. Significance was tested with two tailed Mann-Whitney comparison between treatment conditions. Mann-Whitney p-values: **** <=0.0001, *** <=0.001, ** <=0.01, * <=0.5

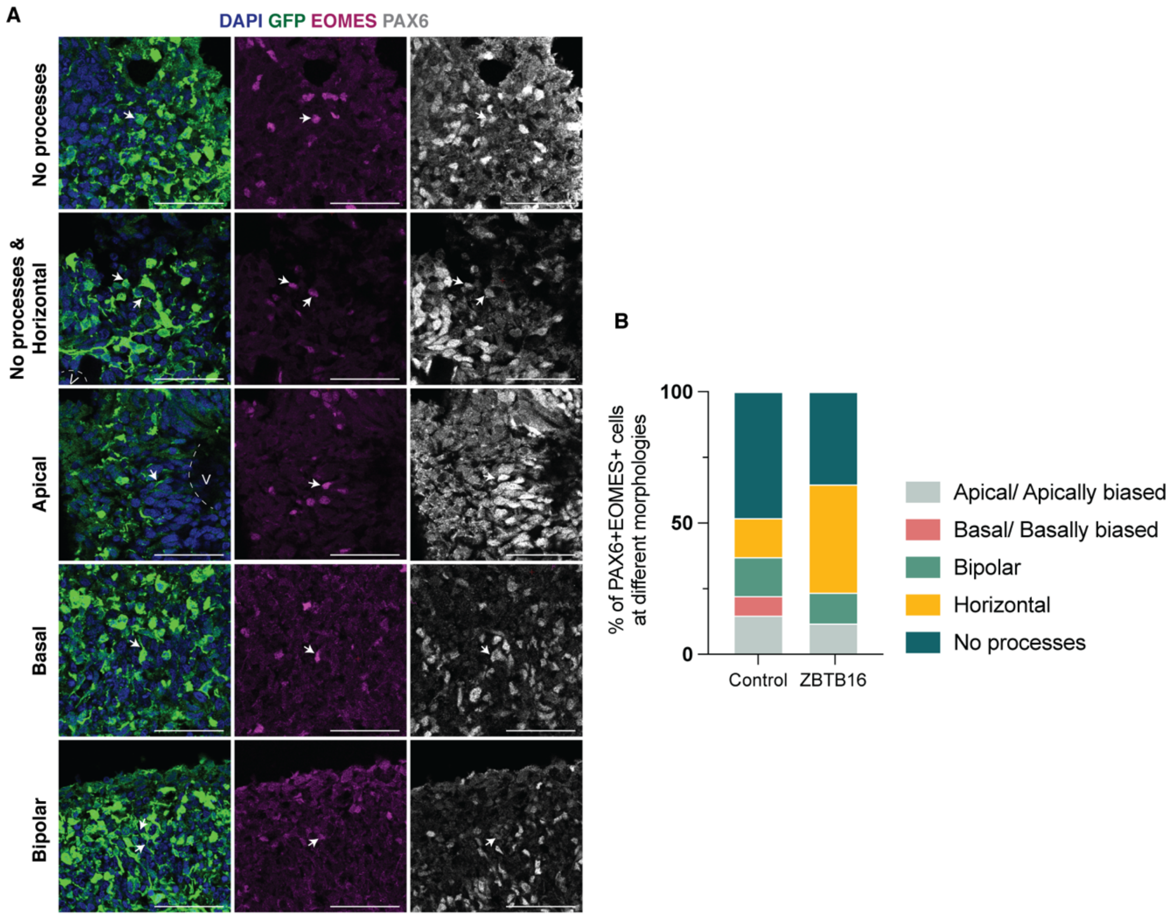


**Figure S6l PAX6+EOMES+ basal progenitors' morphology. A,** Representative images of morphology types of PAX6+EOMS+ progenitors. **B,** Percentage of PAX6+EOMES+ cells at the different morphology types when electroporated with a control or a ZBTB16-overexpressing plasmid. Scale bars, 50µm.

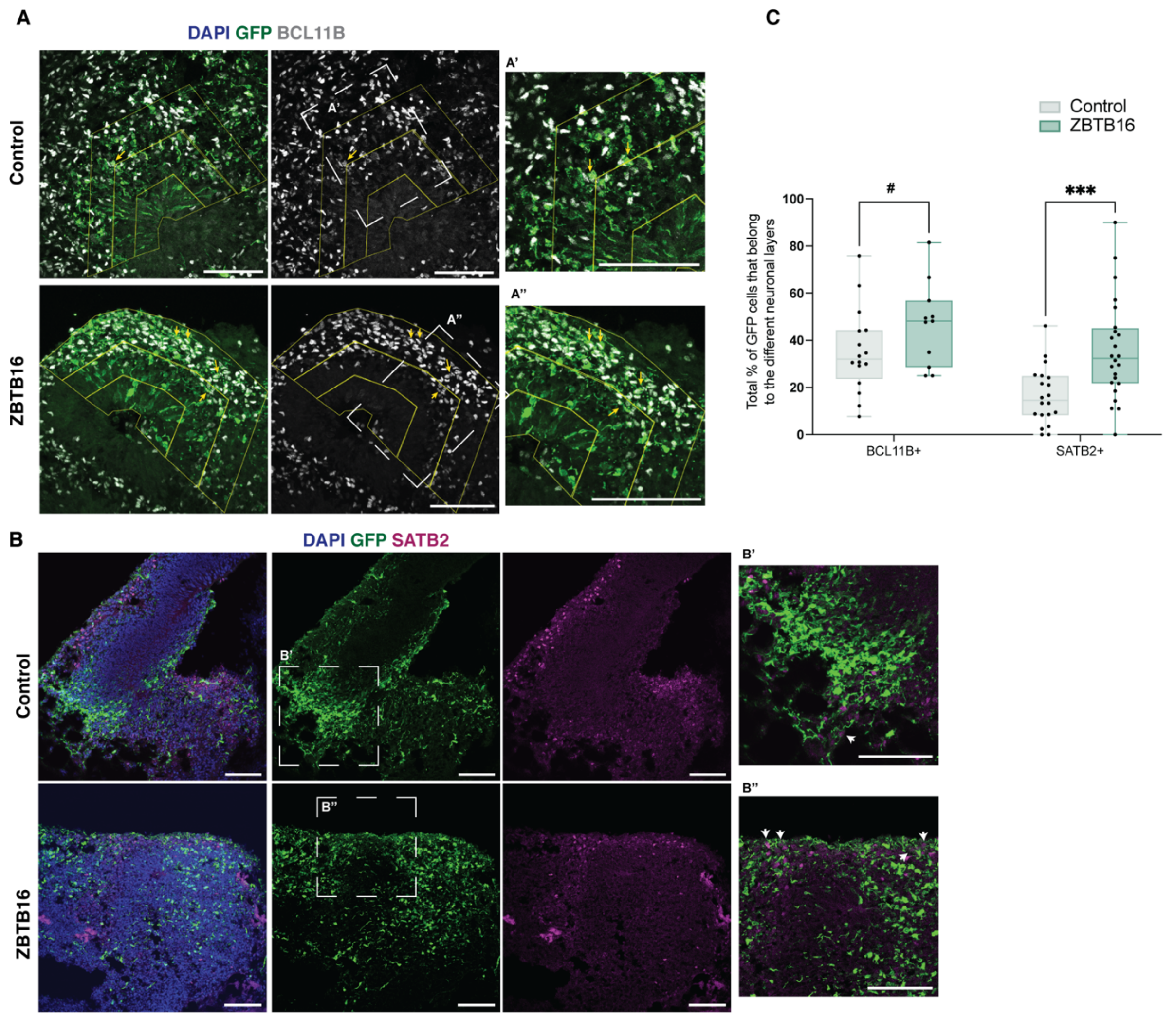


**Figure S7I Effects of ZBTB16 overexpression on neurons in human cerebral organoids. A,** Representative images of day 50 hCOs at control and ZBTB16 OE conditions stained for the layer V neuronal marker BCL11B, GFP and DAPI. Scale bars, 100µm. **B,** Representative images of day 50 hCOs at control and ZBTB16 OE conditions stained for the layer IV neuronal marker SATB2, GFP and DAPI. Scale bars, 100µm. **C,** Quantification of the total number of GFP cells that are BCL11B positive or SATB2 positive normalized by total GFP cells. Significance was tested with two-way ANOVA with Benjamini, Krieger and Yekutieli multiple testing correction (p.interaction= 0.36, p.bin= 0.0011, p.plasmid= 0.0009, F.interaction= 0.82, F.bin= 11.56, F.plasmid= 12.20, DF.interaction= 1, DF.bin = 1, DF.plasmid= 1). Post-hoc p-values: *** <=0.001, # = 0.056

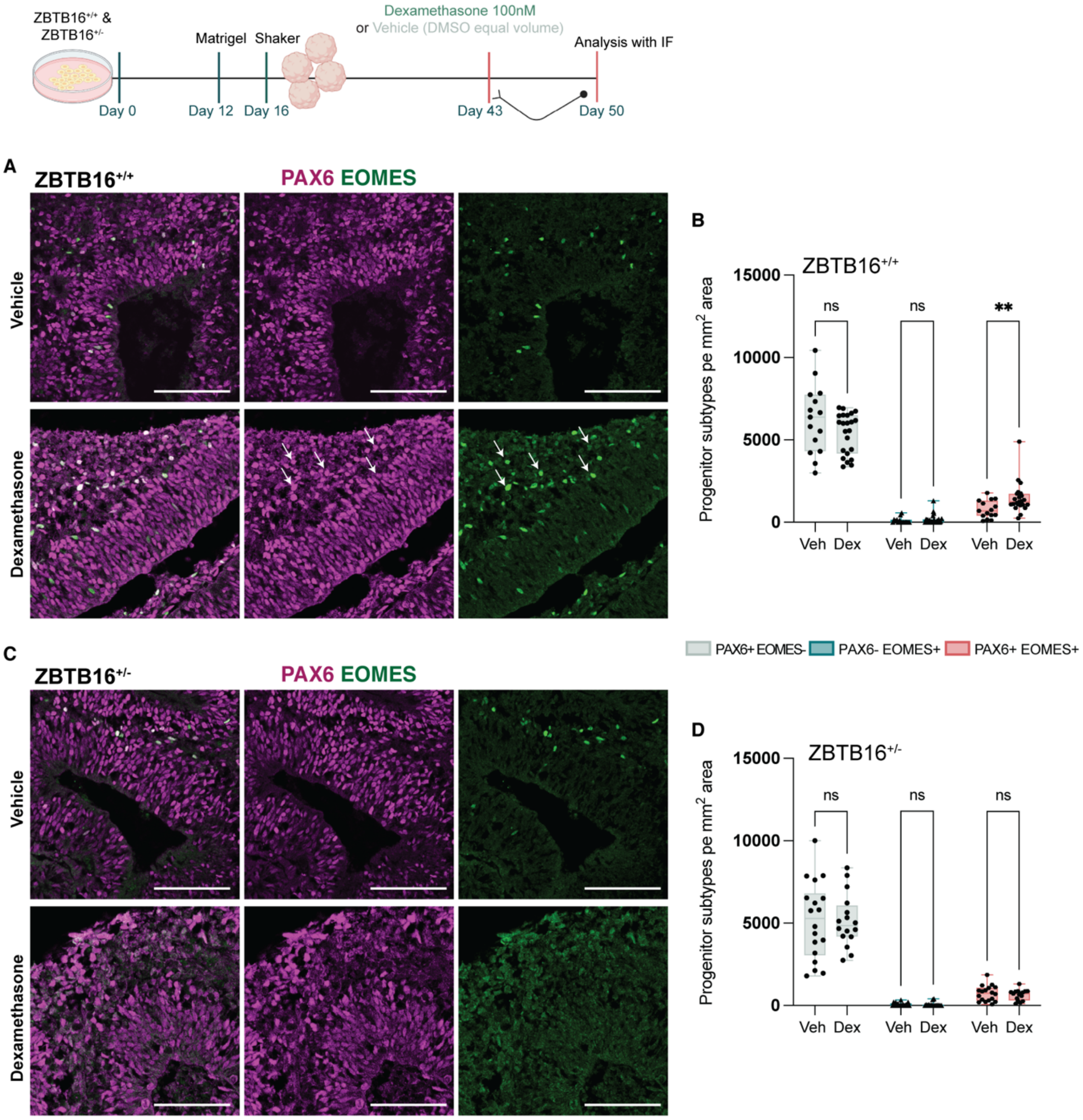


**Figure S8l Effects of dexamethasone treatment in progenitors' populations in wild-type and ZBTB16 heterozygous knock-out organoids. A,** Representative images of day 50 ZBTB16$^{+/+}$ hCOs at vehicle and dexamethasone treatment stained for PAX6 and EOMES. Scale bars, 100µm. **B,** Quantification of progenitors' populations shown as density of cells of square mm in wild-type hCOs. **C,** Representative images of day 50 ZBTB16 ZBTB16$^{+/-}$ hCOs at vehicle and dexamethasone treatment stained for PAX6 and EOMES. Scale bars, 100µm. **D,** Quantification of progenitors' populations shown as density of cells of square mm in ZBTB16 heterozygous KO hCOs. IF, immunofluorescence.

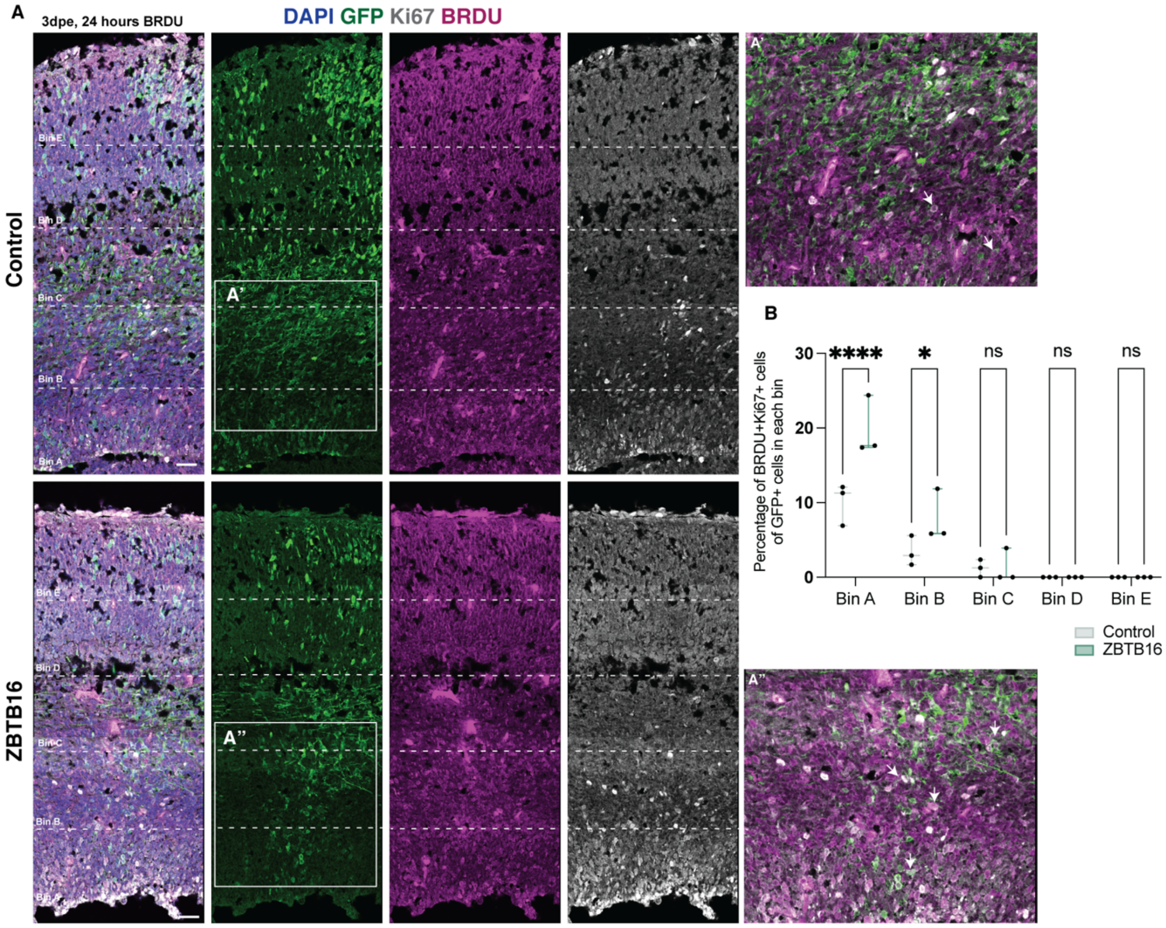


**Figure S9| Self-renewing capacity of cells electroporated with ZBTB16. A,** Representative images of E13.5 to E16.5 in utero electroporated mouse cortices that received a BRDU injection 24 hours before sacrifice stained for BRDU, GFP, Ki67 and DAPI. Scale bars, 100µm. **A' & A",** Zoom-ins of GFP+ cells. **B,** Quantification of GFP+ cells that co-stain for BRDU and Ki67 per bin normalised for total GFP+ cells in each bin. Quantification was done with a Two-way ANOVA with Benjamini, Krieger and Yekutieli multiple testing correction (p.interaction= 0.0024, F.interaction= 6, DF.interaction= 4). Post-hoc p-values: **** <=0.0001, * = 0.05

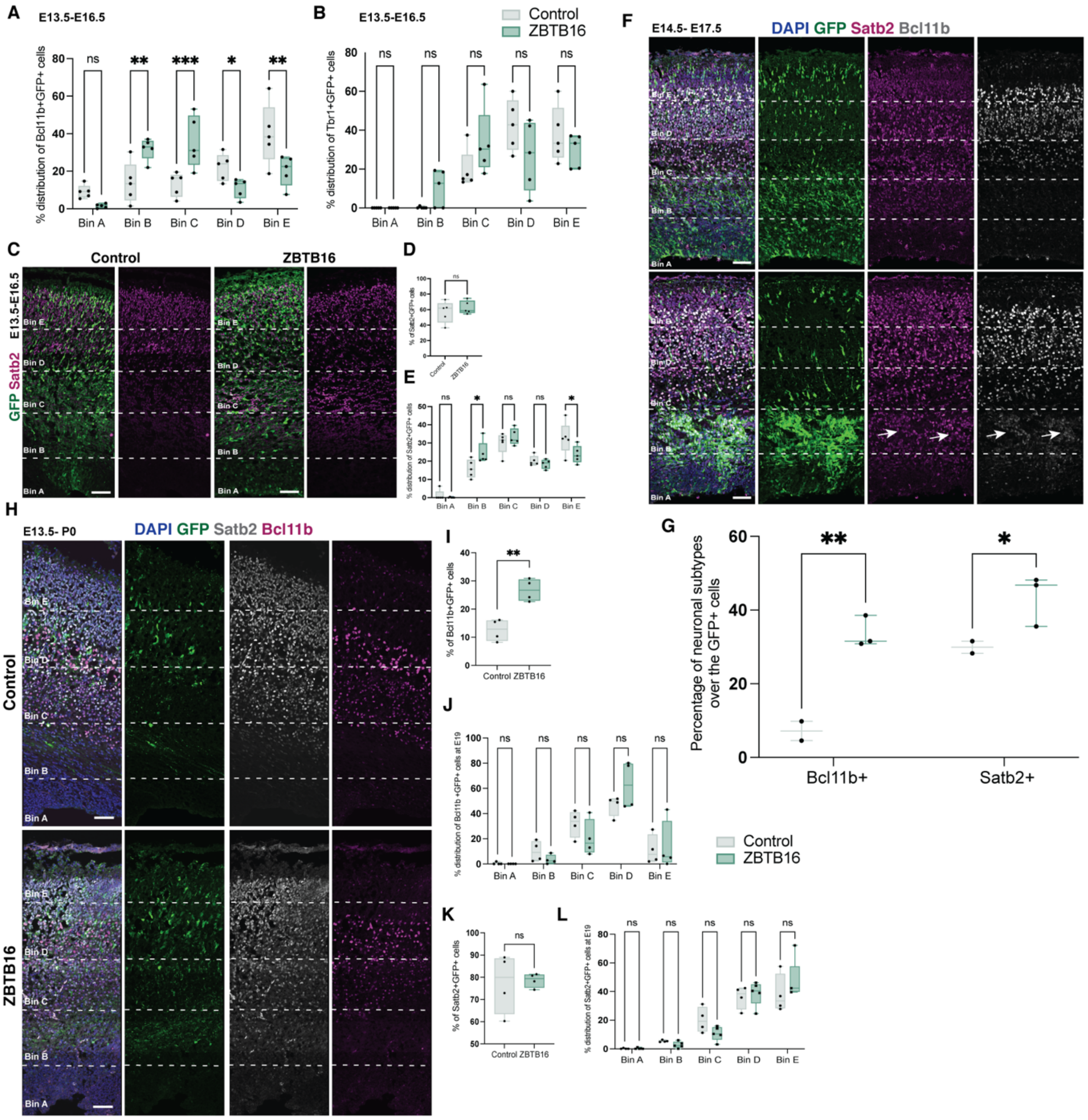


**Figure S10| Effects of ZBTB16 overexpression on neuronal subtypes in fetal mouse cortex. A,** Distribution of Bcl11b+GFP+ cells across bins. **B,** Distribution of Tbr1+GFP+ cells across bins. **C,** Representative images of fetal mouse cortices electroporated at E13.5 and analyzed at E16.5 at control and ZBTB16 OE conditions stained for GFP, the layer IV neuronal marker Satb2 and DAPI. Scale bars, 50µm. **D,** Quantification of the total number of GFP cells that are Satb2 positive normalized by total GFP cells in fetal mouse cortices electroporated at E13.5. **E,** Distribution of Satb2+GFP+ cells across bins. **F,** Representative images of fetal mouse cortices electroporated at E14.5 and analysed at E17.5 at control and ZBTB16 OE conditions stained for GFP, the layer V neuronal marker Bcl11b, the layer IV neuronal marker Satb2 and DAPI. Scale bars, 50µm. **G,** Quantification of the total number of GFP cells that are Satb2 positive and that are Bcl11b positive normalized by total GFP cells in fetal mouse cortices electroporated at E14.5. **H,** Representative images of fetal mouse cortices electroporated at E13.5 and analysed at E19.5/P0 at control and ZBTB16 OE conditions stained for GFP, the layer V neuronal marker Bcl11b, the layer IV neuronal marker Satb2 and DAPI. Scale bars, 50µm. **I,** Quantification of the total number of GFP cells that are Bcl11b+ normalized by total GFP cells in fetal mouse cortices. **J,** Distribution of Bcl11b+GFP+ cells per bin. **K,** Quantification of the total number of GFP cells that are Satb2+ normalized by total GFP cells in fetal mouse cortices. **L,** Distribution of Satb2+GFP+ cells. E16.5, embryonic day 16.5; qPCR, quantitative polymerase chain reaction;

Dex, dexamethasone; E13.5, embryonic day 13.5; P, postnatal. For D,I&K significance was tested with Mann-Whitney comparison between ZBTB16 overexpression and control plasmid. For A,B,E,G,J&L significance was tested with two-way ANOVA with Benjamini, Krieger and Yekutieli multiple testing correction (A, p.interaction< 0.0001, F.interaction= 10.66, DF.interaction=4/ B, p.interaction= 0.029, F.interaction= 2.99, DF.interaction=4/ E, p.interaction= 0.004, F.interaction= 4.39, DF.interaction=4/ G, p.plasmid= 0.0008, F.plasmid= 38.31, DF.plasmid= 1/ J, p.bin< 0.0001, F.interaction= 27.50, DF.interaction=4/ L, p.bin< 0.0001, F.interaction= 53.69, DF.interaction=4). Mann-Whitney p-values for D,I&K or post-hoc p-values for A,B,E,G&L: *** <=0.001, ** <=0.01, * <=0.05, ns >0.05

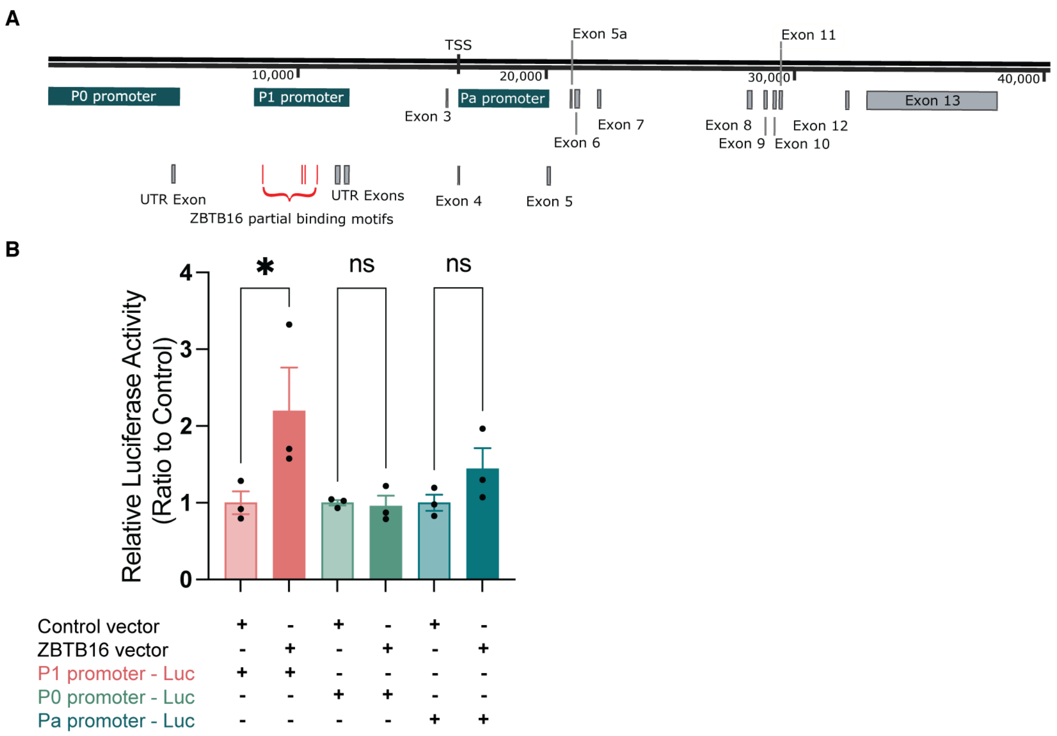


**Figure S11I *ZBTB16* activates the *PAX6* P1 promoter. A,** Schematic of the human *PAX6* locus. **B,** Quantification of the luciferase reporter assay results per promoter region and vector. Results are normalized to control transfections. Significance was tested with one-way ANOVA with Benjamini, Krieger and Yekutieli multiple testing correction (p= 0.04, F= 3.3, DF= 5). Post-hoc p-values: * <=0.05, ns >0.05

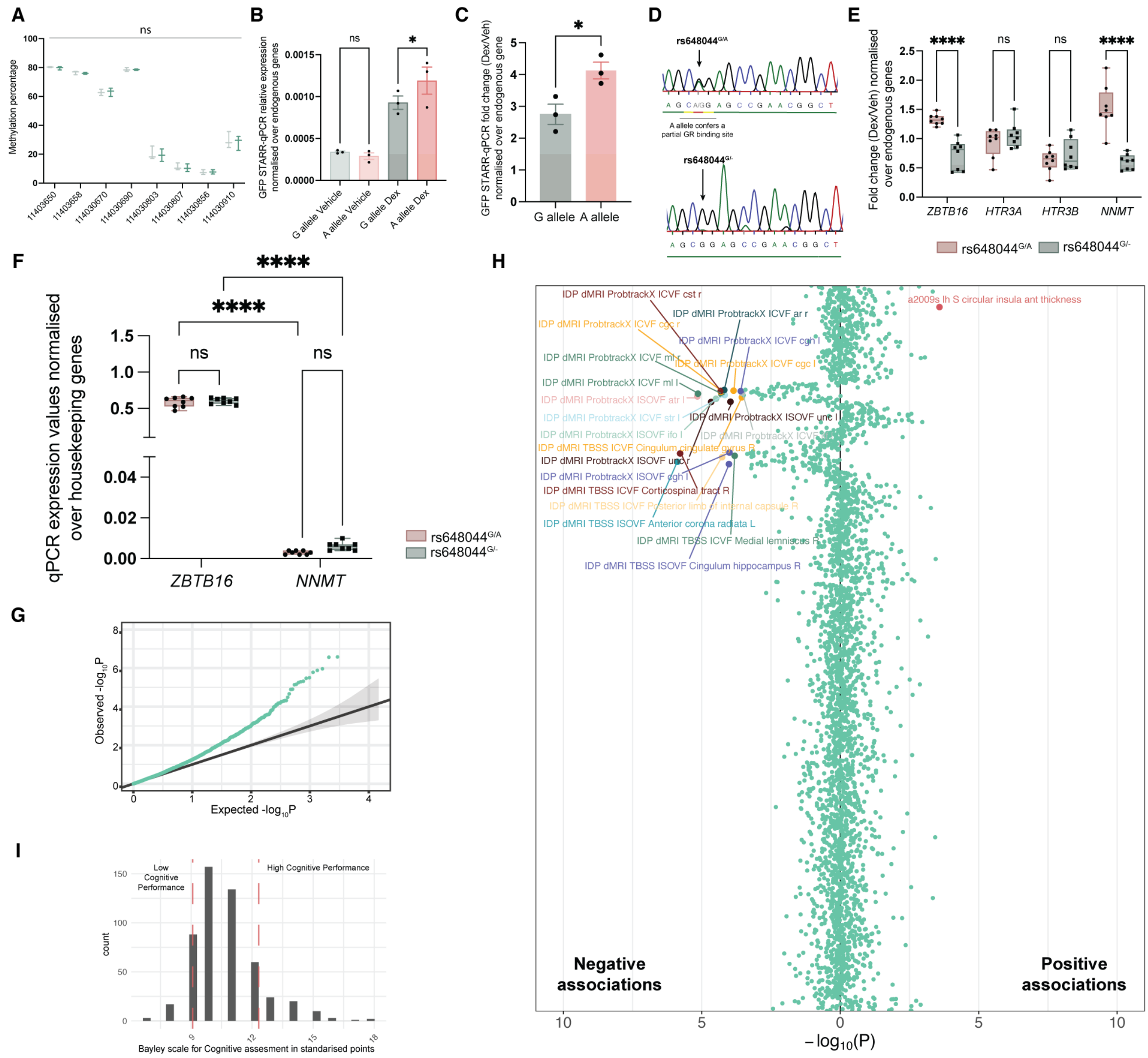


**Figure S12| rs648044 effects on *ZBTB16* transcription and postnatal phenotypes**. **A,** Methylation levels of CpGs of vehicle and dexamethasone treated hCOs in the genetic locus surrounding rs648044 as found with Pyrosequencing. **B,** STARR-qPCR results of the effect of the two alleles of rs648044 on expression normalized over housekeeping genes in U20SGR cells treated with 100nM dex for 12hrs. **C,** STARR-qPCR results of the effect of the two alleles of rs648044 on expression shown as fold changes and normalized over housekeeping genes in GR18 cells treated with 100nM dex for 12hrs. **D,** Sanger sequencing traces for the ZBTB16 locus that includes rs648044, in rs648044$^{G/A}$ Line 2-iPSCs and rs648044$^{G/-}$ edited iPSCs. **E,** Fold changes of the expression of all genes located near rs648044 in rs648044$^{G/A}$ and rs648044$^{G/-}$ day 30 hCOs treated with 100nM dex for 7 days. **F,** qPCR normalized over housekeeping genes expression values of *ZBTB16* and *NNMT* in rs648044$^{G/A}$ and rs648044$^{G/-}$ day 30 hCOs at vehicle condition. Horizontal pleiotropy is not likely within the locus as rs648044 relevantly regulates cis glucocorticoid-effects on brain tissue transcription only for ZBTB16, as *NNMT* is barely detectable in the adult and fetal brain[6] and in the hCOs (see figure), while it is very abundant in the liver. **G,** Quantile-quantile plot of expected versus observed mendelian randomization p-values for the effects of the rs648044 x glucocorticoids predicted ZBTB16 transcription on UK BioBank phenotypes. **H,** Plot describing associations of *ZBTB16* transcription on neuroimaging traits described in Elliot *et al.*, Nature, 2018 (see Table S15). Association with imaging phenotypes are presented based on negative (negative MRa estimate – lower quantitative measures with A-allele effects) and positive (positive MRa estimate - higher quantitative measures with A-allele effects) effects. Traits that remain significant following Benjamini-Hochberg correction are labelled with their variable name.

**I,** Histogram of the Bayley cognitive scale (standardized points) across the ITU cohort. Low cognitive performance was defined as 1 SD below the sample mean whereas high cognitive performance as 1 SD above the sample mean. MRa, mendelian randomization; qPCR, quantitative polymerase chain reaction; STARR, self-transcribing active regulatory region; hCOs, human cerebral organoids; dex, dexamethasone, ICVF, intracellular volume fraction; ISOVF, isotropic volume fraction; TBSS, tract based spatial statistics; IDP, image derived phenotypes; ITU, intra-uterine. For B significance was tested with Mann-Whitney comparison between ZBTB16 overexpression and control plasmid. For B,E&F significance was tested with two-way ANOVA with Benjamini, Krieger and Yekutieli multiple testing correction (B: $p= 0.12$, $F= 2.85$, $DF= 1$/ E: $p< 0.0001$, $F= 20$, $DF= 3$/ F: $p= 0.61$, $F= 0.25$, $DF= 1$). Mann-Whitney p-value for C or post-hoc p-values for B,E&F: **** <=0.0001, * <=0.05, ns >0.05


1. Betizeau, M., Cortay, V., Patti, D., Pfister, S., Gautier, E., Bellemin-Ménard, A., Afanassieff, M., Huissoud, C., Douglas, R.J., Kennedy, H., et al. (2013). Precursor Diversity and Complexity of Lineage Relationships in the Outer Subventricular Zone of the Primate. Neuron *80*, 442–457. 10.1016/j.neuron.2013.09.032.
2. Pebworth, M.P., Ross, J., Andrews, M., Bhaduri, A., and Kriegstein, A.R. (2021). Human intermediate progenitor diversity during cortical development. Proc Natl Acad Sci U S A *118*, 1–10. 10.1073/pnas.2019415118.
3. Garcia, M.T., Chang, Y., Arai, Y., and Huttner, W.B. (2016). S-Phase Duration Is the Main Target of Cell Cycle Regulation in Neural Progenitors of Developing Ferret Neocortex. J Comp Neurol *470*, 456–470. 10.1002/cne.23801.
4. Prasad, R., Jung, H., Tan, A., Song, Y., Moon, S., Shaker, M.R., Sun, W., Lee, J., Ryu, H., Lim, H.K., et al. (2021). Hypermethylation of Mest promoter causes aberrant Wnt signaling in patients with Alzheimer's disease. Sci Rep *11*, 1–10. 10.1038/s41598-021-99562-9.
5. Zhang, M., Haughey, M., Wang, N.Y., Blease, K., Kapoun, A.M., Couto, S., Belka, I., Hoey, T., Groza, M., Hartke, J., et al. (2020). Targeting the Wnt signaling pathway through R-spondin 3 identifies an anti-fibrosis treatment strategy for multiple organs. PLoS One *15*, 1–21. 10.1371/journal.pone.0229445.
6. Uhlén, M., Fagerberg, L., Hallström, B.M., Lindskog, C., Oksvold, P., Mardinoglu, A., Sivertsson, Å., Kampf, C., Sjöstedt, E., Asplund, A., et al. (2015). Tissue-based map of the human proteome. Science (1979) *347*. 10.1126/science.1260419.
